# Genetic nurture and shared environmental effects across cognitive, educational, and mental health outcomes

**DOI:** 10.64898/2026.11.16.767077

**Authors:** Yujing Lin, Ziada Ayorech, Kaito Kawakami, Engin Keser, Thalia C. Eley, Kaili Rimfeld, Margherita Malanchini, Robert Plomin

**Affiliations:** Social, Genetic and Developmental Psychiatry Centre, King’s College London, UK; PROMENTA Research Center, Department of Psychology, University of Oslo, Oslo 0373, Norway; Department of Child & Adolescent Psychiatry, Institute of Psychiatry, Psychology & Neuroscience, King’s College London, UK; National Institute for Health and Care Research (NIHR) Maudsley Biomedical Research Centre (BRC), South London and Maudsley NHS Trust, London, UK; Department of Psychology, Royal Holloway, University of London, UK; School of Biological and Behavioural Sciences, Queen Mary University of London, UK

## Abstract

Genetic nurture, in which parental genotype predicts offspring outcomes independent of offspring genotype, operates at the between-family level, corresponding to the shared environmental effect (C) in the classical twin design. Genetic nurture represents a portion of C captured by parental genotype.

The link between genetic nurture and C has not been systematically quantified across multiple trait domains. Using 2559 families with genotyped parents and 6710 twin pairs, we estimated genetic nurture using polygenic scores (PGS) and C using the twin design for 40 educational, cognitive, and anthropometric outcomes from age 7 to 26, and for 8 psychopathology symptoms at age 26.

Although genetic nurture effects were small, the largest effects were observed for educational outcomes, followed by cognitive abilities, and then psychopathology symptoms. C followed the same pattern. Across eight trait domains, the correlation between genetic nurture and C was 0.68.

However, psychopathology outcomes also showed relatively low PGS prediction, which creates a ceiling for estimated genetic nurture. In proportion to PGS prediction, psychopathology showed as much genetic nurture as educational outcomes. Why does psychopathology show genetic nurture despite showing no C? Why is psychopathology PGS prediction so low? We suggest that nonadditive genetic variance could explain both.

## Introduction

Familial resemblance can arise from shared genes and shared environment; disentangling the two has been the central issue for quantitative genetic research ^1^. By comparing monozygotic (MZ) twins, who are identical for inherited DNA variants, to dizygotic (DZ) twins, whose genetic relatedness is on average 50% for additive genetic effects, the twin design can estimate the contribution of genetic influences: the A, additive genetic variance, in the ACE model. Shared environmental influence (C, common) is the residual resemblance of MZ and DZ twins not explained by their genetic relatedness. A and C are the quantitative genetic representations of the century-old battleground of nature and nurture.

Molecular genetics offers a novel perspective on this issue by enabling, for example, the comparison of parents’ and children’s genotypes. Parents pass on half of their genome; the other half can still shape children’s outcomes through the environment it creates: a phenomenon termed ‘genetic nurture’ ^2,3^. Genetic nurture is estimated as the association between parental genotype and offspring outcome independent of the offspring genotype. It captures a portion of C influences that can be tagged by parents’ measured genotypes, circumventing the need for assessment of the salient environments. Genetic nurture also contributes to polygenic score (PGS) prediction in the population. Because children are genetically related to their parents, the association between the child’s PGS and phenotypic outcome captures both an ‘indirect’ component due to genetic nurture and a ‘direct’ component ^3,4^.

Genetic nurture has been widely reported for educational attainment (i.e., years of schooling) and achievement ^2–9^. For example, a meta-analysis reported a pooled beta of 0.08 for educational outcomes ^10^. Less is known about cognitive abilities ^11,12^. Evidence for psychiatric traits is mixed, often depending on the predictor trait or outcome examined and the developmental stage. For substance use and internalising symptoms, genetic nurture has been detected, though not consistently ^13–21^. Genetic nurture is largely absent for externalising symptoms ^22–25^ and most neurodevelopmental traits ^26–31^. Height and body mass index (BMI) are typically treated as negative controls ^9^, although significant maternal genetic nurture for height has been reported in one study ^3^. Trio genome-wide complex trait analysis (GCTA), relatedness disequilibrium regression, and family-based genome-wide association analysis have also been used to estimate genetic nurture ^27,32–36^, but our focus is on PGS because they offer the most power to detect genetic nurture given a well-powered GWAS while variance decomposition methods require larger samples ^35^.

Here we test empirically a proposed link between genetic nurture and shared environmental influence (C). Both genetic nurture and C are between-family environmentally mediated effects that increase parent–offspring resemblance. Genetic nurture is much narrower, representing a portion of C that is tagged by parental genotypes independent of children’s genotypes, whereas C captures all between-family environmental effects that make siblings in the same family resemble each other, as estimated in twin designs. Thus, genetic nurture should mainly be found for traits that show C although C could be found for traits that show no genetic nurture.

General support for an empirical link between genetic nurture and C comes from the finding that educational attainment, which shows the most consistent evidence for genetic nurture, also shows the largest estimates of C, about 30% ^37^. At the other extreme, psychopathology, for which genetic nurture has seldom been reported, shows much less evidence of C ^1^. However, PGS predictive power creates a ceiling for identifying genetic nurture and that ceiling is over twice as high for educational attainment as for psychopathology ^38^, suggesting an alternative explanation for finding more genetic nurture for educational attainment than for psychopathology.

The correspondence between genetic nurture and C has not been systematically quantified across trait domains in a single sample, to our knowledge. We addressed this using data from the Twins Early Development Study (TEDS) ^39^, a UK-based, population-representative longitudinal twin cohort with genotyped parents (2460 mother–child duos and 1045 father–child duos, including 946 unrelated complete trios, and 6710 twin pairs). We investigated genetic nurture and C for 48 outcomes across diverse trait domains: educational achievement and attainment, cognitive abilities, height and BMI from age 7 to 26, as well as psychopathology symptoms (e.g., depression, eating disorders and alcohol use) at age 26. We were therefore able to estimate genetic nurture and C in the same participants at the same ages with the same measures. This allowed a clean empirical test of whether genetic nurture, a between-family environmental effect captured by parental genotype, is correlated with shared environmental influence (C).

## Results

We combined two family designs within the same sample: a trio polygenic score design to estimate genetic nurture and a classical twin design to estimate shared environmental influence, C. We first present genetic nurture estimates across trait domains and then compare these estimates to estimates of C. We also investigate other models of genetic nurture and the contribution of socio-economic status.

### Overview of the trio design and parental genotype data

We used a trio design to estimate genetic nurture and child direct genetic effects (Figure 1). In a regression model, child, mother, and father polygenic scores jointly predicted the child’s outcome, with child age, sex and genotyping chip as covariates. All PGS were adjusted for ancestry (i.e., the first ten genomic principal components) before entering the model. The child coefficient indexes the child direct genetic effect, capturing the genetic effects attributed to the child’s own PGS independent of maternal and paternal genetic nurture. The maternal and paternal PGS coefficients index their association with the child’s outcome independent of the child’s PGS, representing maternal and paternal genetic nurture, respectively.

**Figure 1.**
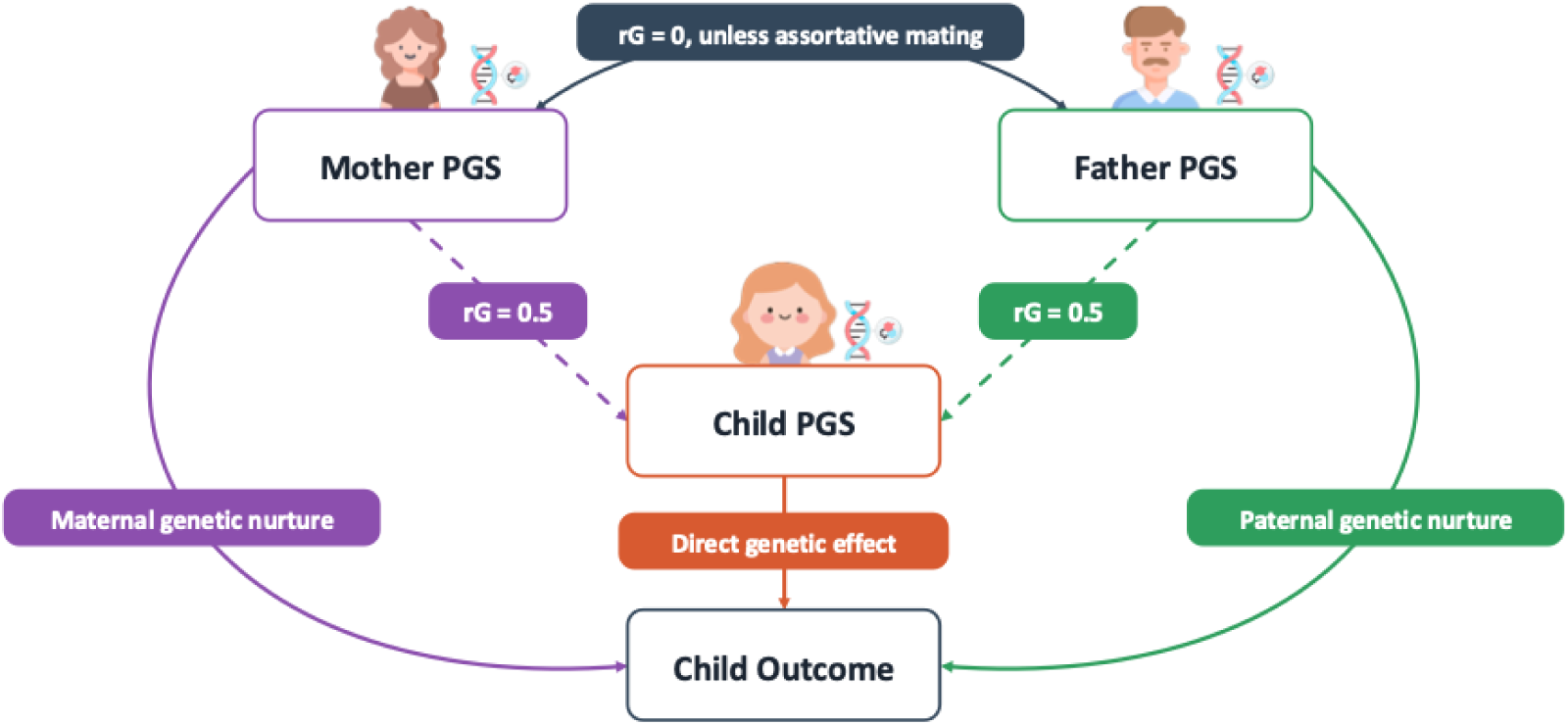
Parent–child trio design using polygenic scores. Child, maternal and paternal polygenic scores for the same trait jointly predict the child’s outcome. The path from the child’s PGS independent of parental PGS indexes the child direct genetic effect; the paths from the maternal and paternal PGS independent of the child’s PGS index maternal and paternal genetic nurture. Dashed arrows denote genetic transmission, with an expected parent–offspring PGS correlation of 0.5 in the absence of assortative mating. The double-headed arrow between parents denotes their PGS correlation, which should approximate zero in the absence of assortative mating. PGS = polygenic score. rG = correlation between polygenic scores. Drawings obtained from https://www.flaticon.com.

These coefficients all contribute to the population PGS prediction, that is, the correlation between the child’s PGS and the phenotypic outcome. The child direct genetic effect is included in the population prediction in full, whereas the parental PGS coefficients enter through parent–child PGS correlations (dashed arrows in Figure 1). The population prediction can therefore be expressed as the child direct genetic effect plus, for each parent, the product of the parental PGS coefficient and the corresponding parent–child PGS correlation (Table 1).

**Table 1.**
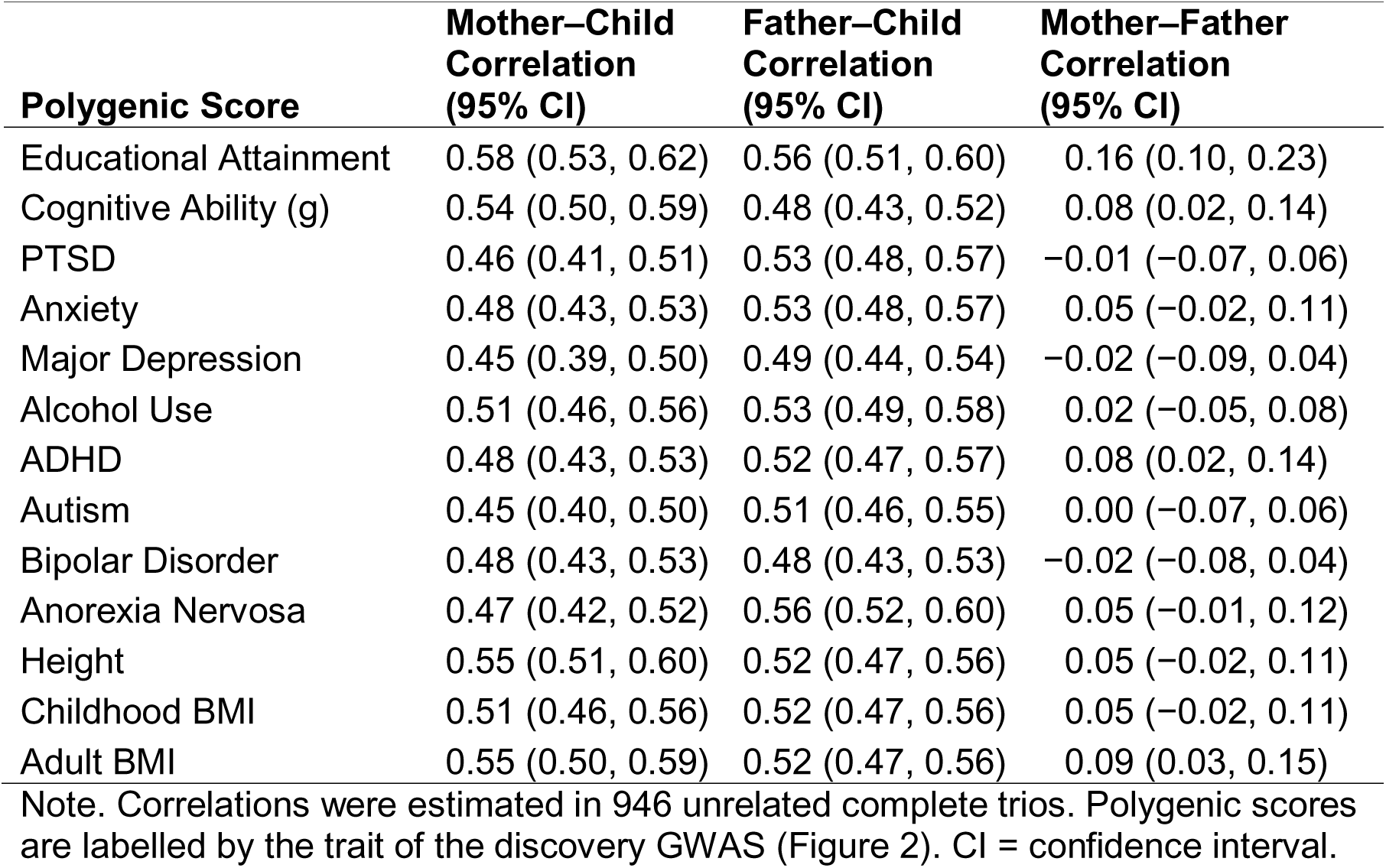
Polygenic score correlations within families.

| **Polygenic Score** | **Mother–Child Correlation (95% CI)** | **Father–Child Correlation (95% CI)** | **Mother–Father Correlation (95% CI)** |
|---|---|---|---|
| Educational Attainment | 0.58 (0.53, 0.62) | 0.56 (0.51, 0.60) | 0.16 (0.10, 0.23) |
| Cognitive Ability (g) | 0.54 (0.50, 0.59) | 0.48 (0.43, 0.52) | 0.08 (0.02, 0.14) |
| PTSD | 0.46 (0.41, 0.51) | 0.53 (0.48, 0.57) | −0.01 (−0.07, 0.06) |
| Anxiety | 0.48 (0.43, 0.53) | 0.53 (0.48, 0.57) | 0.05 (−0.02, 0.11) |
| Major Depression | 0.45 (0.39, 0.50) | 0.49 (0.44, 0.54) | −0.02 (−0.09, 0.04) |
| Alcohol Use | 0.51 (0.46, 0.56) | 0.53 (0.49, 0.58) | 0.02 (−0.05, 0.08) |
| ADHD | 0.48 (0.43, 0.53) | 0.52 (0.47, 0.57) | 0.08 (0.02, 0.14) |
| Autism | 0.45 (0.40, 0.50) | 0.51 (0.46, 0.55) | 0.00 (−0.07, 0.06) |
| Bipolar Disorder | 0.48 (0.43, 0.53) | 0.48 (0.43, 0.53) | −0.02 (−0.08, 0.04) |
| Anorexia Nervosa | 0.47 (0.42, 0.52) | 0.56 (0.52, 0.60) | 0.05 (−0.01, 0.12) |
| Height | 0.55 (0.51, 0.60) | 0.52 (0.47, 0.56) | 0.05 (−0.02, 0.11) |
| Childhood BMI | 0.51 (0.46, 0.56) | 0.52 (0.47, 0.56) | 0.05 (−0.02, 0.11) |
| Adult BMI | 0.55 (0.50, 0.59) | 0.52 (0.47, 0.56) | 0.09 (0.03, 0.15) |

Note. Correlations were estimated in 946 unrelated complete trios. Polygenic scores are labelled by the trait of the discovery GWAS (Figure 2). CI = confidence interval.

We first estimate genetic nurture as the parental PGS coefficients from the trio model. We then examine its contribution to the population PGS prediction.

We applied the trio model to the TEDS families. DNA newly collected for the parents of TEDS twins yielded 946 unrelated complete trios (54% females) after quality control (Supplementary Method 1 and Supplementary Figure 1). In total, 2559 unrelated children (20% of the 13,093 unrelated children in TEDS) had at least one genotyped parent (56% females): 2460 with maternal and 1045 with paternal DNA. When available, we included both members of dizygotic twins to increase statistical power. Because twins from the same family are not independent observations, we used standard errors that account for clustering within families ^40,41^. Sibling ‘genetic nurture’ effects are addressed later.

As shown in Table 1 (details in Supplementary Table 1), correlations between 13 parent and offspring PGS were consistent with population genetic expectations given some assortative mating. The average parent–child PGS correlations were 0.50 for mothers (range 0.45 to 0.58) and 0.52 for fathers (range 0.48 to 0.56) across the 13 PGS. PGS assortative mating can be seen in mother–father PGS correlations greater than zero, especially for the educational attainment PGS (r = 0.16), for which phenotypic assortative mating is greatest (r = 0.55) ^42^. Because maternal, paternal and child PGS enter the trio model simultaneously, assortative mating indexed by the mother–father PGS correlation is accounted for in our estimates of genetic nurture.

As shown in Figure 2, and in line with the core aims of the study, we focused on isomorphic comparisons between PGS and traits – for example, the cognitive ability PGS with cognitive abilities and the educational attainment PGS with educational achievement and attainment –rather than cross-trait or multi-trait analyses.

**Figure 2.**
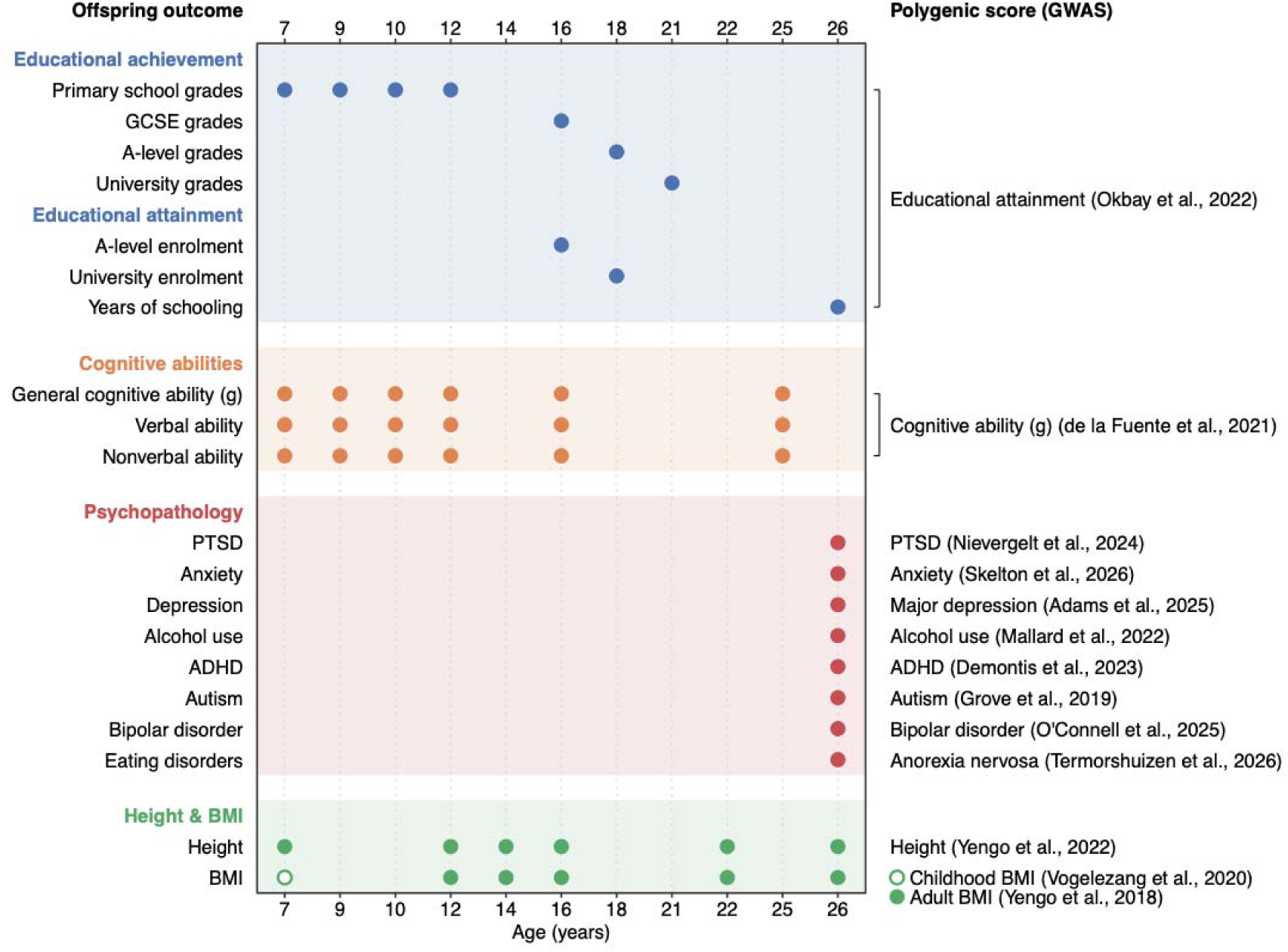
Overview of phenotypic outcomes and polygenic score predictors. Each row is an offspring outcome, grouped into educational achievement, educational attainment, cognitive abilities, psychopathology, height and BMI. The x-axis marks the age at which the outcome was assessed. Cognitive, educational and anthropometric outcomes were assessed from age 7 to 26, and psychopathology symptoms at age 26. The right-hand column lists the polygenic score used to predict each outcome, with its discovery GWAS in parentheses; brackets indicate a single polygenic score predicting several outcomes. Cognitive abilities were predicted by the same cognitive PGS at all ages because childhood-specific GWAS of intelligence have small sample sizes and genetic effects on intelligence are highly correlated across age groups ^43^. Consistent with this, PGS from adult GWAS predict cognitive abilities in childhood ^44^. BMI at age 7 (open circle) was predicted by a childhood BMI polygenic score, and BMI at later ages (filled circles) by an adult BMI polygenic score. Eating disorder symptoms were predicted by an anorexia nervosa polygenic score. For the analysis of each trait, the same PGS was used for child, mother, and father. For the presentation of the results in the text (with complete analyses in the supplementary material), estimates were averaged within eight trait domains: educational achievement, educational attainment, general cognitive ability, verbal ability, nonverbal ability, height, BMI and psychopathology. GWAS = genome-wide association study; GCSE = General Certificate of Secondary Education; PTSD = post-traumatic stress disorder; ADHD = attention-deficit/hyperactivity disorder; BMI = body mass index.

In the following sections, we present the results from the trio analyses for each domain and then compare these results to twin estimates of C.

### Genetic nurture

Table 2 lists the child direct genetic effects and parental genetic nurture from the trio regressions. Figure 3 expresses these effects as contributions to the population PGS prediction. These analyses are based on complete trios: up to 893 families, or 1356 participants counting both dizygotic twins (Supplementary Table 2).

**Figure 3.**
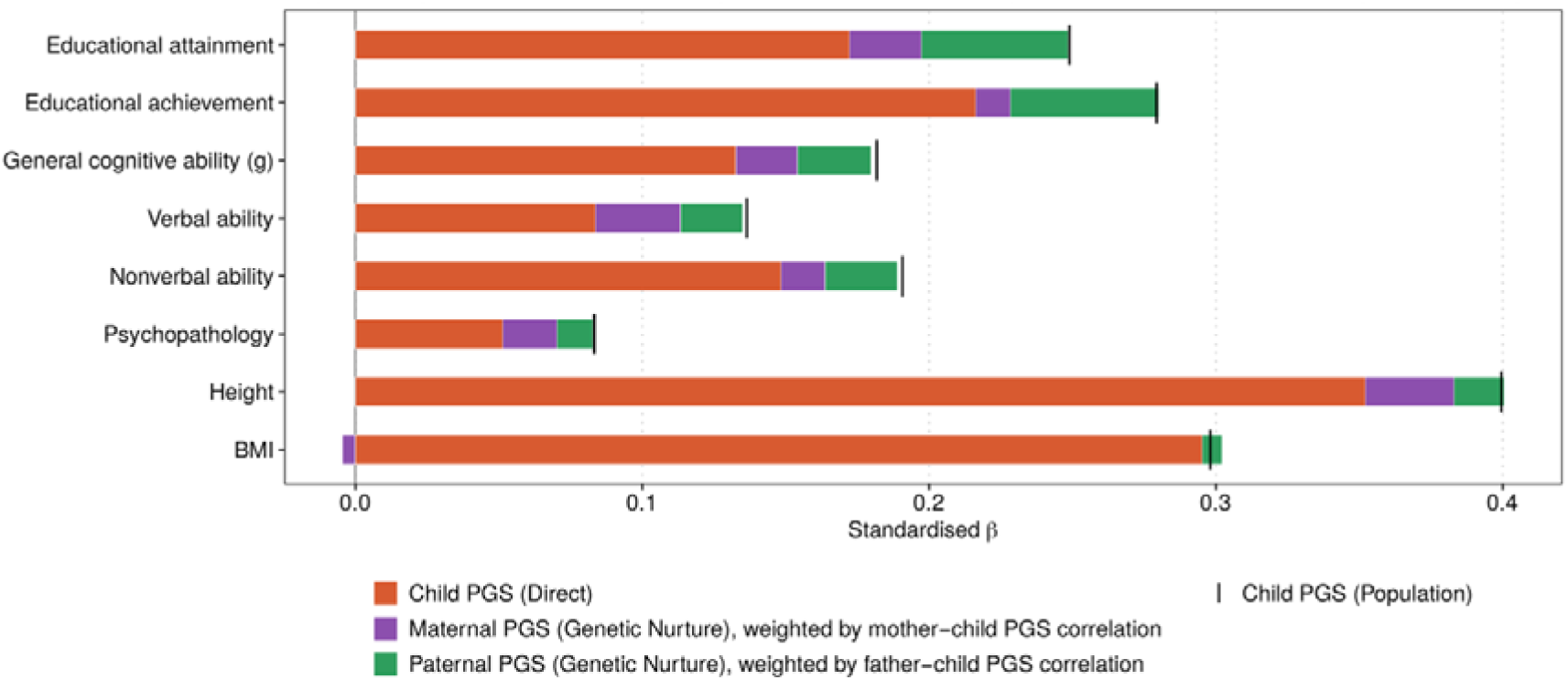
Effect sizes for child direct genetic effects and genetic nurture within the population PGS prediction, averaged within trait domains. The black vertical line at the end of each bar represents the population prediction (as in Table 2), that is, the correlation (standardised regression) between the child’s PGS and the child’s phenotypic outcome. The population prediction can be expressed as the sum of child direct genetic effects, maternal genetic nurture, and paternal genetic nurture, where maternal and paternal genetic nurture are the parental PGS coefficients (Table 2) multiplied by the corresponding parent–child PGS correlations in Table 1. The child direct genetic effect enters this sum unchanged (as in Table 2). The small gaps between the population prediction and the sum of its three components arise because these correlations were estimated in unrelated complete trios, whereas the regression models were fitted for each outcome’s analysis sample, which also includes dizygotic co-twins. Because the bars are stacked, confidence intervals are not plotted here. Supplementary Figure 3 shows the corresponding figure for all 48 outcomes. Estimates with 95% confidence intervals and statistical significance are presented in Supplementary Tables 2 and 3.

**Table 2.** Parental genetic nurture, child direct genetic effect, and population PGS effect.

| Trait Domain | Maternal Genetic Nurture (95% CI) | Paternal Genetic Nurture (95% CI) | Direct Genetic Effect (95% CI) | Population Prediction (95% CI) |
|---|---|---|---|---|
| Educational Attainment | 0.04<br>(−0.01, 0.09) | 0.09<br>(0.04, 0.14) | 0.17<br>(0.12, 0.23) | 0.25<br>(0.21, 0.29) |
| Educational Achievement | 0.02<br>(−0.04, 0.08) | 0.09<br>(0.04, 0.14) | 0.22<br>(0.16, 0.28) | 0.28<br>(0.24, 0.32) |
| General Cognitive Ability (g) | 0.04<br>(−0.02, 0.10) | 0.05<br>(0.00, 0.11) | 0.13<br>(0.07, 0.19) | 0.18<br>(0.13, 0.23) |
| Verbal Ability | 0.05<br>(−0.00, 0.11) | 0.05<br>(−0.01, 0.10) | 0.08<br>(0.03, 0.14) | 0.14<br>(0.09, 0.18) |
| Nonverbal Ability | 0.03<br>(−0.02, 0.09) | 0.05<br>(−0.00, 0.10) | 0.15<br>(0.09, 0.21) | 0.19<br>(0.15, 0.24) |
| Psychopathology | 0.04<br>(0.01, 0.07) | 0.02<br>(−0.01, 0.05) | 0.05<br>(0.02, 0.09) | 0.08<br>(0.06, 0.11) |
| Height | 0.06<br>(0.01, 0.10) | 0.03<br>(−0.01, 0.08) | 0.35<br>(0.31, 0.40) | 0.40<br>(0.37, 0.43) |
| BMI | −0.01<br>(−0.06, 0.04) | 0.01<br>(−0.03, 0.06) | 0.30<br>(0.24, 0.35) | 0.30<br>(0.26, 0.34) |

Note. Standardised regression coefficients from complete trios, averaged across the outcomes in each domain, after accounting for covariates. Maternal genetic nurture, paternal genetic nurture and child direct genetic effect correspond to the mother’s, father’s and child’s PGS coefficients in the trio regression model. Population prediction is the correlation (standardised regression) between the child’s PGS and the phenotypic outcome. The 95% confidence intervals for the domain averages are percentile intervals from 1000 cluster bootstrap resamples of families. Results including standardised regression coefficients and 95% confidence intervals for all 48 outcomes are plotted in Supplementary Figure 2 and presented in Supplementary

We focus on average effect sizes aggregated across traits within a domain rather than point estimates for all 48 outcomes (Figure 2) for three reasons. First, our central question concerns the correspondence between genetic nurture and C across trait domains. Second, we find that effect sizes of genetic nurture for each parent are very small, ranging from −0.05 to 0.15, accounting for less than 2% of the outcome variance. Power analyses indicate that 1000 trios are needed for 80% power to detect genetic nurture accounting for 1.5% of the outcome variance per parent ^35^. As a result, the confidence intervals for our point estimates for the 48 variables are wide (Supplementary Figure 2), and averaging results within a domain better represents the domain. The third reason is presentational: to portray the general landscape of genetic nurture across domains.

For these reasons, Table 2 and Figure 3 present average effect sizes for eight domains. Results for all 48 variables for each parent are listed in Supplementary Figures 2 and 3. Numerical estimates and confidence intervals are presented in Supplementary Tables 2 and 3.

### Educational attainment and achievement

For educational attainment, as shown in Table 2, maternal and paternal genetic nurture were 0.04 and 0.09, which are the coefficients of the mother’s and father’s PGS predicting the child’s educational attainment independent of the child’s own PGS in the trio regression model. The child direct genetic effect was 0.17.

At the population level, the child’s PGS correlated 0.25 with educational attainment. Figure 1 shows how Tables 1 and 2 can be combined to produce this population prediction. The population prediction can be expressed as the sum of the child direct genetic effect and genetic nurture, as summarised in Figure 3. Because the population prediction uses only the child’s PGS, parental genetic nurture can appear in it only to the extent that the child’s PGS reflects the parents’ PGS. The child’s PGS correlates about 0.50 with each parent’s PGS (Figure 1), so the population prediction picks up only about half of each parent’s genetic nurture, whereas the child direct genetic effect enters in full. For the educational attainment PGS, these correlations between parent and child PGS are 0.58 for mothers and 0.56 for fathers (Table 1), higher than the expected 0.50, consistent with assortative mating. Each parent’s contribution to the population prediction is therefore their genetic nurture (Table 2) multiplied by their PGS correlation with the child’s PGS (Table 1).

Thus, 29% of the population prediction for educational attainment can be attributed to genetic nurture (0.04 × 0.58 + 0.09 × 0.56) / 0.25, and 68% to the child direct genetic effect (0.17 / 0.25). For illustration, the percentages are calculated from the rounded estimates shown in Tables 1 and 2; Supplementary Table 3 reports the values calculated from unrounded estimates. The two percentages do not sum exactly to 100%, both because of rounding and because the parent–child PGS correlations were estimated in unrelated complete trios rather than in each outcome’s analysis sample (Figure 3).

For educational achievement, maternal and paternal genetic nurture were 0.02 and 0.09, and the child direct genetic effect was 0.22 (Table 2). The population prediction was 0.28. After applying the parent–child PGS correlations from Table 1 to the estimates from Table 2, 22% of the population prediction for educational achievement can be attributed to genetic nurture and 79% to the child direct genetic effect, as shown in Figure 3.

### Cognitive abilities

For general cognitive ability, maternal and paternal genetic nurture estimates were 0.04 and 0.05, respectively, and the child direct genetic effect was 0.13 (Table 2). The population prediction was 0.18. The correlations between parent and child PGS were 0.54 for mothers and 0.48 for fathers (Table 1). After applying these correlations, 25% of the population prediction for general cognitive ability can be attributed to genetic nurture and 72% to the child direct genetic effect.

For verbal and nonverbal ability, maternal genetic nurture was 0.05 for verbal ability and 0.03 for nonverbal ability and paternal genetic nurture was 0.05 for both. The child direct genetic effect was 0.08 for verbal and 0.15 for nonverbal. The population predictions were 0.14 for verbal and 0.19 for nonverbal. Applying the same parent–child PGS correlations, genetic nurture accounted for 36% of the population prediction for verbal ability and 21% for nonverbal ability, and the child direct genetic effect for 57% and 79%, respectively.

### Psychopathology

For psychopathology, maternal and paternal genetic nurture estimates were 0.04 and 0.02, and the child direct genetic effect was 0.05. The population prediction was 0.08. Each psychopathology outcome was predicted by its own PGS, with parent–child PGS correlations averaging 0.47 for mothers and 0.52 for fathers. After applying these correlations, 37% of the population prediction for psychopathology can be attributed to genetic nurture and 63% to the child direct genetic effect. Supplementary Figures 2 and 3 show the estimates for each of the eight psychopathology outcomes.

### Height and BMI

For height, maternal and paternal genetic nurture estimates were 0.06 and 0.03, and the child direct genetic effect was 0.35. The population prediction was 0.40. With parent–child PGS correlations of 0.55 for mothers and 0.52 for fathers, 12% of the population prediction for height can be attributed to genetic nurture and 88% to the child direct genetic effect.

For BMI, maternal and paternal genetic nurture were −0.01 and 0.01, and the child direct genetic effect was 0.30 (Table 2). The population prediction was 0.30. BMI was predicted by the childhood BMI PGS at age 7 and by the adult BMI PGS at later ages, with parent–child PGS correlations of 0.51 and 0.55 for mothers and 0.52 at both ages for fathers (Table 1). After applying these correlations, 0% of the population prediction for BMI can be attributed to genetic nurture and 100% to the child direct genetic effect (Figure 3).

### Additional genetic nurture analyses

#### Duo and structural equation models

The results described above used complete trios. However, TEDS contains a further 1514 children with maternal but not paternal DNA and 99 children with paternal but not maternal DNA, which are discarded when analyses are restricted to complete trios. While duo models cannot separately identify child direct and parental genetic nurture effects unless maternal and paternal effects are assumed equal ^45^, requiring both parents to provide DNA may itself select the sample. Supplementary Note 1 examines the robustness of the results by comparing duo and trio models, and duo and trio families (Supplementary Figures 4–5 and Supplementary Table 2). Additionally, we used all children with at least one genotyped parent in a three-group structural equation model (SEM) in which complete trios, mother–child duos and father–child duos are incorporated ^46^ (Supplementary Note 2, Supplementary Figures 6–12 and Supplementary Tables 4–6). Duo models systematically attenuated parental genetic nurture estimates, although using duo families did not significantly change the results. The point estimates were largely unchanged as well in SEM, while the standard errors of maternal estimates decreased by about a third.

#### Siblings

We also explored sibling indirect genetic effects because siblings might genetically ‘nurture’ each other. Not including siblings in the trio model when they are present could affect the estimates of parental genetic nurture ^31^. Using up to 466 dizygotic quad families, we added the co-twin’s PGS to the trio model. The co-twin’s PGS coefficient therefore represents the sibling PGS prediction of the index twin’s outcome independently of the index twin’s own and parental PGS. The co-twin’s PGS had an average coefficient of 0.01 (range −0.09 to 0.11; Supplementary Table 2 and Supplementary Figure 13). Notably, accounting for sibling effects left the maternal and paternal estimates largely unchanged, although standard errors were wider in this quad model because of the smaller sample size.

#### Sensitivity analyses

Sensitivity analyses tested the effects of principal component adjustment, sex, zygosity, and restriction to unrelated trios. Point estimates were consistent with the main results, with higher standard errors when the sensitivity subsamples were more restricted (Supplementary Figures 14–16).

#### Mediation by SES

We explored the extent to which genetic nurture operates through family socio-economic status (SES). The analyses, described in Supplementary Note 3 (Supplementary Table 7 and Supplementary Figures 17–19), indicated that for educational and cognitive outcomes, genetic nurture was largely mediated by SES, with the child direct genetic effect largely unchanged. For psychopathology, height, and BMI, there was no evidence of SES mediation, consistent with the weak associations of these parental PGS with SES.

The next section presents estimates of C for the same sample using the same measures at the same ages in order to compare estimates of genetic nurture and C across domains.

### Shared environmental influence (C)

We used univariate twin models to estimate the proportions of phenotypic variance attributable to additive genetic (A), shared environmental (C), and nonshared environmental (E) influences, as applied to the subset of TEDS families with genotyped parents (up to 476 monozygotic and up to 1259 dizygotic twin pairs; Supplementary Figure 20 and Supplementary Table 8). A robustness check using the whole TEDS sample regardless of the presence of DNA for parents or twins yielded similar results: 48 estimates of A and C correlated at 0.93 for A and 0.92 for C (up to 2391 monozygotic and up to 4319 dizygotic twin pairs; also see Methods, Supplementary Figures 21, 22 and Supplementary Table 9).

Figure 4 shows the ACE estimates for the eight trait domains averaged over age. C was 17.0% for educational achievement, 24.9% for educational attainment, 22.9% for general cognitive ability, 24.1% for verbal ability, and 16.5% for nonverbal ability. In contrast, C was negligible (1.7%) for psychopathology. For height and BMI, C was 11.2% and 7.2%, respectively. ACE results for all 48 variables are shown in Supplementary Figure 20.

**Figure 4.**
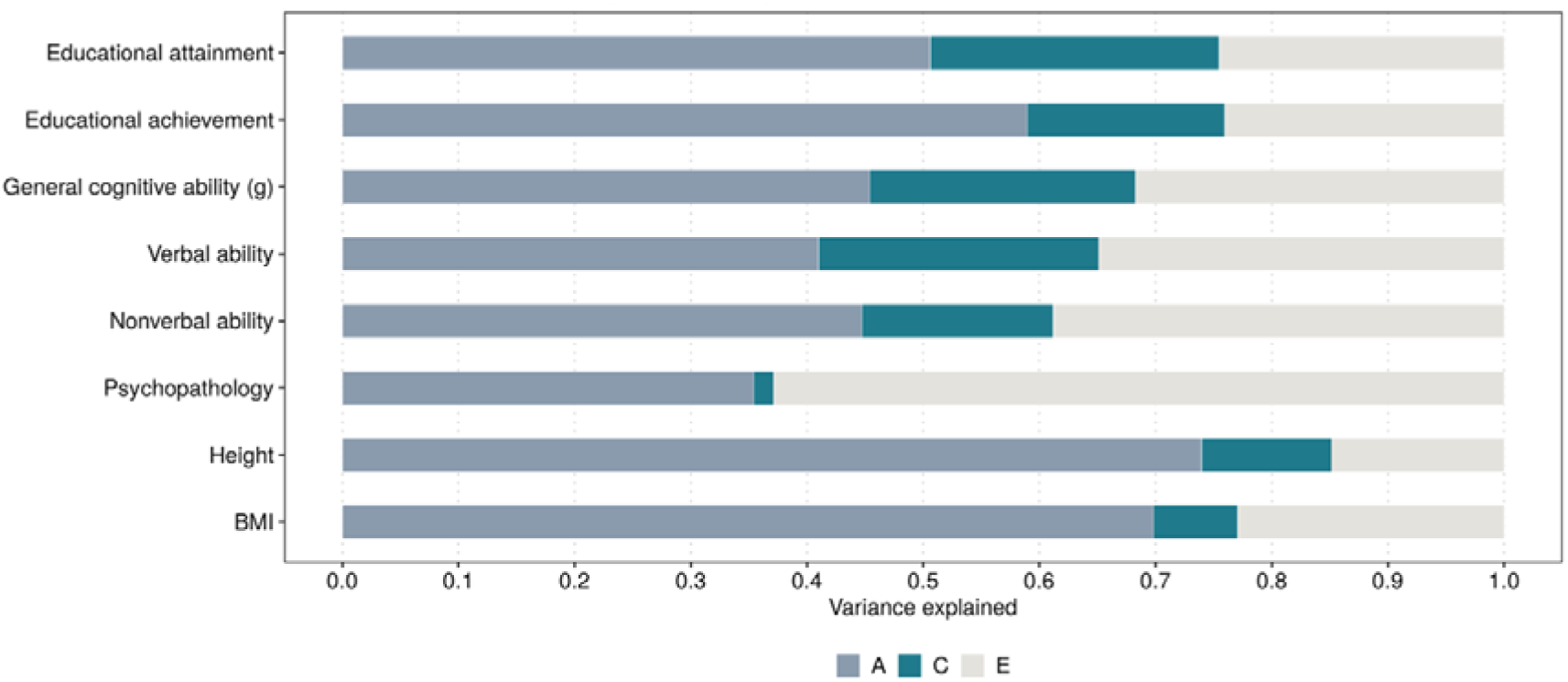
Twin additive genetic (A), shared environmental (C), and nonshared environmental (E) estimates. Bars show the proportion of phenotypic variance attributable to additive genetic, shared environmental, and nonshared environmental influences, which sum to one. Estimates are from univariate twin models fitted in TEDS families in which the offspring were genotyped and at least one parent provided DNA (151 to 476 monozygotic and 295 to 1259 dizygotic twin pairs). This subsample includes both duo and trio families and is therefore broader than the complete-trio sample used for the genetic nurture analyses. For the 46 continuous measures, phenotypes were residualised on age and sex and standardised before modelling. A-level enrolment at age 16 and university enrolment at age 18 are binary and were modelled using a liability threshold model with age and sex as covariates. These two estimates refer to the latent liability rather than the observed dichotomy. Although different models were used, the estimates are essentially comparable. Sample size for each trait, point estimates, and 95% confidence intervals are presented in Supplementary Table 8.

### Comparing genetic nurture and shared environment

Figure 5 shows the scatterplot of the relationship between estimates of genetic nurture (from Figure 3) and shared environmental influence (from Figure 4). C path coefficients rather than variance explained (c^2^) are used, so that both C and genetic nurture are expressed in standard deviation units.

**Figure 5.**
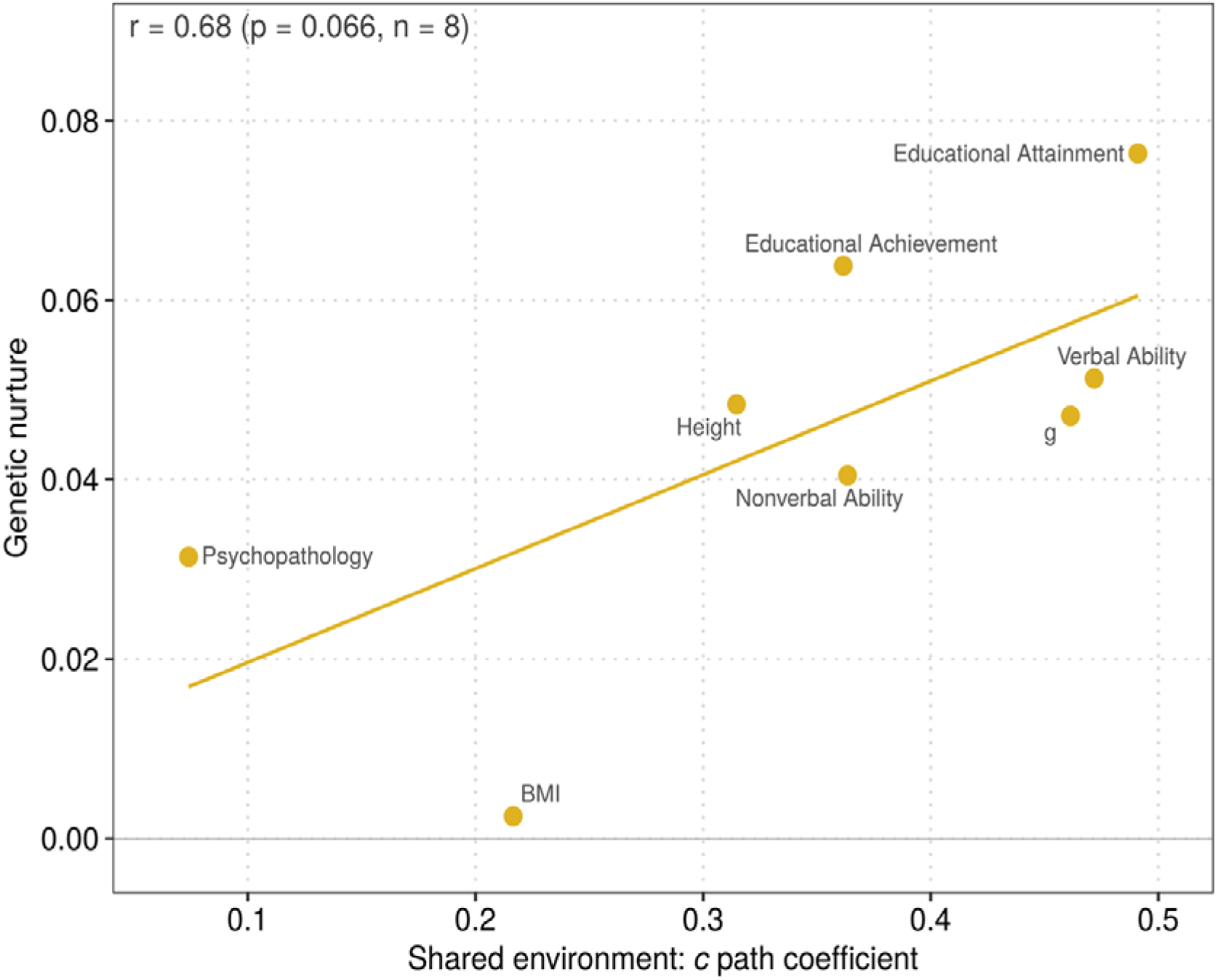
Scatterplot between genetic nurture and shared environment. Each point represents one of eight trait domains. The x-axis shows the shared environmental path coefficient, calculated as the square root of c² (Figure 4) and averaged within each domain. The y-axis shows the sum of maternal and paternal genetic nurture within the population PGS prediction (Figure 3). Both axes are on a standard deviation scale. The line shows the least-squares regression across the eight domains (r = 0.68, p = 0.07, n = 8). Correlations of maternal and paternal genetic nurture with shared environment are shown separately in Supplementary Figure 23.

Despite the small and generally nonsignificant point estimates of genetic nurture for the 48 variables, averaging the point estimates by domain shows a relationship between genetic nurture and C. Educational attainment and educational achievement, which show higher estimates of genetic nurture, also show high C, followed closely by verbal ability and general cognitive ability. Nonverbal ability shows less C and somewhat less genetic nurture, followed by height, and anchored by BMI, which shows some C but no genetic nurture. Although psychopathology is in the lower left quadrant of the scatterplot, it deviates from the regression by showing some genetic nurture despite showing the least C. However, interpretation of the regression between C and genetic nurture is limited given the modest estimates of genetic nurture that range only from 0.00 to 0.08 in standardised units.

Nonetheless, the correlation between genetic nurture and C for these eight domains is 0.68, suggesting a substantial relationship. However, a perspicacious reader may have noticed in Figure 3 that genetic nurture is correlated with predictive power of the PGS, which creates a ceiling for estimates of genetic nurture. Most notably, PGS prediction for psychopathology is only a third that of educational PGS. In addition, the greater estimates of genetic nurture for educational variables as compared to cognitive abilities could be due to greater PGS predictive power of the educational attainment PGS as compared to the cognitive abilities PGS.

For this reason, we reconsidered genetic nurture proportionate to the population PGS prediction. This transformed the scatterplot in Figure 5, as shown in Figure 6. Proportionate to population polygenic score prediction, the most striking change is that psychopathology now shows the most genetic nurture, suggesting that with an equally powerful PGS, psychopathology might show as much genetic nurture as educational and cognitive traits. In addition, genetic nurture is now comparable for cognitive and educational variables.

**Figure 6.**
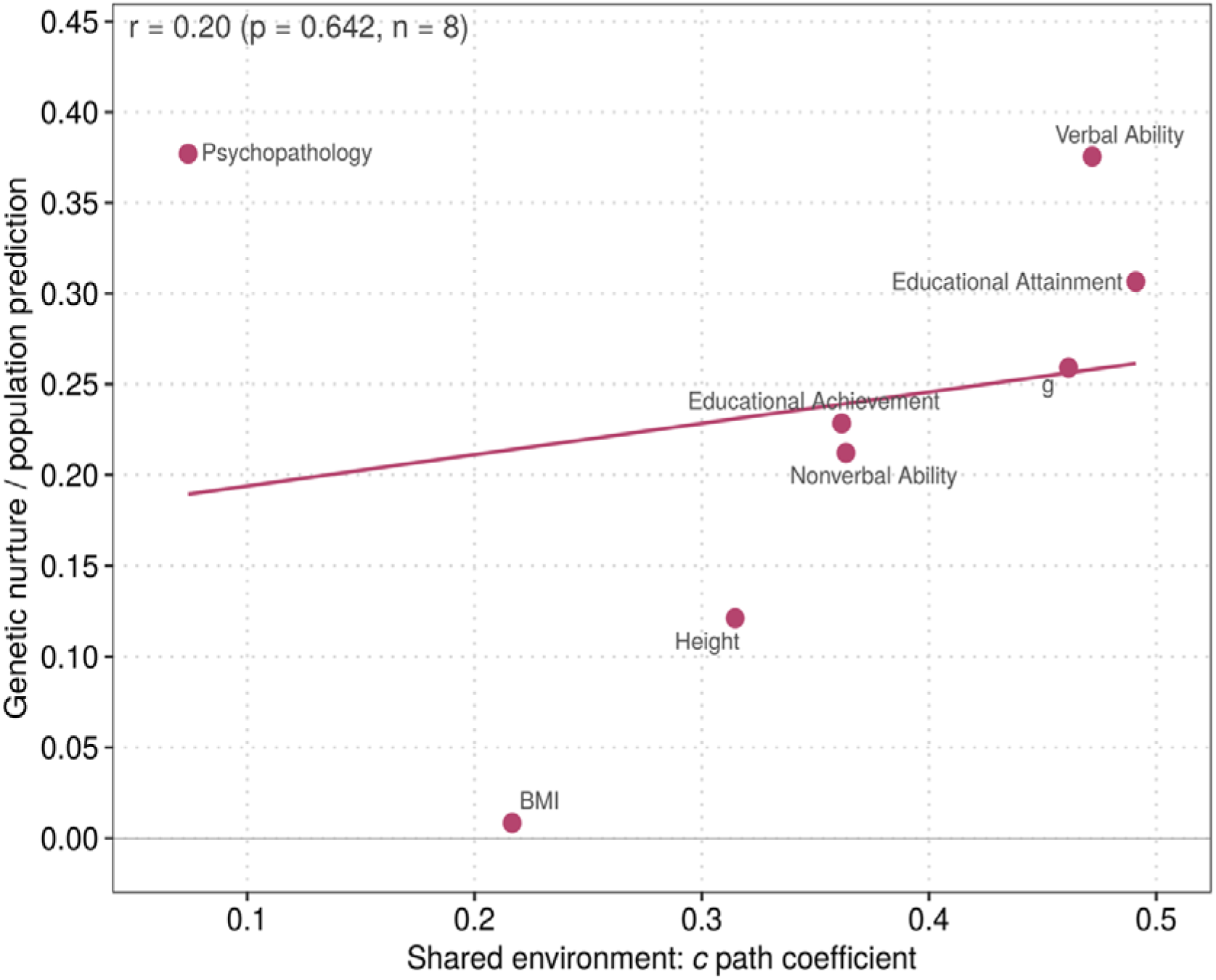
Scatterplot between genetic nurture as a proportion of the population PGS prediction and shared environment. Each point represents one of eight trait domains. The x-axis shows the shared environmental path coefficient, calculated as the square root of c² (Figure 4) and averaged within each domain, as in Figure 5. The y-axis shows genetic nurture within the population PGS prediction divided by the population PGS prediction (Figure 3). Estimates were averaged within each domain before the ratio was calculated. The line shows the least-squares regression across the eight domains (r = 0.20, p = 0.64, n = 8). Across the seven domains other than psychopathology, the correlation was 0.95 (p = 0.001). Correlations of maternal and paternal genetic nurture with shared environment are shown separately in Supplementary Figure 24.

## Discussion

Using genotyped parents and their twin children from a longitudinal twin cohort, we systematically examined the influences of genetic nurture and shared environment (C) for educational achievement, educational attainment, cognitive abilities, psychopathology symptoms, and height and BMI.

### Small effects of genetic nurture

Our most general finding, in line with the literature, is that genetic nurture effects are very small (Figure 3), accounting for less than ∼2% of the phenotypic variance. Genetic nurture is the product of three ingredients: a heritable component of the environment, the ability of the parental PGS to predict this heritable component of the environment, and a correlation between these unmeasured environmental factors and children’s outcomes. The product of reasonable estimates of these three ingredients is a very small number. Environmental measures widely used in behavioural research show substantial genetic influence ^47^, but the average twin heritability of these environmental measures is about 25% ^48^, half that of the average heritability of behavioural measures ^49^. Current PGS are unlikely to account for more than 10% of the heritable influence on environmental measures, just as PGS for behavioural traits generally account for less than 10% of the variance. Environmental measures such as parenting account for only about 5% of the variance in children’s psychopathology ^50,51^. The product of these three chains is a very small number (0.25 twin heritability of environmental measures × 0.10 heritable variation in environmental measures accounted for by the PGS × 0.05 variance in children’s psychopathology explained by environmental measures = 0.1%). However, a benefit of genetic nurture is that it is not limited by measured environments; these unmeasured environments could potentially account for much more variance in children’s outcomes. Even doubling all three estimates in recognition of the benefit of incorporating unmeasured environments, the product of these three chains (0.50 × 0.20 × 0.10) is only 1%, the typical effect size of genetic nurture.

Our study was powered to detect genetic nurture effects that account for 1.5% of the variance, but only 4 of 48 variables yielded effects of this magnitude and the largest effect was 2.1% ^35^. The small genetic nurture effects meant that power became a major limitation for our analyses because detecting such small genetic nurture effects with adequate power requires at least twice as many trios as in our sample. We alleviated this problem by focusing on effect sizes averaged across variables in a domain rather than on point estimates for the 48 variables with the goal of describing the general domain landscape of genetic nurture.

### The relationship between genetic nurture and C

Our twin study estimates of C are substantial, explaining up to 25% of the total trait variance. The pattern across the eight domains – with a high of 25% for educational attainment and a low of less than 2% for psychopathology – corresponds to the research literature ^1^. Although the small estimates of genetic nurture make it difficult to investigate whether estimates of genetic nurture are related to C, a general pattern emerged (Figure 5) in that educational attainment, which shows the most C, also shows the highest estimates of genetic nurture, followed closely by verbal ability and general cognitive ability, which also show substantial C. Nonverbal ability shows slightly less genetic nurture and less C, and educational achievement shows more genetic nurture but a comparable C. An intriguing but tenuous finding is that height suggests some genetic nurture and also some C. BMI corresponds to the trend in that it shows no genetic nurture and little C (7%), which is surprising because for decades it had been assumed that BMI runs in families for reasons of shared environment ^52^. Psychopathology at age 26 is the only domain that shows no C, which corresponds to previous research. Although psychopathology shows some genetic nurture, the correlation between genetic nurture and C for these eight domains is 0.68 (Figure 5).

However, Figure 3 shows that estimates of genetic nurture are also associated with the predictive power of the domain’s PGS. For the behavioural domains, the strongest PGS prediction is for the educational variables and the weakest is for psychopathology. When we considered genetic nurture as a ratio to PGS prediction in the population, the association between genetic nurture and C disappeared (Figure 6). This loss of the correlation, however, is entirely due to psychopathology. Without psychopathology, a very strong association emerged between C and the ratio of genetic nurture to the population PGS prediction (r = 0.95). The most striking change was that, proportionate to its PGS prediction, psychopathology showed as much genetic nurture as educational attainment despite showing the least C. The implication is that if the PGS for psychopathology were as predictive as the PGS for educational attainment, psychopathology would show as much genetic nurture as educational variables. This implication assumes that a more predictive psychopathology PGS would increase the direct and genetic nurture components in proportion to their current contributions, although it is possible that larger GWAS could instead raise prediction primarily through direct effects or between-family effects.

How is it possible for psychopathology to show genetic nurture given that it shows no C? Genetic nurture requires C – it captures a portion of C that can be tagged by parents’ measured genotypes. A seemingly unrelated question is why PGS prediction for psychopathology remains so low despite large GWAS sample sizes. We suggest that nonadditive genetic variance could be the answer to both questions.

### How can psychopathology show genetic nurture in the absence of C?

The answer to this question might be that C in fact affects psychopathology but has been underestimated in twin studies because of nonadditive genetic variance. Nonadditive genetic variance reduces the genetic coefficient of relationship for DZ but not MZ twins, so that the hallmark of nonadditive genetic variance is a pattern of twin correlations in which MZ > 2DZ. In this way, nonadditive genetic variance can mask C, which is identified as MZ < 2DZ ^53^. Twin studies of psychopathology symptoms provide some evidence for nonadditive genetic variance, especially for ADHD ^54^, although estimates are inconsistent across measures and studies and are difficult to distinguish from several alternative explanations in the classical twin design ^1^. Specifically, the classical twin design cannot estimate C and nonadditive genetic variance together with additive genetic variance. When other family and adoptive relatives are added to the twin design, they permit estimates of C in addition to additive and nonadditive genetic influence. These studies find evidence of C of about 10%, compared with the 2% estimated in the current study, in addition to nonadditive genetic influence ^55–58^.

Based on these findings, we suggest that there is C for psychopathology and that evidence for genetic nurture will be found when PGS for psychopathology become more powerful and when developmentally specific psychopathology phenotypes are examined. We admit that the small effect sizes for genetic nurture mean that our reasoning can at best be viewed as hypotheses for future research. One hypothesis is that genetic nurture will be similar in magnitude for psychopathology as for educational and cognitive traits when their estimates of genetic nurture are indexed as a ratio to the predictive power of their PGS. Another testable hypothesis is that C, a prerequisite for genetic nurture, will be found in quantitative genetic analyses of psychopathology that include other family members in addition to twins, which enables simultaneous analyses of C and nonadditive genetic variance.

If psychopathology shows more nonadditive genetic variance than educational and cognitive traits, the same logic could explain why PGS for psychopathology predict less variance than PGS for educational and cognitive traits. GWAS and PGS assume additivity. Thus, their ceiling is the additive heritability, which is lower for traits that show more nonadditive genetic variance. Although attempts to integrate dominance ^59^ and epistasis ^60^ in GWAS of psychopathology have not improved PGS prediction, such analyses are challenged by daunting demands for power ^61,62^.

### SES mediation of genetic nurture

While a merit of the genetic nurture framework is that it can detect an environmental effect without directly measuring the environment responsible, to further contextualise the genetic nurture effect, studies have identified a measurable pathway through which genetic nurture likely operates: socio-economic status ^9,10,63^. Our results indicated that SES mediated genetic nurture for cognitive and educational outcomes, but not for height, BMI, or psychopathology. We interpret this mediation with caution: SES reflects partly heritable parental characteristics such as parental education, occupation and family income ^64^. Consequently, SES may be a measurable mediator of genetic nurture without being an environmental pathway.

Additionally, what we attribute to parents might in part be dynastic, running through wider family lines rather than through parents themselves ^7,65^. Moreover, assortative mating might ‘create’ genetic nurture through genetic effects untagged by the child’s PGS ^65^, and a portion of this genetic nurture may load on A, the additive genetic component. C estimates are also not immune to assortative mating. Therefore, some of these seemingly environmental effects may reflect genetic effects of the wider family and may not be modifiable.

## Limitations and conclusion

Our study also has several limitations in addition to those related to the predictive power of current PGS and the small effect size of genetic nurture, which we have highlighted. One general limitation is that our sample consists entirely of White British participants. More specific limitations include our focus on eight general trait domains rather than point estimates of all 48 variables because of our lack of power, although we include results for the 48 variables. Another limitation is our focus on within-trait analyses rather than including cross-trait analyses, which was an attempt to simplify our presentation. Additionally, we only examined psychopathology at age 26, when shared environmental influences are lower for other domains as well; genetic nurture effects may be more prominent at earlier ages.

Nevertheless, our study is unique in bridging quantitative and molecular genetics to empirically explore the relationship between genetic nurture and C. It raises novel issues for psychopathology, such as a greater role for genetic nurture when considered in relation to PGS predictive power, a greater role for C which has been underestimated in twin studies, and the possible role of nonadditive genetic variance in accounting for both of these issues.

## Methods

### Participants

We used participants from the Twins Early Development Study (TEDS), a longitudinal twin cohort born in England and Wales between 1994 and 1996 ^39^. A total of 13,759 families took part. The twins have been assessed repeatedly since infancy, with data collection ongoing. Following standard TEDS practice, we excluded twin pairs in which either twin had a severe medical condition or severe perinatal complications, or in which zygosity was unknown (1850 individuals). Ethical approval was granted by the King’s College London Research Ethics Committee (PNM/09/10–104 and HR/DP-20/2122060), and informed consent was obtained at each wave of data collection.

Genotype data were available for 10,346 individual twins, who provided DNA through buccal swabs or saliva samples across several waves and were genotyped in two batches on Affymetrix and Illumina arrays ^66,67^. Both the full cohort and its genotyped subsample remain broadly representative of the UK population for this birth cohort.

DNA was subsequently collected from 3789 parents of twins using saliva samples, and 3543 parents were retained after quality control. Parent and twin genotype data were then harmonised and merged. Parent–offspring relationships were inferred from genome-wide genotype data. To ensure that all family units were independent, we removed 15 families (31 individuals) that were related to another family in the sample (third-degree relatives or closer). Full details of genotypic information are provided in Supplementary Method 1.

The final sample comprised 2559 families in which at least one parent was genotyped (2460 with maternal and 1045 with paternal DNA), including 946 complete parent–offspring trios (Supplementary Table 10). All participants in the present study were of European genetic ancestry.

Compared with genotyped children whose parents were not genotyped, children with at least one genotyped parent scored higher on educational attainment (mean Cohen’s d = 0.31), educational achievement (0.24) and cognitive abilities (0.20), and lower on psychopathology (−0.15) and adult BMI (ages 22 and 26; −0.16), whereas height did not differ (Supplementary Table 11). They also came from families with higher SES (mean d = 0.35). Their own polygenic scores showed a similar pattern on a smaller scale: scores for cognitive ability and educational attainment were slightly higher, scores for four of the eight psychopathology traits (PTSD, anxiety, depression and ADHD) were lower, and scores for height and BMI did not differ. These differences may reflect selection of the families who took part in parental DNA collection, which has implications for the generalisability of our results. During the DNA collection, mothers were encouraged to participate including follow-up reminders. Mothers were also asked to engage the biological father of the twins but this phase of DNA collection was not followed up in case there were concerns about paternity.

Additionally, among children without genotyped parents, those with and without their own genotype data did not differ in SES at birth or in psychopathology, while cognitive and educational outcomes from adolescence onwards were slightly higher in children without genotype data (d = −0.11 to −0.23; Supplementary Table 12).

### Polygenic Scores

Using the merged parent–offspring genotype data described above, we calculated polygenic scores for all genotyped twins and parents, based on the latest GWAS summary statistics of each trait (Figure 2). We used LDpred2-auto ^68,69^, a Bayesian approach that re-weights SNP effects from GWAS summary statistics to account for linkage disequilibrium, drawing on a reference built from the HapMap3+ variant set. The auto version infers from the summary statistics each trait’s genetic architecture, or its SNP heritability and the fraction of variants with non-zero effects, so the scores did not need to be tuned in a held-out sample. Polygenic scores were based on approximately 1.1 million SNPs.

To control for population stratification, each polygenic score was residualised on the first ten genetic principal components of the individual to whom it belonged: child scores on the child’s own principal components, and maternal and paternal scores on the mother’s and father’s own principal components. The principal components were computed in an unrelated subset comprising one randomly selected child per family and then projected onto the full sample (Supplementary Method 1). All subsequent analyses used these residualised scores, and principal components were not entered again as covariates.

We focused on within-trait comparisons between PGS and traits. A polygenic score for general cognitive ability (g) ^70^ predicted g, and verbal ability and nonverbal ability at ages 7, 9, 10, 12, 16 and 25. A polygenic score for educational attainment ^71^ predicted primary school grades, General Certificate of Secondary Education (GCSE) grades, A-level enrolment, A-level grades, university enrolment, university grades and years of schooling. A height polygenic score ^72^ predicted height at ages 7, 12, 14, 16, 22 and 26. A childhood BMI polygenic score ^73^ predicted BMI at age 7, and an adult BMI polygenic score ^74^ predicted BMI at the remaining ages. For psychopathology, symptoms of post-traumatic stress disorder (PTSD), depression, alcohol use, anxiety, eating disorders, ADHD, bipolar disorder and autism were each predicted by the corresponding polygenic score ^75–82^.

### Phenotypes

We analysed 48 outcomes across eight trait domains (Figure 2); Supplementary Method 2 describes each measure in full.

In childhood (ages 7, 9 and 10), verbal ability was assessed with tests from the Wechsler Intelligence Scale for Children (WISC) ^83^. Nonverbal ability was assessed with WISC subtests and other nonverbal tests, including Raven’s Progressive Matrices at age 10 ^84^. General cognitive ability (g) was the standardised mean of the verbal and nonverbal scores. In adolescence, verbal ability was assessed with language and reading tests at age 12 and the Mill Hill Vocabulary test ^85^ at age 16, and nonverbal ability with mathematics tests at age 12 and Raven’s Progressive Matrices at age 16. At age 12, g was assessed with a separate set of WISC and Raven’s tests rather than derived from verbal and nonverbal scores. In adulthood (age 25), cognitive abilities were assessed with Pathfinder, a gamified online battery of verbal and nonverbal tests ^86^.

Educational achievement comprised school grades from age 7 to 21: teacher-rated primary school grades at ages 7, 9, 10 and 12, GCSE grades at age 16, A-level grades at age 18, and university grades at age 21.

Educational attainment included A-level and university enrolment as well as years of schooling, derived from the highest qualification reported at age 26, or at age 21 when age-26 data were unavailable. We grouped the two enrolment outcomes with years of schooling because they mark successive transitions that make up educational attainment: A-levels are required for university entry, and university enrolment is required for a degree.

Psychopathology was assessed at age 26 using self-report questionnaires: the 6-item PTSD Checklist (PCL-6) ^87^ for PTSD symptoms, the Generalized Anxiety Disorder – Dimensional scale (GAD-D) ^88^ for anxiety symptoms, the shortened Mood and Feelings Questionnaire (MFQ) ^89^ for depressive symptoms, the Alcohol Use Disorders Identification Test (AUDIT) ^90^ for alcohol use, the inattention scale of the Conners 3rd Edition Self-Report ^91^ for ADHD symptoms, the 6-item Ritvo Autism Asperger Diagnostic Scale (RAADS) ^92^ for autistic traits, the symptom checklist of the Mood Disorder Questionnaire (MDQ) ^93^ for bipolar symptoms and the Psychiatric Genomics Consortium (PGC) eating disorder screener for eating disorder symptoms.

Height and weight were assessed at ages 7, 12, 14, 16, 22 and 26. BMI was calculated as weight in kilograms divided by height in metres squared.

Family SES was a standardised composite of mother and father education, mother and father occupation, and household income, assessed at birth and at ages 7, 16 and 21.

### Power

To obtain 80% power to detect genetic nurture effects with a trio-PGS design, approximately 1000 trios are required for a trait with a strong PGS, such as educational attainment (assuming PGS R² = 7.5%, genetic nurture from both parents combined explaining 3% of the variance), 2500 trios for a trait with a moderate PGS, such as personality (R² = 2.5%, parental effects 1%), and 7500 trios for a trait with a weak PGS, such as ADHD (R² = 1%, parental effects 0.4%) ^35^. With 946 complete trios, our sample was therefore close to adequately powered for educational outcomes, less so for cognitive outcomes, and underpowered for psychopathology.

### Analytical Approaches

All analyses were conducted using R version 4.6.0.

### Data preparation

Missing data were handled separately for each participant and each measure. A scale score was computed only if the participant answered at least half of its items; for example, at least 6 of 11 items were required. For composite measures made up of several tests or scales, such as cognitive abilities, the same rule was applied to the subscales.

All phenotypes and polygenic scores were standardised using the mean and standard deviation of the unrelated subsample (one twin per family), and these values were applied to all participants. Age was entered as a covariate on its original scale. Model coefficients are therefore comparable to Pearson’s r, or to point-biserial correlations for the binary outcomes.

### Descriptive statistics

Descriptive statistics for each phenotype at each age are presented for complete trios and for all families with at least one genotyped parent, using one child per family (Supplementary Tables 13 and 14). About half of the phenotypes were associated with sex and nearly two thirds with age (Supplementary Table 15). Sex and age were therefore included as covariates, together with genotyping chip. Zygosity explained at most 1% of the variance in any phenotype and 1% in family SES at birth, falling to 0.1% by age 7. We therefore examined zygosity in a sensitivity analysis, fitting the models separately in monozygotic and dizygotic twins. Polygenic scores, residualised on the first ten principal components, were largely independent of the covariates (Supplementary Table 16).

### Regression models

We estimated genetic nurture with trio polygenic score regressions (Figure 1). For each outcome, we fitted two models. The population model regressed the outcome on the child’s polygenic score. The trio model regressed the outcome on the child’s, mother’s and father’s polygenic scores for the same trait:

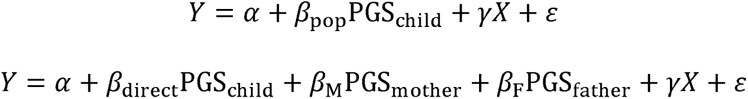

where *X* denotes the covariates: child age, sex and genotyping chip. At the population level, the child’s coefficient *β*_po*p*_ represents the correlation between PGS and the phenotypic outcome. Conditional on the parental scores, the child’s coefficient *β*_direct_ represents the child direct genetic effect. The parental coefficients *β*_M_ and *β*_F_ represent maternal and paternal genetic nurture, respectively.

Both dizygotic twins were included where available to increase statistical power. Standard errors and 95% confidence intervals for the coefficients were obtained from a cluster-robust (sandwich) estimator ^40,41^. Percentile confidence intervals for the ratios of coefficients were obtained from 1000 cluster bootstrap resamples of families.

The two binary enrolment outcomes were analysed with the same linear regression models as the continuous outcomes, that is, as linear probability models ^94^. In complete trios, 75% of the twins enrolled in A-levels and 69% in university. At these proportions, linear probability models give results close to those of logistic regression.

To relate the population and trio models, as demonstrated in the Results section, the population coefficient can be expressed as *β*_po*p*_ ≈ *β*_direct_ + *r*_MC_*β*_M_ + *r*_FC_*β*_F_. Here *r*_MC_ and *r*_FC_ are the mother–child and father–child polygenic score correlations (Table 1). Parental genetic nurture enters the population coefficient through each parent’s PGS correlation with the child. Because the parent–child correlations are close to 0.5, each parent’s genetic nurture contribution to the population prediction is about half of that parent’s coefficient.

Moreover, because the population model is nested within the trio model, the incremental R² from adding the parental PGS indicates how much parental genetic nurture adds to the prediction of the child’s outcome beyond the child’s own PGS (Supplementary Table 2 and Supplementary Figure 25). However, this incremental R² is not the variance explained by genetic nurture, because part of genetic nurture is already carried into the population R² by the child’s PGS. Hence, we expressed the coefficients as proportions of outcome variance by squaring them. The variance explained by genetic nurture was 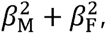 and that by the child direct genetic effect was 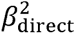 10.

We focused on effect sizes rather than statistical significance. P values for each coefficient were obtained from the models and were adjusted for multiple testing (see Multiple testing correction).

### Sensitivity analyses for genetic nurture

#### Duo models

We used duo models, which include the child’s PGS and one parent’s PGS, for two comparisons (Supplementary Note 1). First, to examine the consequences of omitting one parent, we fitted the duo and trio models in the same sample of complete trios. Second, to separate the effect of the model from that of the sample, we compared the duo models fitted in complete trios with the same duo models fitted in all families in which that parent was genotyped (2460 mother–child and 1045 father–child duos). Because the duo models are nested within the trio model, these comparisons also show how much each parent adds to prediction. For example, the incremental R² from the mother–child duo model to the trio model is the father’s unique contribution (Supplementary Figure 25). As noted above, however, incremental R² does not capture the full variance explained by a parent’s genetic nurture.

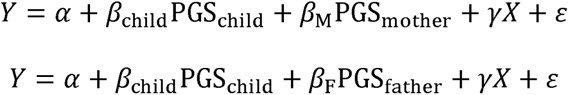

#### Structural equation model

To make use of all families with parental DNA, we fitted a three-group structural equation model ^46^. Complete trios, mother–child duos and father–child duos contributed jointly through full information maximum likelihood, and regression coefficients, latent variances and latent covariances were constrained to be equal across groups (Supplementary Note 2). Models were fitted in *lavaan* with a robust maximum likelihood estimator and standard errors clustered by family ^95^.

#### Sibling indirect genetic effects

Indirect genetic effects of siblings may affect estimates of parental genetic nurture ^31^. We therefore added the dizygotic co-twin’s polygenic score to the trio model in families in which both parents and both dizygotic twins were genotyped (up to 466 families).

#### Subgroups

We fitted the trio model separately in female and male offspring, without sex as a covariate; in monozygotic and dizygotic twins; in one child per family (unrelated subsample); and with polygenic scores that were not residualised on principal components.

#### Mediation by SES

We tested whether genetic nurture operated through family SES with a mediation model in which the child’s, mother’s and father’s PGS predicted both SES and the outcome (Supplementary Figure 17; Supplementary Note 3). Indirect effects were the products of the paths from each parental PGS to SES and from SES to the outcome. The main analysis used SES at birth, and a sensitivity analysis used the most recent SES assessment before each outcome (Supplementary Figures 26 and 27).

### Twin ACE analyses

We estimated additive genetic (A), shared environmental (C) and nonshared environmental (E) components with univariate twin models. Models were fitted in OpenMx ^96^. Continuous outcomes were residualised on age and sex and standardised before modelling. The two binary enrolment outcomes were analysed with liability threshold models with age and sex as covariates, so their estimates refer to the underlying liability. Because both sets of estimates are proportions of variance, they are directly comparable.

The main twin analyses used families in which the twins and at least one parent were genotyped. These are the families that contribute to the genetic nurture analyses. As a robustness check, we fitted the same models in the whole TEDS twin sample.

### Aggregation of results and comparison of genetic nurture with C

Results were averaged within eight trait domains: general cognitive ability, verbal ability, nonverbal ability, educational achievement, educational attainment (A-level enrolment, university enrolment and years of schooling), psychopathology, height and BMI (Figure 2). We first averaged the coefficients within each domain and then calculated variance components and ratios from these averages. These ratios of averages were close to the corresponding averages of ratios.

### Multiple testing correction

P values for coefficients from the regression, structural equation and mediation models were adjusted with the Benjamini–Hochberg false discovery rate ^97^. Significance after adjustment is indicated in the Supplementary Figures and Tables.

