## Supplementary Information for "Genetic nurture and shared environmental effects across cognitive, educational, and mental health outcomes"

### Supplementary Methods

#### Supplementary Method 1. Genotypic Information

##### The TEDS Parents Genotyping and Quality Control

###### *DNA Sample Collection*

DNA for the TEDS parents was extracted from a total of 3789 saliva samples at the National Institute for Health Research Biomedical Research Centre for Mental Health Bioinformatics Core at the Social, Genetic and Developmental Psychiatry Centre. Following DNA extraction, 3722 samples passed processing. DNA quality control (QC) indicated that 2712 samples had a DNA concentration >50 ng/µL, 817 samples had a concentration between 20 ng/µL and 50 ng/µL, and 193 samples had a concentration <20 ng/µL.

###### *Genotyping and Quality Control*

The DNA samples were then hybridised to Illumina genotyping microarrays (genotype chip: GSAMD.24v3.0.EA_20034606_A1.csv). The Illumina-specific software GenomeStudio was used to call genotypes, with SNP coordinates mapped to the Genome Reference Consortium Human Build 37 (GRCh37) [1]. A total of 3708 samples and 730,059 SNPs were initially subjected to QC. After manual review in GenomeStudio, 31,093 SNPs failed QC, leaving 698,966 SNPs that passed QC.

Subsequently, the data were processed through a custom automated QC pipeline. First, samples with a call rate below 90% were removed in GenomeStudio to prevent interference with the GenCall clustering algorithm. Then, a stricter sample call rate threshold of 95% was applied. A total of 162 samples were removed, and no duplicated samples were identified, resulting in a final sample size of 3546.

The zCall algorithm was applied to uncalled SNPs in GenomeStudio [2]. An optimal threshold of z=5 was calculated and used to distinguish rare allele signals from background noise. The zCall output was then converted to PLINK1.9 binary format [3], and A/B alleles were updated to the Illumina TOP strand based on the genotyping chip used. This rescued genotype calls for rare variants that were not assigned by the standard clustering algorithm, maintaining the total SNP count of 698,966.

Further iterative filtering was applied. This involved alternating thresholds of SNP call rate (>95%) and sample call rate (>95%) to prune the data [4]. This pruned dataset was used solely to flag potentially problematic samples. Five sex discrepancies were identified, where genetically inferred sex (based on X-chromosome F-statistics) differed from the reported sex [5]. Additionally, 65 heterozygosity outliers (>3 standard deviations) were found. High levels of heterozygosity can indicate samples of low quality, while low levels can be due to inbreeding [6]. Outliers were observed in both directions, indicating both poor sample quality and potential inbreeding. Lastly, four unique related samples were identified via Identity by Descent (IBD) estimation. Given the primary aim of constructing a merged family dataset, these individuals were flagged but retained, as downstream analyses account for relatedness and sample quality.

###### *Ancestry Proportions*

Ancestry was estimated via principal component (PC) projection analysis using the 1000 Genomes Project Phase 3 as a reference panel [7]. First, linkage disequilibrium

(LD) pruning was performed using a window size of 1500 kb, a step size of 150 variants, and an $R^2$ threshold of 0.2. Known high-LD genomic regions and non-autosomal SNPs were excluded, leaving a total of 192,079 variants. Second, principal components were computed on the reference panel to establish a stable population structure reference space. The TEDS parents were then projected onto these reference PCs using PLINK1.9.

To assign super-population groups, a penalised multinomial regression model was trained on the first 10 PCs of the reference panel using the *glmnet* R package [8]. A classification probability threshold of 0.5 was applied for group assignment. Based on this predictive model, 3544 individuals were robustly assigned to the European (EUR) reference population, and one individual was assigned to the South Asian (SAS) cluster (Supplementary Figure 1). One individual did not meet the probability threshold for any population cluster and remained unclassified. All individuals were retained for subsequent imputation regardless of ancestry assignment.

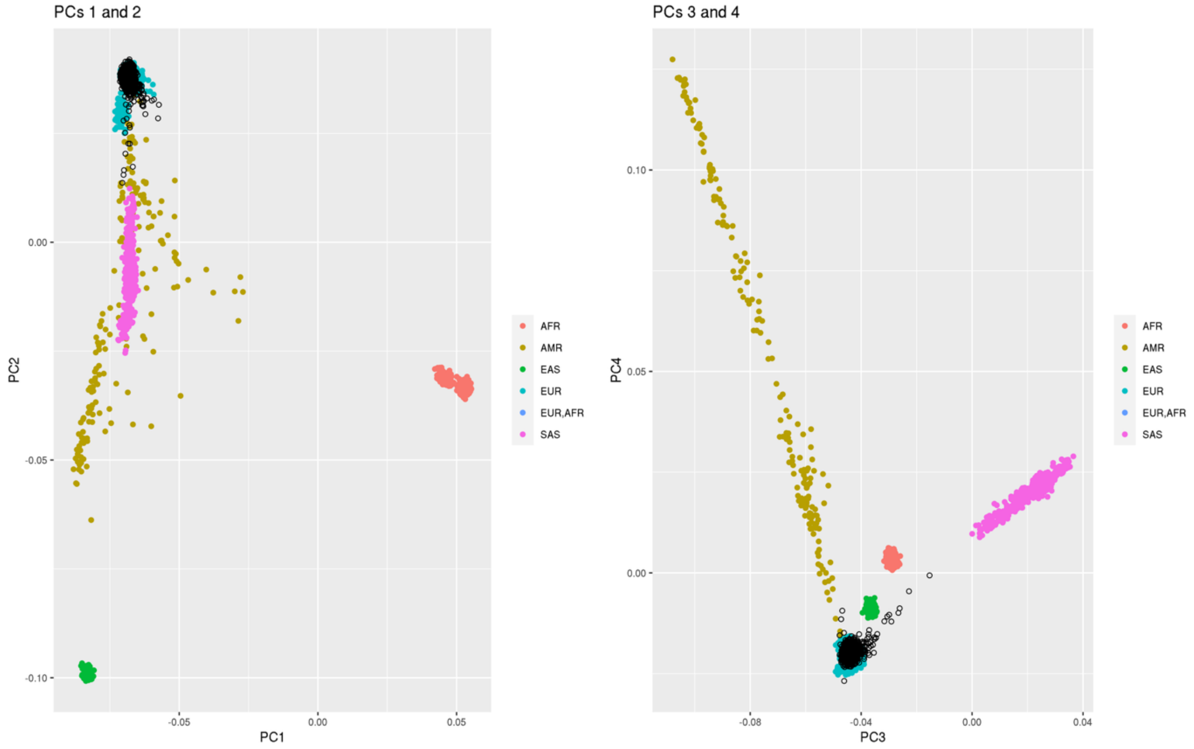

**Supplementary Figure 1. Ancestry Estimation Using Principal Component Analysis.** Plots display PC1 vs PC2 (left) and PC3 vs PC4 (right). Reference populations from the 1000 Genomes Phase 3 panel are coloured by super-population: AFR = African; AMR = Admixed American; EAS = East Asian; EUR = European; EUR,AFR = admixed European–African; SAS = South Asian. The TEDS parent genotyped samples are superimposed and coloured in black.

##### *Imputation*

Prior to imputation, the genotyped parental dataset (N = 3546) was lifted over [9,10] to the Genome Reference Consortium Human Build 38 patch release 14 (GRCh38.p14) [11]. A total of 696,147 SNPs were retained after lifting over, followed by additional quality control. SNPs with minor allele frequency (MAF) < 0.01, SNP and sample call rate < 95%, and deviation from the Hardy–Weinberg equilibrium

(HWE) exact test ($p < 1\times10^{-6}$) were excluded. The final pre-imputation dataset comprised 502,744 SNPs and 3546 individuals.

Genotyped data were phased using Eagle2 [12] and imputed into the TOPMed Reference Panel (Freeze 8) [13] via the Michigan Imputation Server (pipeline version 1.7.1) [14]. Imputation was performed using Minimac4 (v1.0.2).

Post-imputation quality control was applied by excluding SNPs with an imputation quality score ($R^2$) < 0.3, MAF < 0.001, SNP and sample missingness > 5%, and deviation from HWE ($p < 1\times10^{-10}$), resulting in 14,798,531 SNPs retained for downstream analyses.

**Overview of TEDS Twin Genotypic Data**
Genotypic data for the TEDS twins have been extensively described in previous publications and are summarised here for completeness [15,16]. DNA samples were collected across multiple waves using buccal cheek swabs and saliva kits, and genotyping was conducted in two primary batches using Affymetrix and Illumina platforms.

The first genotyping phase was carried out as part of the Wellcome Trust Case Control Consortium phase 2 (WTCCC-2), using the Affymetrix Genome-Wide Human SNP Array 6.0. The second phase (OEE study) expanded the sample using the Illumina HumanOmniExpressExome array. Standard QC procedures were applied within each dataset, including filtering based on sample call rate, minor allele frequency, HWE, relatedness, heterozygosity, and ancestry.

Following platform-specific quality control, genotype data from both platforms were phased using Eagle2 and imputed to the Haplotype Reference Consortium (HRC, release 1.1) reference panel via the Sanger Imputation Service [12,17,18]. Post-imputation filtering and harmonisation across platforms resulted in a final dataset comprising 7,363,646 SNPs and 10,346 individuals.

Full details of DNA collection, genotyping, quality control, and imputation procedures are available in the TEDS data dictionary (https://datadictionary.teds.ac.uk/studies/dna.htm) and prior publications.

**Harmonisation and Merging of TEDS Parent and Twin Genotype Data**
To enable joint analyses of the TEDS parents and twins, genotype data from both datasets were harmonised and merged following imputation and quality control.

As the TEDS twin data were imputed on the GRCh37 reference genome [1], whereas the parent data were imputed on GRCh38 [11], genomic coordinates of the twin dataset were lifted over to GRCh38 using the UCSC liftOver tool (https://genome.ucsc.edu/cgi-bin/hgLiftOver) with the hg19ToHg38 chain file [9,10]. Variants that could not be successfully mapped during liftOver were flagged and excluded in subsequent steps. Following liftOver and removal of unmapped variants, 7,360,741 SNPs remained in the twin dataset.

Variants were then matched between the parent and twin datasets based on chromosome and base-pair position, yielding 7,311,628 overlapping SNPs. Among these, allele alignment was carefully evaluated to ensure consistency between datasets. Allele coding was compared using the *snp_match* function in the *bigsnpr* R package [19], which identifies variants that match directly, after allele reversal (swapping of reference and alternative alleles), or after strand complementation. Strand-ambiguous (palindromic) variants (i.e., A↔T, C↔G) were removed, as a strand flip cannot be distinguished from an allele swap. Inconsistent variants between datasets after accounting for reversal and complementation were excluded. Palindromic variants account for the majority of the reduction at this step. Following harmonisation, 6,198,700 SNPs were retained that showed consistent genomic position and allele coding across both datasets.

Prior to merging, multiallelic variants identified in the parental genotype data (i.e., variants with more than two alleles across datasets) were excluded, resulting in 6,190,706 SNPs. Although no multiallelic variants were present in the processed twin dataset, the twin SNP set was restricted to the same 6,190,706 variants retained in the parental data to ensure consistency across datasets. In addition, three parents were excluded from the parental dataset due to ambiguous sex assignment based on X chromosome data, resulting in a final parental sample of 3543 individuals to be combined with 10,346 twins. The final merged dataset comprised 6,190,706 SNPs across 13,889 individuals.

***Relatedness Checks and Quality Control***

Following the merging of the TEDS parent and twin datasets, relatedness within the combined sample was assessed using KING [20]. Prior to this stage, parents and twins had been treated as unrelated individuals; KING was applied post-merge to identify and resolve cryptic relatedness across the full dataset. Individuals inferred as parent–offspring pairs (InfType = PO) were assigned as the parents of the corresponding twin family. Parental sex was inferred from X-chromosome homozygosity (F statistic; $F > 0.8$ classified as male, $F < 0.2$ as female). This determined whether each parent was the female or male parent of the family.

Relatedness inference was conducted iteratively. In the first stage, KING was run on the full merged dataset (N = 13,889), identifying three parental samples as genetic duplicates of existing twin participants (Dup/MZ, PropIBD > 0.95). These individuals had been enrolled as both a parent and a twin and were removed, reducing the sample to 13,886. In the second stage, KING was rerun with up to second-degree relatedness inference. Ten pairs of between-family relatives were identified beyond those accounted for by the provided pedigree. These comprised two pairs of full siblings among parents (InfType = FS, PropIBD > 0.45) and additional second-degree relative pairs. For each pair of related families, the smaller family unit was removed to maximise retention of complete trios, resulting in the exclusion of nine individuals across four families (N = 13,877). In a third stage, 22 individuals belonging to families with third-degree relatedness across family units were further removed to ensure independence of family units for downstream trio-based analyses, yielding a final relatedness-checked sample of 13,855.

Standard genotypic quality control was then applied, filtering SNPs on HWE ($p < 1\times10^{-6}$), genotype missingness (> 5%), and minor allele frequency (< 0.01). Five individuals with individual-level genotype missingness exceeding 5% were also removed. Following the extraction of SNPs passing imputation quality (INFO > 0.3) and MAF filters (> 0.01), the final dataset comprised 5,439,847 SNPs and 13,850 individuals, including 2460 female parents, 1045 male parents, and 946 complete trios.

***Principal Component Analysis***

Principal components (PCs) were computed on an unrelated subset comprising one randomly selected child per family, following LD pruning (window = 50 SNPs, step = 5, $R^2 < 0.2$). PCs were then projected onto the full merged sample using PLINK2 score-based projection [21], with variant loadings and reference allele frequencies derived from the unrelated subset. The first 10 PCs were retained as covariates for downstream analyses.

***Preparation of Genotype Data for Polygenic Score Calculation***

For polygenic score computation using LDpred2 [22], which requires complete genotype data with no missing calls, a separate version of the dataset was prepared. As the parent and twin genotype data were generated on different platforms and imputed to different reference panels, residual missingness was present predominantly in the parental data following merging. Missing genotype calls were filled with the homozygous reference allele using the PLINK --fill-missing-a2 function. This approach is acknowledged as a simplification that discards missingness information, and the resulting dataset is therefore reserved exclusively for LDpred2 polygenic score computation and is not used in any other downstream analyses.

### Supplementary Method 2. Phenotypic Information

Phenotypic information used in the present study is summarised below by trait domain. Full documentation is available in the TEDS data dictionary (https://datadictionary.teds.ac.uk/home.htm).

##### Cognitive Abilities

###### *Childhood*

At age 7, cognitive tests were administered by telephone. Verbal ability was assessed with the Similarities and Vocabulary subtests of the Wechsler Intelligence Scale for Children, third edition (WISC-III) [23], and nonverbal ability with a Conceptual Grouping test and the WISC-III Picture Completion subtest [24].

At ages 9 and 10, verbal ability was assessed with the Vocabulary and General Knowledge subtests of the WISC-III as a Process Instrument (WISC-III-PI) [25]. Nonverbal ability was assessed with the Figure Classification and Figure Analogies subtests of the Cognitive Abilities Test 3 (CAT3) at age 9 [26], and with WISC-III-UK Picture Completion and Raven's Progressive Matrices at age 10 [27]. Tests were completed in booklets sent by post at age 9 and online at age 10.

###### *Adolescence*

At age 12, verbal ability was assessed with three language tests, covering syntax [28], semantics and pragmatics [29], and four reading tests: the Peabody Individual Achievement Test [30] and the GOAL Formative Assessment in Literacy [31] for comprehension, and the Woodcock-Johnson III Reading Fluency test [32] and the Test of Word Reading Efficiency [33] for fluency. Nonverbal ability was assessed with mathematics tests from the National Foundation for Educational Research [26]. Verbal and nonverbal scores were summed across their component tests. General cognitive ability was assessed separately with WISC-III-PI Vocabulary and General Knowledge, WISC-III-UK Picture Completion and a pattern test adapted from Raven's Standard Progressive Matrices [23,27], and g was the mean of the four standardised test scores.

At age 16, verbal ability was assessed with the Mill Hill Vocabulary test [34] and nonverbal ability with Raven's Standard and Advanced Progressive Matrices [27].

At ages 7, 9, 10 and 16, verbal and nonverbal composites were the standardised means of their component tests, and g was the standardised mean of the two composites.

###### *Adulthood*

At age 25, cognitive abilities were assessed with Pathfinder, a gamified online measure developed by the TEDS researchers [35]. The 20 verbal items came from Mill Hill vocabulary, missing letter and verbal reasoning tests, and the 20 nonverbal items from Raven's Standard Progressive Matrices and three visual puzzle tests on analogies, grouping and logical sequences. Verbal and nonverbal scores were the numbers of correct items (0 to 20 each), and g was their sum (0 to 40).

##### Educational Achievement

At ages 7, 9, 10 and 12, teachers rated achievement in English and mathematics, and from age 9 also in science, against the National Curriculum levels (https://www.gov.uk/national-curriculum/overview). Ratings ranged from 0 to 4 at age 7 and from 0 to 9 at later ages, and achievement was the mean of the standardised subject ratings.

At age 16, achievement was the mean grade in the three compulsory General Certificate of Secondary Education (GCSE) subjects, English, mathematics and science, reported by the twins or their parents and coded from 4 (G) to 11 (A*). GCSEs are national examinations taken at the end of compulsory education in the UK.

At age 18, A-level achievement was the mean self-reported grade across A-level subjects, coded from 1 (E) to 6 (A*). A-levels are two-year qualifications taken after compulsory education and required for university entry.

At age 21, university degree classification was self-reported on a scale from 1 (lowest pass) to 5 (first-class honours) [36].

**Educational Attainment**

At age 18, two binary measures were recorded. Twins who reported at least one A-level or Advanced Subsidiary (AS) level were coded as enrolled in A-levels [37], and twins who went on to university were coded as enrolled in university, as opposed to employment, training or other destinations.

Years of schooling was derived from the highest qualification reported at age 26, or at age 21 when age-26 data were missing; highest qualifications at the two ages correlated at $r = 0.86$ [15]. Qualifications were converted as follows: GCSE, 10 years; A-level, 12 years; higher education below degree level, 14 years; bachelor’s degree, 15 years; master’s degree, 16 years; doctorate, 20 years. No qualification was treated as missing.

**Psychopathology**

All psychopathology measures were self-reported at age 26. A scale score required responses to at least half of its items, except where noted, and higher scores indicate more symptoms.

PTSD symptoms over the past month were measured with the six-item PTSD Checklist (PCL-6), a short form of the civilian PTSD Checklist [38], rated on five points from ‘not at all’ to ‘extremely’.

Anxiety symptoms over the past seven days were measured with the 10-item Generalized Anxiety Disorder – Dimensional scale (GAD-D) [39], rated on five points from ‘never’ to ‘all of the time’.

Depressive symptoms over the past two weeks were measured with the 13-item Mood and Feelings Questionnaire (MFQ) [40], rated as not true, sometimes true or true.

Alcohol use was measured with the Alcohol Use Disorders Identification Test (AUDIT), which covers drinking frequency and quantity and indicators of hazardous or harmful use in the past year [41].

ADHD symptoms were measured with the 11 inattention items of the Conners 3rd Edition self-report scale [42], rated on four points from 'not true at all' to 'definitely true'.

Autistic traits were measured with the RAADS-6, a six-item short form of the Ritvo Autism Asperger Diagnostic Scale derived by TEDS researchers from the RAADS-14 [43]. Three items cover social interaction and communication, and three cover sensory sensitivity, attention to detail and resistance to change. Each item is rated by whether it applied never, only in childhood (up to age 16), only now, or both, so that higher totals indicate more persistent traits.

Bipolar symptoms were measured with the Mood Disorder Questionnaire (MDQ) [44]. Its first section asks whether each of 13 manic or hypomanic symptoms has ever been experienced; later sections are shown only to participants who endorse symptoms. Because the MDQ has no standard dimensional score, we used the number of symptoms endorsed in the first section, as derived by the TEDS data management team.

Eating disorder symptoms were measured with the screener developed by the Psychiatric Genomics Consortium Eating Disorders Working Group, also used in the UK Biobank. Based on DSM-5 criteria, it covers lifetime symptoms of anorexia nervosa, bulimia nervosa and binge-eating disorder. Follow-up items skipped after a negative answer to a screening question were coded 0, and the score was the mean of the available items. Thus, the score does not distinguish between subtypes of eating disorders.

**Height and BMI**
Height and weight were assessed at ages 7, 12, 14, 16, 22 and 26, and BMI was calculated as weight in kilograms divided by height in metres squared.

**Socio-Economic Status**
Family socio-economic status (SES) was assessed at birth and at ages 7, 16 and 21. Each composite was standardised and calculated from parental occupational status (coded with the UK Standard Occupational Classification, https://www.ons.gov.uk/methodology/classificationsandstandards/standardoccupationalclassificationsoc), parental highest educational qualifications and household income. In the main mediation analyses, SES at birth was used for all outcomes, so that the mediator always preceded the outcome. As a sensitivity analysis, each outcome was instead paired with the most recent SES assessment that did not postdate it: SES at age 7 for outcomes at ages 7 to 14, at age 16 for outcomes at ages 16 and 18, and at age 21 for outcomes from age 21 onwards (Supplementary Note 3).

### Supplementary Notes

#### Supplementary Note 1. Duo versus Trio Models and Families.

Our main analyses used complete trios. However, TEDS contains a further 1514 families with maternal but not paternal DNA and 99 with paternal but not maternal DNA, and restricting analyses to complete trios discards them. While duo models cannot separately identify child direct and parental genetic nurture effects unless maternal and paternal effects are assumed equal [45], requiring both parents to provide DNA may itself select the sample.

To incorporate duo families and increase statistical power, particularly for maternal genetic nurture estimates, we first conducted a model-level robustness check (Supplementary Figure 4). To separate the effect of the model from that of the sample, we fitted the duo model within the complete trio sample, using the same families as the main analysis. Under duo models, maternal and paternal estimates were consistently attenuated, with ratios of the mean estimates to the corresponding trio model estimates of 0.62 and 0.82, respectively. None of the duo estimates differed significantly from its trio counterpart.

We then conducted a sample-level robustness check for duos: those from the complete trio families and those from all families in which that parent was genotyped. The two duo models differ only in sample composition (Supplementary Figure 5). Paternal estimates were largely unchanged (averaged $\beta_{paternal}$=0.04 in both, r=0.98 across estimates), as the average sample size across traits only increased from 885 to 977. Maternal estimates changed with domain: they were larger for educational outcomes (averaged $\beta_{maternal}$=0.00 became 0.04) and height (averaged $\beta_{maternal}$=0.04 became 0.06), and smaller for cognitive abilities (averaged $\beta_{maternal}$=0.03 became 0.01) and psychopathology (averaged $\beta_{maternal}$=0.03 became 0.01). None of these changes was significant. The consistent gain was in precision: as the mean maternal sample rose from 885 to 2129, standard errors decreased by 34%. Therefore, more maternal estimates were statistically significant.

We also examined incremental $R^2$, which addresses a different question: how much each parent's PGS adds to the prediction of the child's outcome (Supplementary Figure 25; Supplementary Table 2). Taking the mother as an example, comparing the child–mother duo model with the population model shows how much adding the mother's PGS to the child's PGS improves the prediction. One potential issue with this comparison is that the mother's PGS also partly indexes paternal genetic nurture through the correlation between the parents' PGS (assortative mating). We therefore also compared the trio model with the child–father duo model, which shows how much the mother's PGS adds beyond both the child's and the father's PGS. The same comparisons were made for fathers. Overall, genetic nurture from both parents added on average 0.4% to the prediction of the child's outcome (range: less than 0.01% to 1.6%), and 0.8% for educational outcomes (0.1% to 1.6%), most of which came from fathers. Each parent's contribution beyond the child's and the other parent's PGS was larger than its contribution beyond the child's PGS alone, consistent with the attenuation of duo estimates described above.

#### Supplementary Note 2. Three-Group Structural Equation Model.

The trio model is the optimal way to estimate genetic nurture. Although duo models attenuate parental estimates, their smaller standard errors indicated a gain in precision. To use the additional 1514 mother–child duos and 99 father–child duos without reintroducing omitted-parent bias, two approaches are available. The first is Mendelian imputation, in which the missing parent's genotype is reconstructed from the observed family members according to Mendelian laws. Where the mother and both twins are genotyped, the father's transmitted alleles are determined and the remaining alleles are imputed using the population allele frequency. Where only the twins are genotyped, the same logic recovers the mid-parent genotype rather than each parent separately. This procedure yields robust and consistent estimates of direct genetic effects for child PGS prediction [45]. It is less suited to the present study aims, however. Simulations show that adding imputed duos improves the power to detect indirect parental effects far less than adding complete trios, reflecting the unavoidable loss of independent parental information, and consequently shifts the ratio of direct to indirect effects in favour of direct effects [46]. This approach therefore has little power to detect genetic nurture effects. We instead took the second approach, a three-group structural equation model, inspired by a similar missing-data SEM framework that combines the likelihoods of complete and incomplete families under a missing-at-random assumption [47]. Because this approach does not reconstruct the missing parent's genotype, we treat it as a 'soft imputation', in contrast to Mendelian imputation.

A multi-group structural equation model (SEM) estimates the same parameters simultaneously in several groups defined by their data structure, with selected parameters constrained to be equal across groups so that groups with incomplete data contribute to the shared estimates [48]. We defined three groups by which parental polygenic scores were observed: complete trios, mother–child duos, and father–child duos, with complete trios as the reference group. Regression coefficients, latent variances and latent covariances were constrained to be equal across groups. The model is estimated by full information maximum likelihood (FIML), which evaluates the likelihood on all observed data without generating a value for the missing parent. Maternal and paternal polygenic scores are entered as single-indicator latent variables. The child's score is regressed on both with paths fixed at 0.5, the expected value under Mendelian transmission, and the two parental variables are allowed to covary to represent assortative mating (Supplementary Figure 6). A robustness check indicated that freeing the transmission paths from 0.5 and allowing them to be estimated left the genetic nurture estimates unchanged. The population-level model has the same structure but includes only the child's polygenic score in the outcome equation, so that its coefficient indexes the population-level prediction rather than the direct genetic effect. Estimation relies on the assumption that parental genotype data are missing at random.

The major advantage of using the SEM over the regression is borrowing information from the duo families. The three-group FIML lets families with only a genotyped mother contribute to the shared $\beta_{maternal}$.

We checked that the model was coded correctly by fitting it as a single group in complete trios, where its estimates should coincide with the regression. Our results indicated that estimates perfectly aligned between the SEM and regression model (Supplementary Figure 7).

With 99 additional father–child duos, paternal estimates from the structural equation model were essentially identical to our main results. Maternal estimates, in contrast, changed in domain-specific ways (Supplementary Figure 8; Supplementary Tables 4 and 5). Across all outcomes, maternal standard errors decreased by 31%, allowing more estimates to reach statistical significance. Height was the clearest case. Its maternal point estimates barely changed between models (0.06 in the trio regression and 0.07 in the structural equation model; Supplementary Table 6), yet maternal genetic nurture became significant at all six ages. The gain in significance therefore came from the greater precision of including all duos, rather than from larger estimates. BMI estimates were equally stable but close to zero throughout, providing no evidence of genetic nurture. Maternal genetic nurture for alcohol use also reached significance, again with an unchanged point estimate and a smaller standard error.

For cognitive outcomes and psychopathology, maternal estimates were slightly lower than in the trio regression (0.02 compared with 0.04 for both), whereas for educational outcomes they were slightly higher (0.06 compared with 0.03). None of these differences was significant based on overlapping confidence intervals, but the same pattern appeared when the regression model was fitted to all duos, suggesting potential differences between families in which only the mother provided DNA and those in which both parents did.

Additionally, we conducted sensitivity tests examining how estimates changed in different subsamples as well as for girls versus boys. Supplementary Figures 9 and 10 show how estimates change as the sample is widened from complete trios only, to unrelated duos, to the whole duo sample. For mothers, precision improved as sample size increased, with mean standard errors falling to 0.80 and then 0.69 of the complete-trio value, and the number of significant estimates rising from none to 10 and then 12. Point estimates were largely unchanged for height and BMI, so the additional significance for these traits reflects precision rather than a shift in effect size. Educational estimates, by contrast, moved upwards (from 0.03 in complete trios to 0.07 and 0.06). For fathers there was no comparable gain, since only 99 additional father–child duos were available. Standard errors were in fact 17% larger in the sample restricted to unrelated individuals than in complete trios, despite the larger number of individuals, because that restriction discards the dizygotic co-twins whose observations contribute independent information. Once co-twins are restored in the whole sample, paternal standard errors and the number of significant estimates return to their complete-trio values.

Supplementary Figures 11 and 12 show estimates fitted separately in female and male offspring. On average, maternal estimates were higher in daughters than in sons for cognitive abilities (0.04 and 0.00, respectively) and educational outcomes (0.06 and 0.05), whereas paternal estimates were higher in sons for educational outcomes (0.07 and 0.12), height (0.01 and 0.07), and BMI (−0.01 and 0.07). Three maternal estimates were significant in daughters and two in sons. However, no paternal estimate was significant in daughters, against seven in sons. These differences were tested directly for each trait using an independent-samples z test on the SEM coefficients, adjusted for multiple testing. None reached significance for either parent. We therefore do not have sufficient evidence to establish sex differences in genetic nurture.

#### Supplementary Note 3. Mediation of Genetic Nurture via Socio-Economic Status.

We explored the extent to which genetic nurture operates through family socio-economic status (Supplementary Figure 17). SES was measured as the standardised mean of parental highest educational qualification, occupational status, and household income, available at four time points: birth and ages 7, 16 and 21. The main analysis used SES measured at birth, which retained the largest analytic sample (N = 525 to 1338 individuals from 377 to 879 families). Results using wave-matched SES are reported in Supplementary Table 7. Correlations between polygenic scores and SES, and between phenotypes and SES, were of similar magnitude at all four time points (Supplementary Figures 26 and 27).

For cognitive abilities and educational outcomes, genetic nurture ran largely through SES (Supplementary Figures 18 and 19). Both parental polygenic scores predicted family SES (paths $a_2$ and $a_3$ in Supplementary Figure 17) and SES predicted the outcome (path b). The indirect effects ($a_2 \times b$ and $a_3 \times b$) accounted for most of the total parental effect, leaving the direct paths close to zero (paths $c_2$ and $c_3$). The child's own polygenic score no longer predicted family SES once both parental scores were in the model (path $a_1$), as expected given that SES is a parental characteristic, while its direct path to the outcome remained substantial (path $c_1$).

For height, BMI, and psychopathology, there was no evidence of SES mediation. Parental polygenic scores were weakly related to SES, and SES was also weakly related to the outcomes. Indirect paths were near zero, so that the small total genetic nurture effects in these domains were carried by the direct path.

### Supplementary Figures

#### Supplementary Figure 1. Ancestry Estimation Using Principal Component Analysis.

*Note*. This figure is presented in Supplementary Method 1, alongside the description of the ancestry analysis.

### Supplementary Figure 2. Population PGS, Genetic Nurture, and Child Direct Genetic Estimates for All 48 Outcomes Across Ages and Trait Domains.

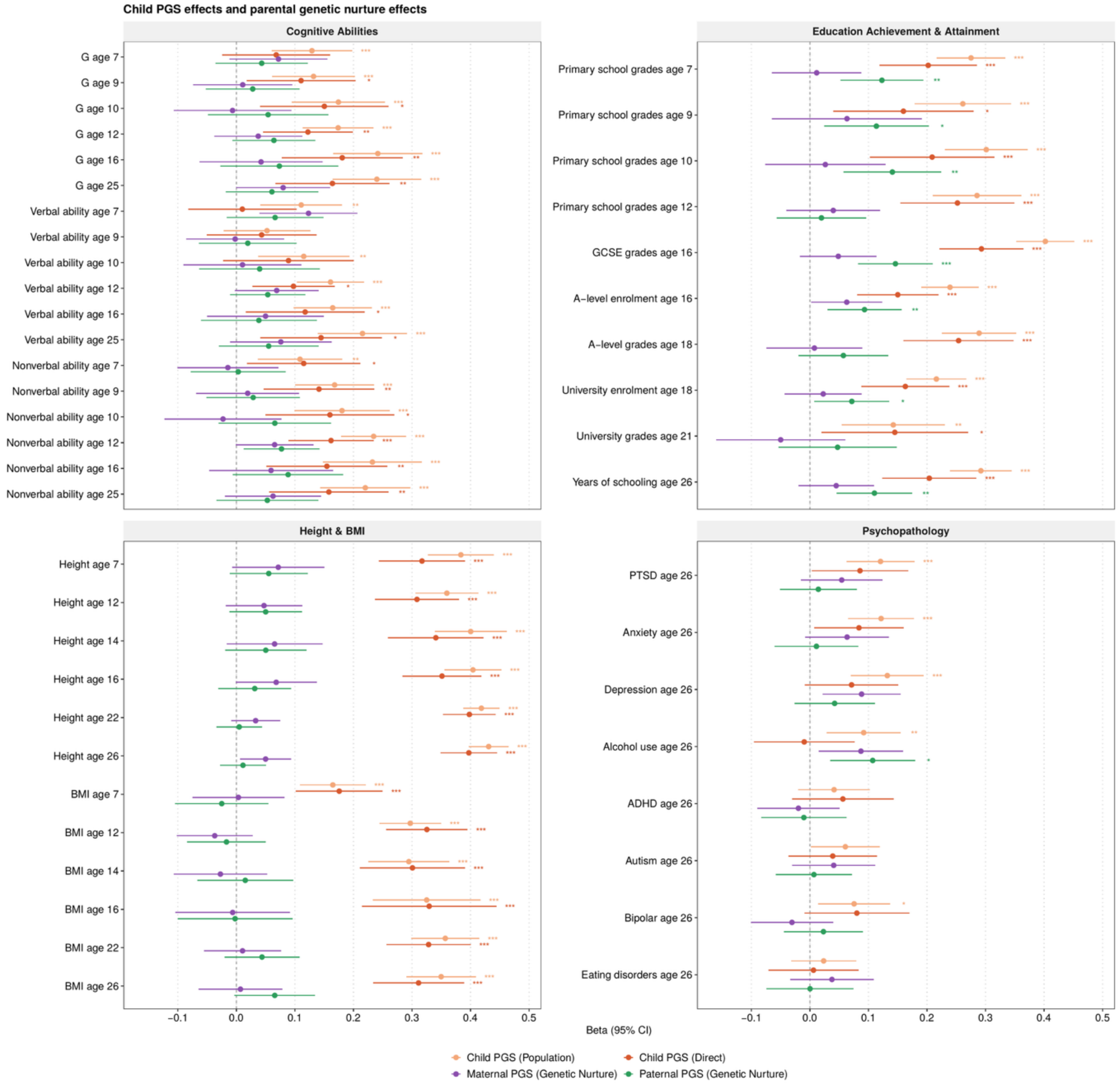

*Note*. Regressions are calculated using complete trios (N = 530 to 1356 individuals from 384 to 893 families), with child age, sex and genotyping chip as covariates. The beta estimates are standardised and considered comparable to Pearson's r coefficient. Error bars represent 95% confidence intervals generated using a cluster-robust sandwich estimator accounting for relatedness within families. Asterisks denote significance after false discovery rate (FDR) correction (Benjamini–Hochberg) applied within each of four broad domains (cognitive abilities, educational achievement and attainment, height and BMI, and psychopathology). Point estimates, standard errors, 95% confidence intervals, variance explained, and adjusted p-values are presented in Supplementary Table 2.
* FDR-adjusted p-value < 0.05; ** < 0.01; *** < 0.001

#### Supplementary Figure 3. Population PGS Prediction Expressed as Child Direct and Genetic Nurture Effects for All 48 Outcomes.

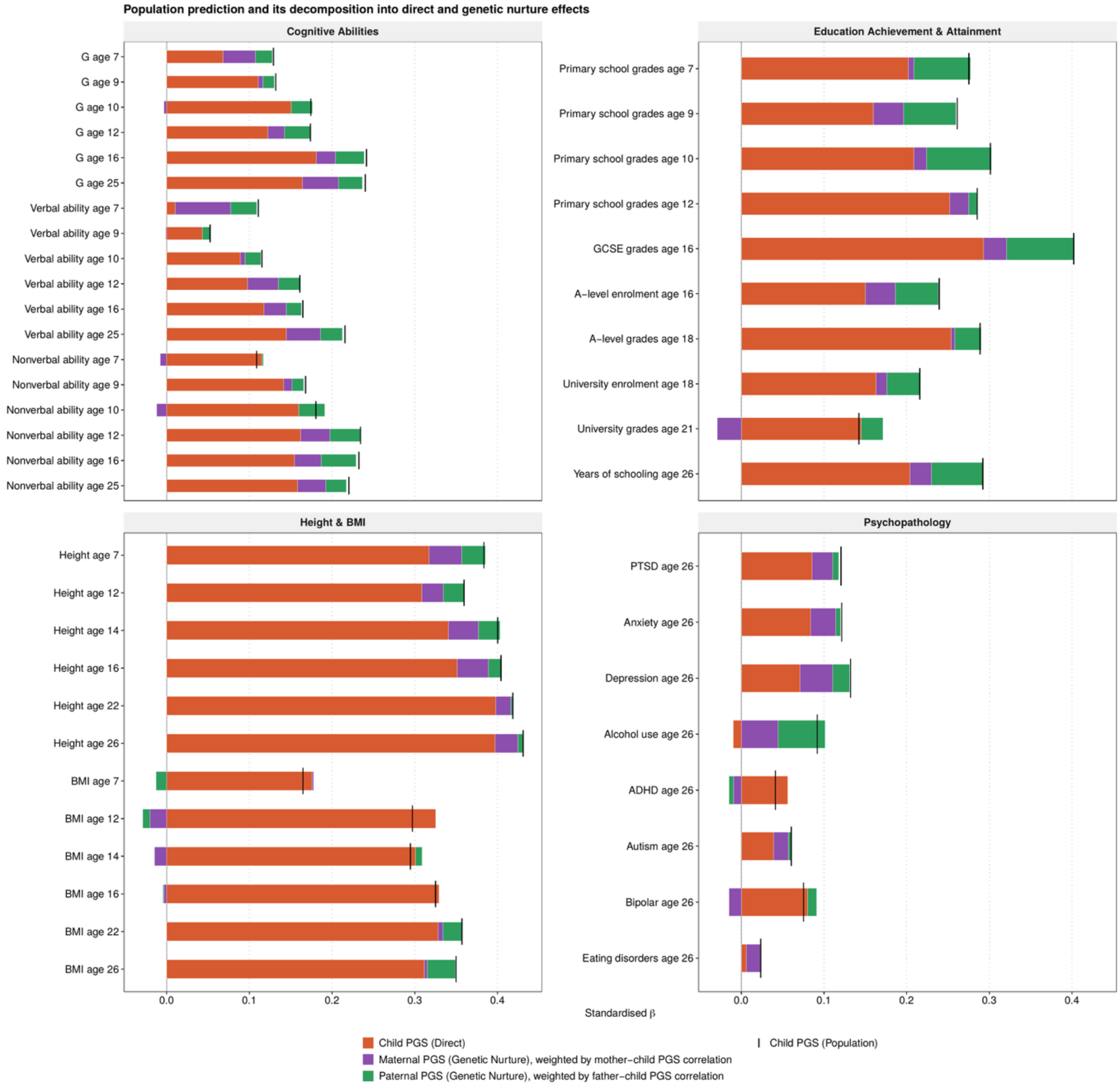

*Note.* For each outcome, bars show the child direct genetic effect and the maternal and paternal genetic nurture effects within the population polygenic score prediction, where each parental coefficient is multiplied by the corresponding parent–child polygenic score correlation (Table 1). The black vertical line marks the population prediction, that is, the standardised coefficient of the child's polygenic score predicting the child's outcome. Regressions are calculated using complete trios (N = 530 to 1356 individuals from 384 to 893 families), with child age, sex and genotyping chip as covariates. Negative components are drawn to the left of zero, so the black line does not always coincide with the end of the bar. Small gaps between the population prediction and the sum of its components arise because the parent–child correlations were estimated in unrelated complete trios, whereas the regression models were fitted in each outcome's analysis sample, which also includes dizygotic co-twins. Figure 3 shows estimates averaged within trait domains. Corresponding estimates are presented in Supplementary Tables 2 and 3.

#### Supplementary Figure 4. Model Comparison: Duo versus Trio Models When Estimating Genetic Nurture Effects.

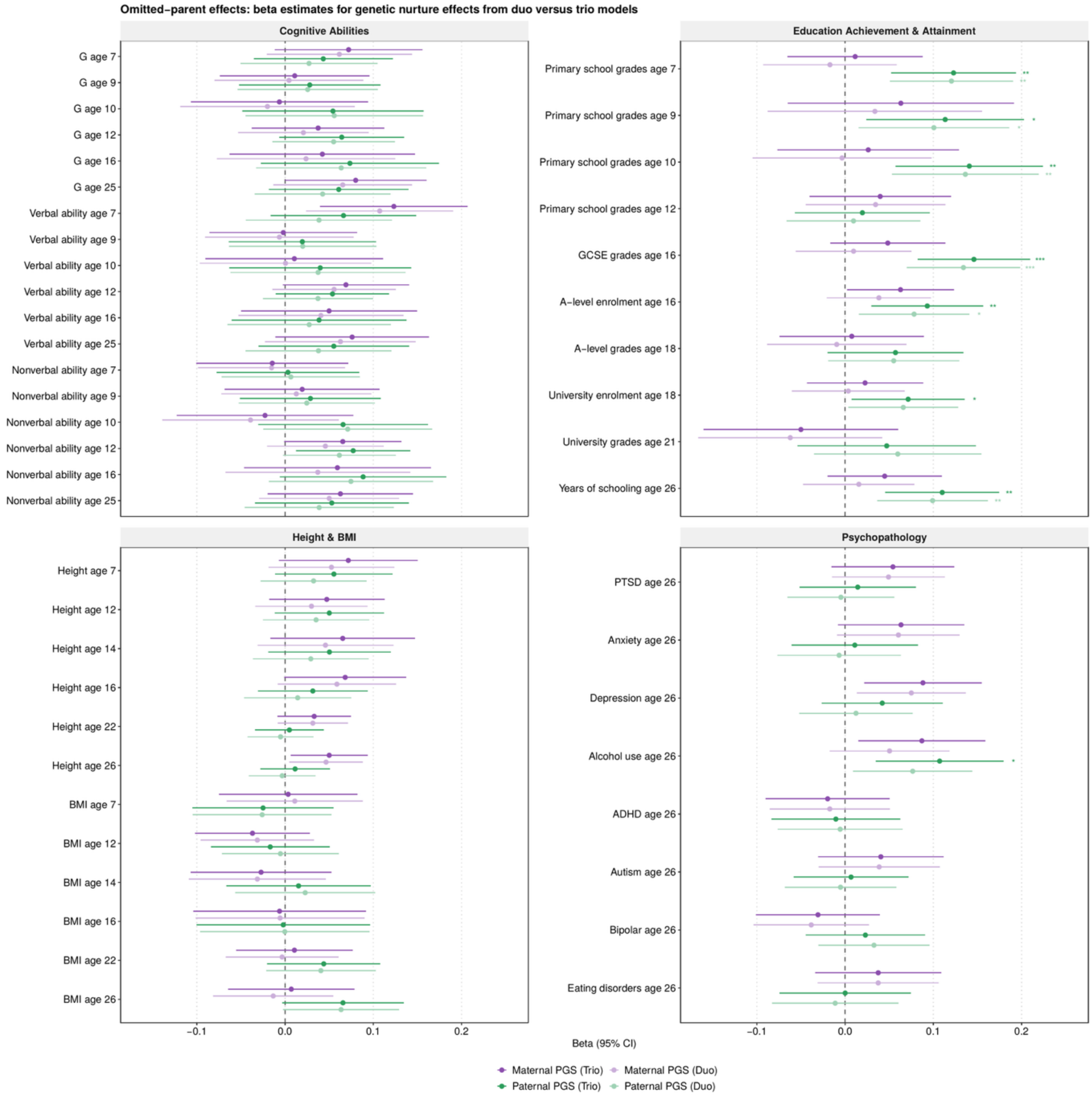

*Note*. Duo and trio regression models are fitted on the same complete trio sample (N = 530 to 1356 individuals from 384 to 893 families), with child age, sex and genotyping chip as covariates. Trio models include the child, maternal and paternal polygenic scores; duo models include the child and one parent only, so that the sample size is identical and the duo models have one fewer parameter. Any difference between the two models therefore reflects the omission of the other parent rather than a loss of participants. Standard errors were slightly smaller under the duo models (mean ratio 0.97), while point estimates were attenuated. Darker shades denote trio model estimates and lighter shades duo model estimates. The beta estimates are standardised and considered comparable to Pearson’s r coefficient. Error bars represent 95% confidence intervals generated using a cluster-robust sandwich estimator accounting for relatedness within families. Asterisks denote significance after false discovery rate correction (Benjamini–Hochberg) applied within each of the four broad domains. Corresponding estimates are presented in

Supplementary Table 2, with domain summaries in Supplementary Table 6.
Discussed in Supplementary Note 1.
* FDR-adjusted p-value < 0.05; ** < 0.01; *** < 0.001

### Supplementary Figure 5. Sample Comparison: Duo and Trio Families When Estimating Genetic Nurture Effects.

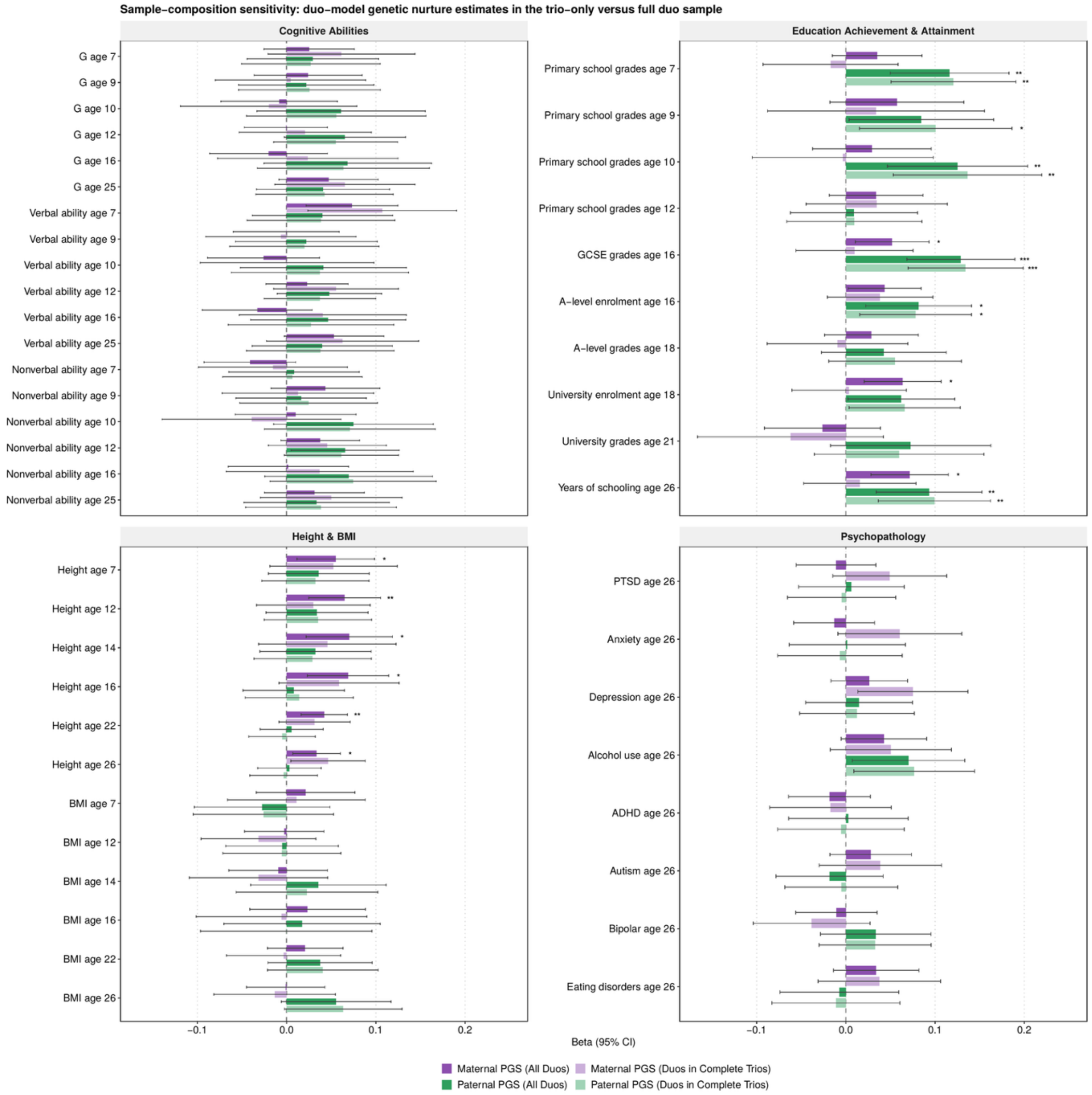

*Note*. The same duo model is fitted to two samples: one restricted to complete trio families (lighter shades; N = 530 to 1356 individuals from 384 to 893 families; the same as the duo subset from Supplementary Figure 4), the other including all families in which that parent provided DNA (darker shades; N = 1257 to 3337 individuals from 918 to 2241 families for mothers and 601 to 1493 individuals from 428 to 984 families for fathers), with child age, sex and genotyping chip as covariates. Because the model is held constant, the omission of the other parent affects both samples equally and the difference between them mainly reflects sample composition rather than model specification. Maternal standard errors decreased as the sample increased, whereas paternal standard errors were largely unchanged, as few additional fathers were available. The beta estimates are standardised and considered comparable to Pearson's r coefficient. Error bars represent 95% confidence intervals generated using a cluster-robust sandwich estimator accounting for relatedness within families. Asterisks denote significance after false discovery rate correction (Benjamini–Hochberg) applied within each of the four broad domains. Corresponding estimates are presented in Supplementary Table 2, with domain summaries in Supplementary Table 6. Discussed in Supplementary Note 1.

* FDR-adjusted p-value < 0.05; ** < 0.01; *** < 0.001

#### Supplementary Figure 6. Three-Group Structural Equation Model.

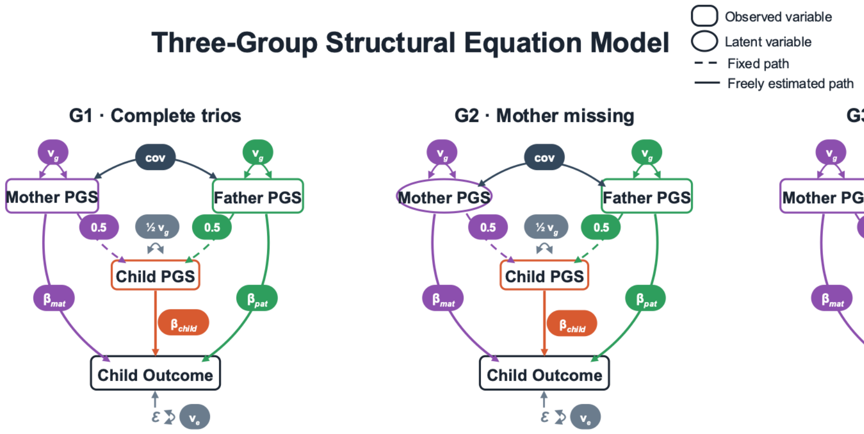

*Note*. Path diagram of the multi-group SEM fitted to the whole sample. Maternal and paternal polygenic scores enter as single-indicator latent variables with residuals fixed at zero. Dashed arrows denote genetic transmission, with paths fixed at 0.5, the expected value under Mendelian transmission. The double-headed arrow between parental variables represents couple PGS correlation. The child's polygenic score predicts the outcome (direct genetic effect) alongside the maternal and paternal variables (maternal and paternal genetic nurture). Child age, sex and genotyping chip are included as covariates. The same structure is fitted in three groups defined by which parental scores are observed: complete trios, mother–child duos and father–child duos, with regression coefficients, latent variances and latent covariances constrained to be equal across groups. Discussed in Supplementary Note 2.

#### Supplementary Figure 7. Three-Group SEM and Regression Give the Same Estimates in Complete Trios.

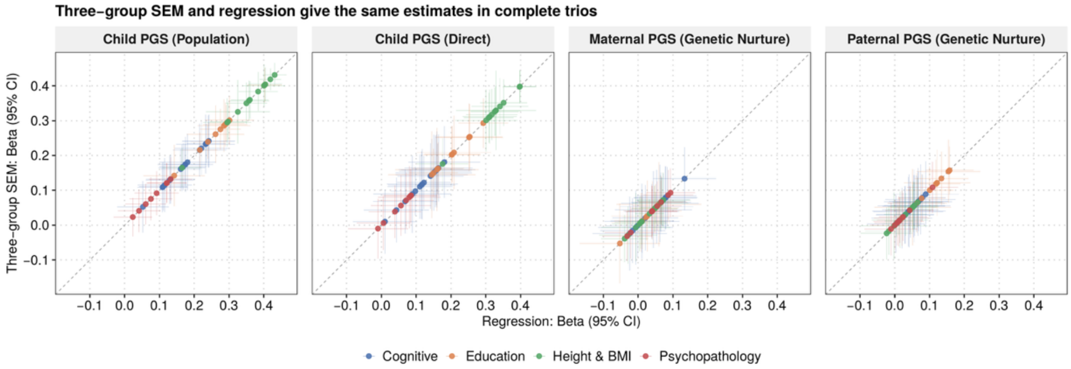

*Note*. Each point represents a trait at an age, with the estimate from the regression model reported in the main text on the x-axis and the corresponding estimate from the structural equation model on the y-axis. Both models are fitted to the same complete trios (N = 530 to 1356 individuals from 384 to 893 families). Panels show the four parameters: the child's polygenic score at the population level and as a direct genetic effect, and maternal and paternal genetic nurture. Colours denote trait domain. The dashed line is the identity line (y = x), on which the estimates from the two models would coincide exactly. Horizontal error bars give the 95% confidence interval from the regression and vertical error bars are from the structural equation model. More specifically, a cluster-robust sandwich estimator is used for the regression and robust maximum likelihood with clustering by family is used for the structural equation model. Corresponding estimates are presented in Supplementary Tables 2 (regression) and 4 (structural equation model). Discussed in Supplementary Note 2.

#### Supplementary Figure 8. Three-Group Structural Equation Model in the Whole Sample.

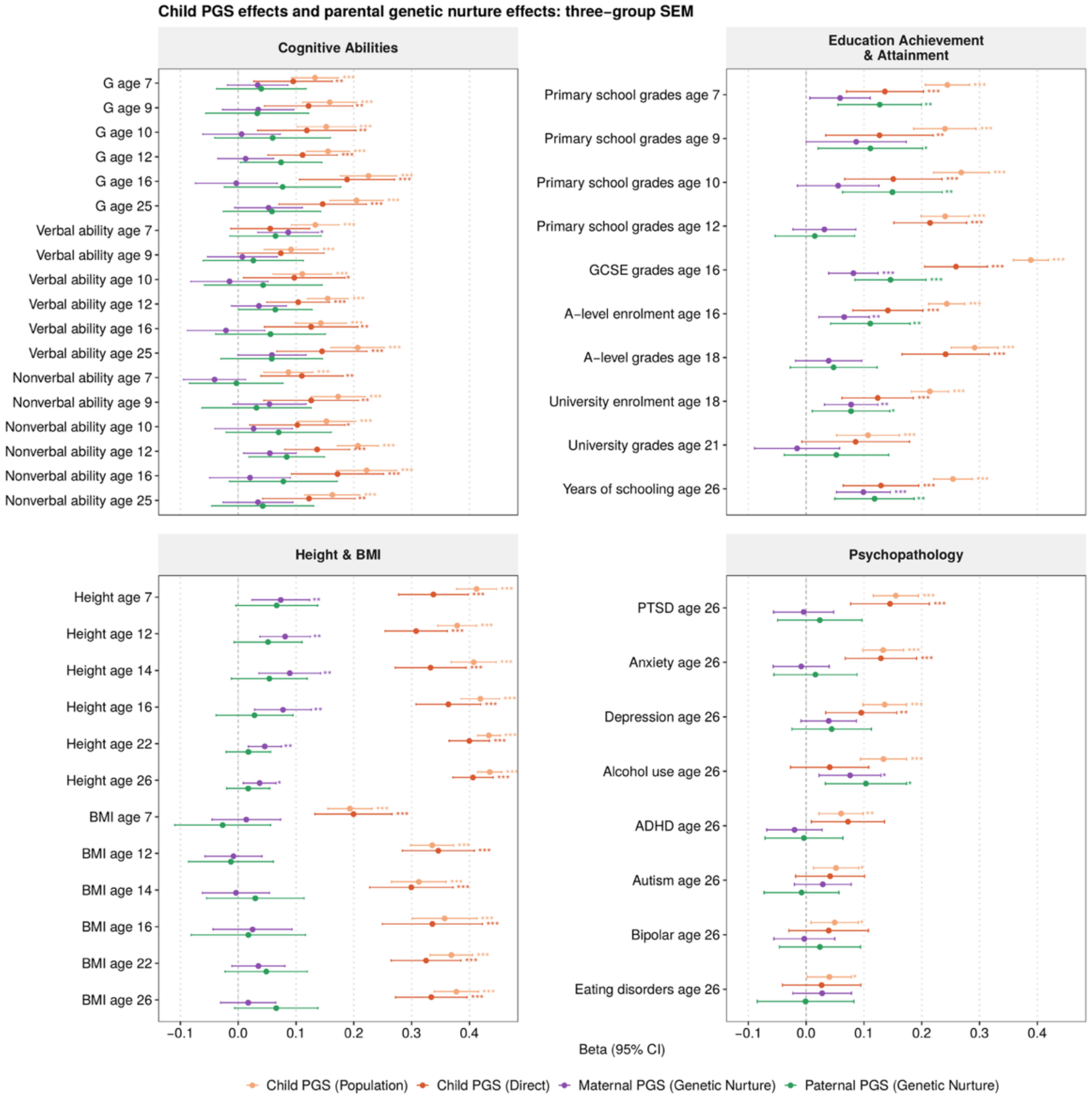

*Note.* Estimates are calculated from a three-group SEM in which complete trios, mother–child duos and father–child duos contribute jointly under full information maximum likelihood (N = 1328 to 3474 individuals, of whom 530 to 1356 are from complete trios), with child age, sex and genotyping chip as covariates. Regression coefficients, latent variances, and latent covariances are constrained to be equal across the three groups. The beta estimates are standardised and considered comparable to Pearson's r coefficient. Error bars represent 95% confidence intervals from a robust maximum likelihood estimator with clustering by family. Asterisks denote significance after false discovery rate correction (Benjamini–Hochberg) applied within each of four broad domains (cognitive abilities, educational achievement and attainment, height and BMI, and psychopathology). Point estimates, standard errors, 95% confidence intervals, and adjusted p-values are presented in Supplementary Table 4; model fits are presented in Supplementary Table 5. Discussed in Supplementary Note 2.

* FDR-adjusted p-value < 0.05; ** < 0.01; *** < 0.001

#### Supplementary Figure 9. Maternal Genetic Nurture Estimates: Sample-Composition Sensitivity for Three-Group SEM.

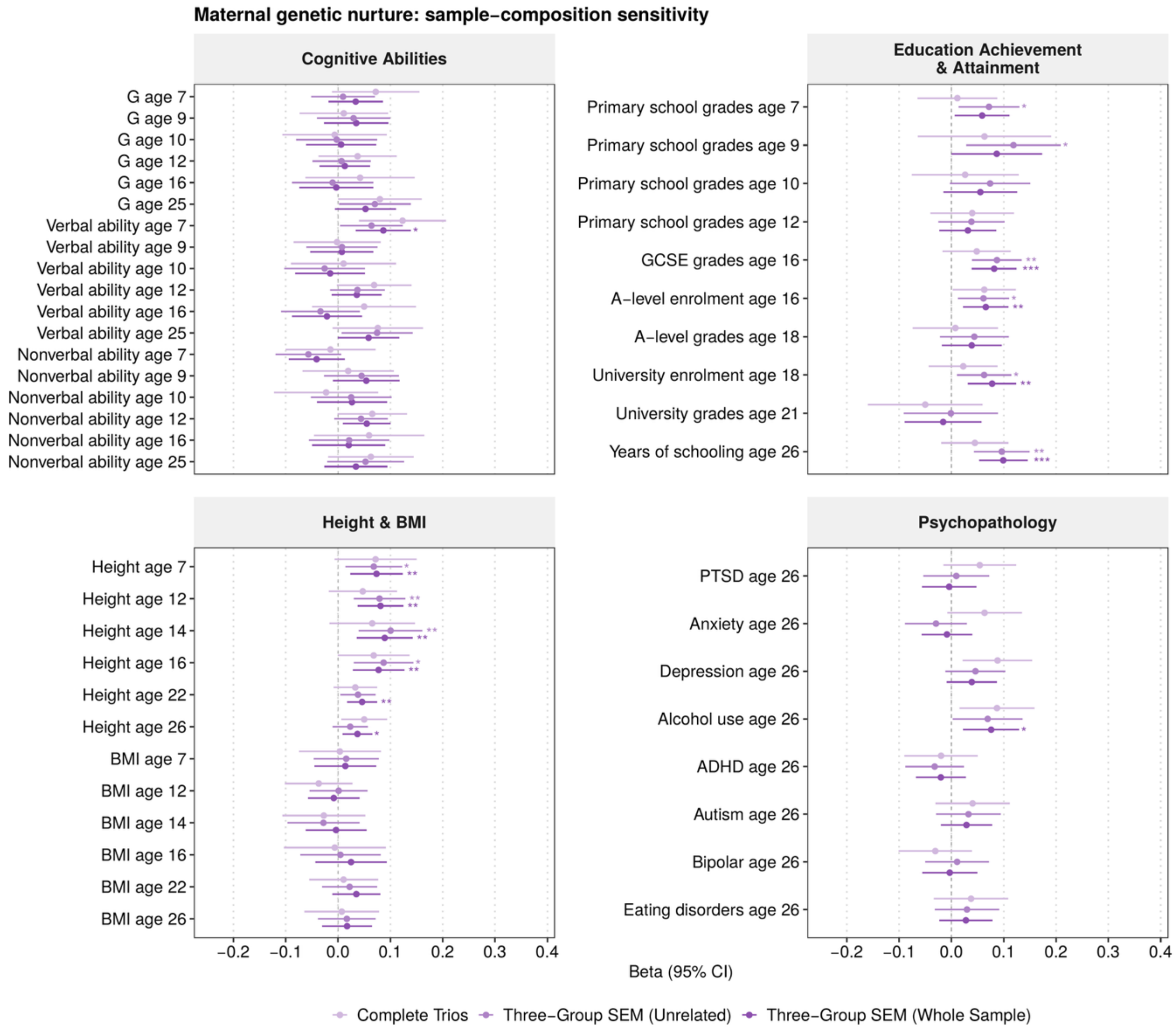

*Note*. Estimates are shown for three sample definitions: complete trios only (N = 530 to 1356 individuals), unrelated twins (N = 906 to 2321) and the whole sample (N = 1328 to 3474). All three are fitted with the same three-group structural equation model, with child age, sex and genotyping chip as covariates. For mothers, mean standard errors fell to 0.80 and then 0.69 of the complete-trio value as the sample widened. The beta estimates are standardised and considered comparable to Pearson’s r coefficient. Error bars represent 95% confidence intervals from a robust maximum likelihood estimator with clustering by family. Asterisks denote significance after false discovery rate correction (Benjamini–Hochberg) applied within each of the four broad domains and separately within each sample definition. Corresponding estimates are presented in Supplementary Table 4, with model fit in Supplementary Table 5. Discussed in Supplementary Note 2.

* FDR-adjusted p-value < 0.05; ** < 0.01; *** < 0.001

### Supplementary Figure 10. Paternal Genetic Nurture Estimates: Sample-Composition Sensitivity for Three-Group SEM.

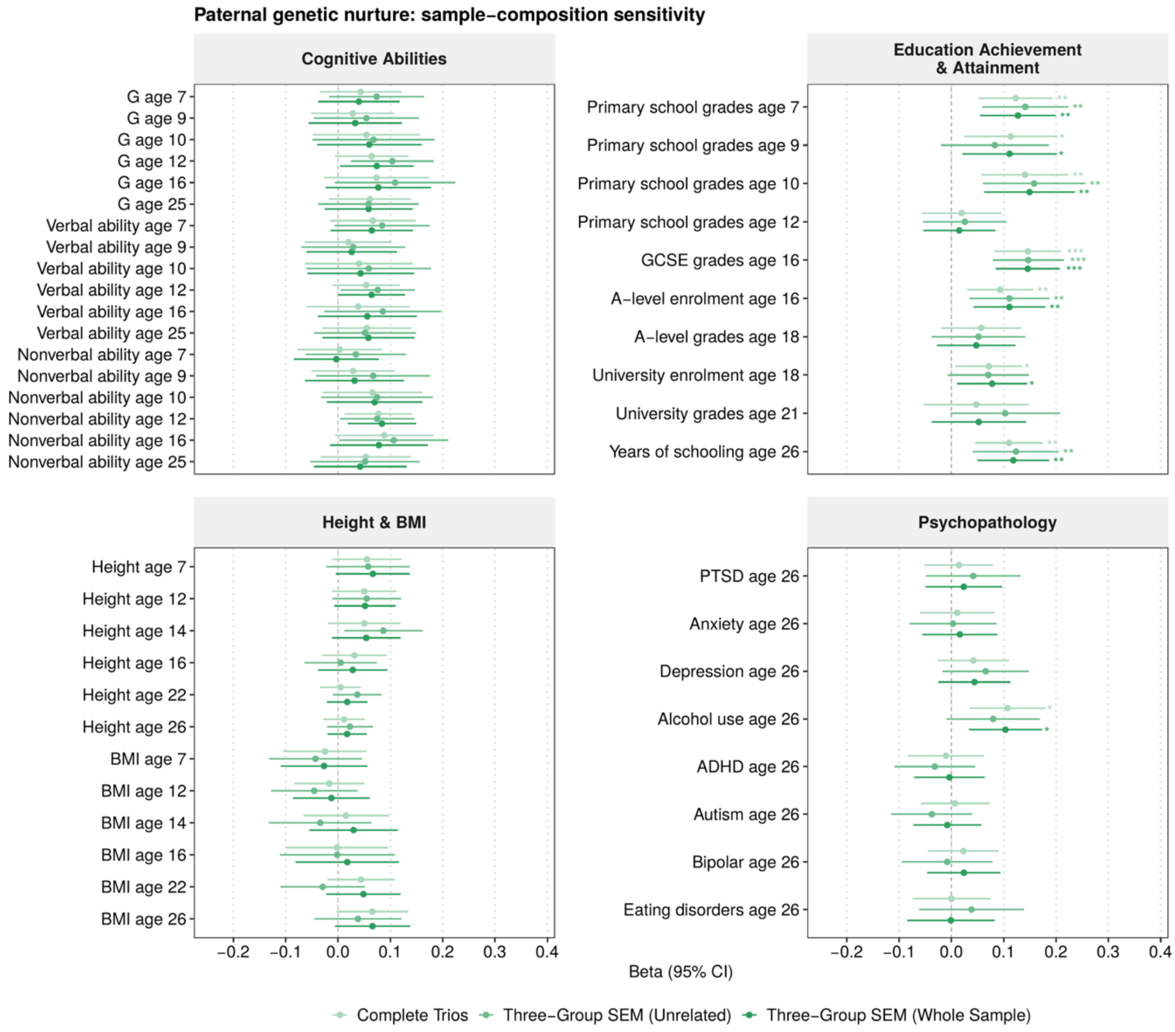

*Note*. Estimates are shown for three sample definitions: complete trios only (N = 530 to 1356 individuals), unrelated twins (N = 906 to 2321) and the whole sample (N = 1328 to 3474). All three are fitted with the same three-group structural equation model, with child age, sex and genotyping chip as covariates. Only 99 additional father–child duos were available, so the sample definitions differ little in the information they contribute for fathers. The beta estimates are standardised and considered comparable to Pearson’s r coefficient. Error bars represent 95% confidence intervals from a robust maximum likelihood estimator with clustering by family. Asterisks denote significance after false discovery rate correction (Benjamini–Hochberg) applied within each of the four broad domains and separately within each sample definition. Corresponding estimates are presented in Supplementary Table 4, with model fit in Supplementary Table 5. Discussed in Supplementary Note 2.
* FDR-adjusted p-value < 0.05; ** < 0.01; *** < 0.001

#### Supplementary Figure 11. Maternal Genetic Nurture Effects in Female versus Male Offspring: Three-Group SEM.

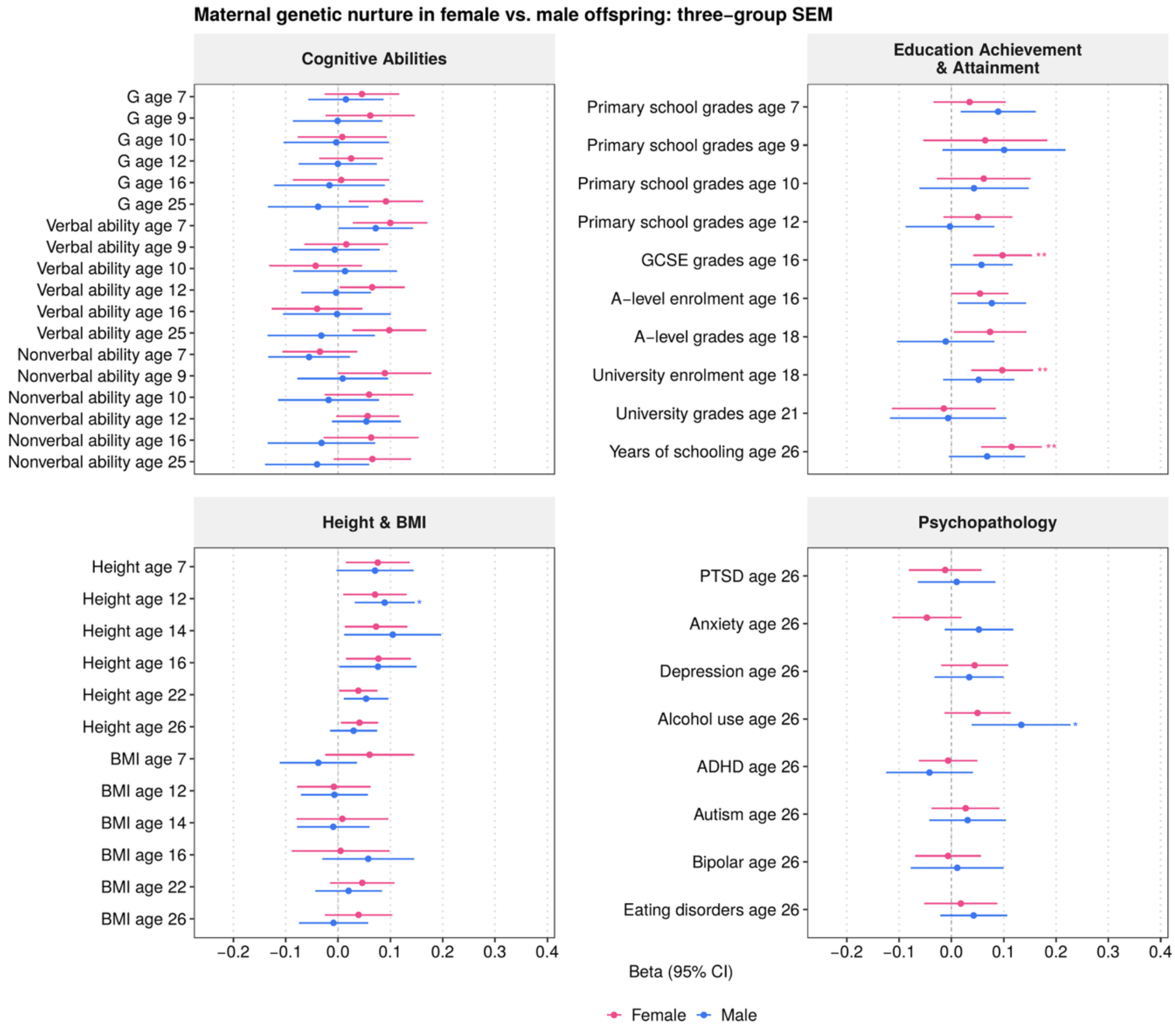

*Note*. The three-group structural equation model is fitted separately in female offspring (N = 767 to 1908 individuals, of whom 283 to 720 are from complete trios) and male offspring (N = 530 to 1566, of whom 235 to 636 are from complete trios), with child age and genotyping chip as covariates. The beta estimates are standardised and considered comparable to Pearson's r coefficient. Error bars represent 95% confidence intervals from a robust maximum likelihood estimator with clustering by family. Asterisks denote significance after false discovery rate correction (Benjamini–Hochberg) applied within each of the four broad domains and separately within each subgroup. Sex differences were tested for each trait using an independent-samples z test on the coefficients with the same multiple testing correction; none reached significance. Corresponding estimates are presented in Supplementary Table 4, with model fit in Supplementary Table 5. Discussed in Supplementary Note 2.

* FDR-adjusted p-value < 0.05; ** < 0.01; *** < 0.001

### Supplementary Figure 12. Paternal Genetic Nurture Effects in Female versus Male Offspring: Three-Group SEM.

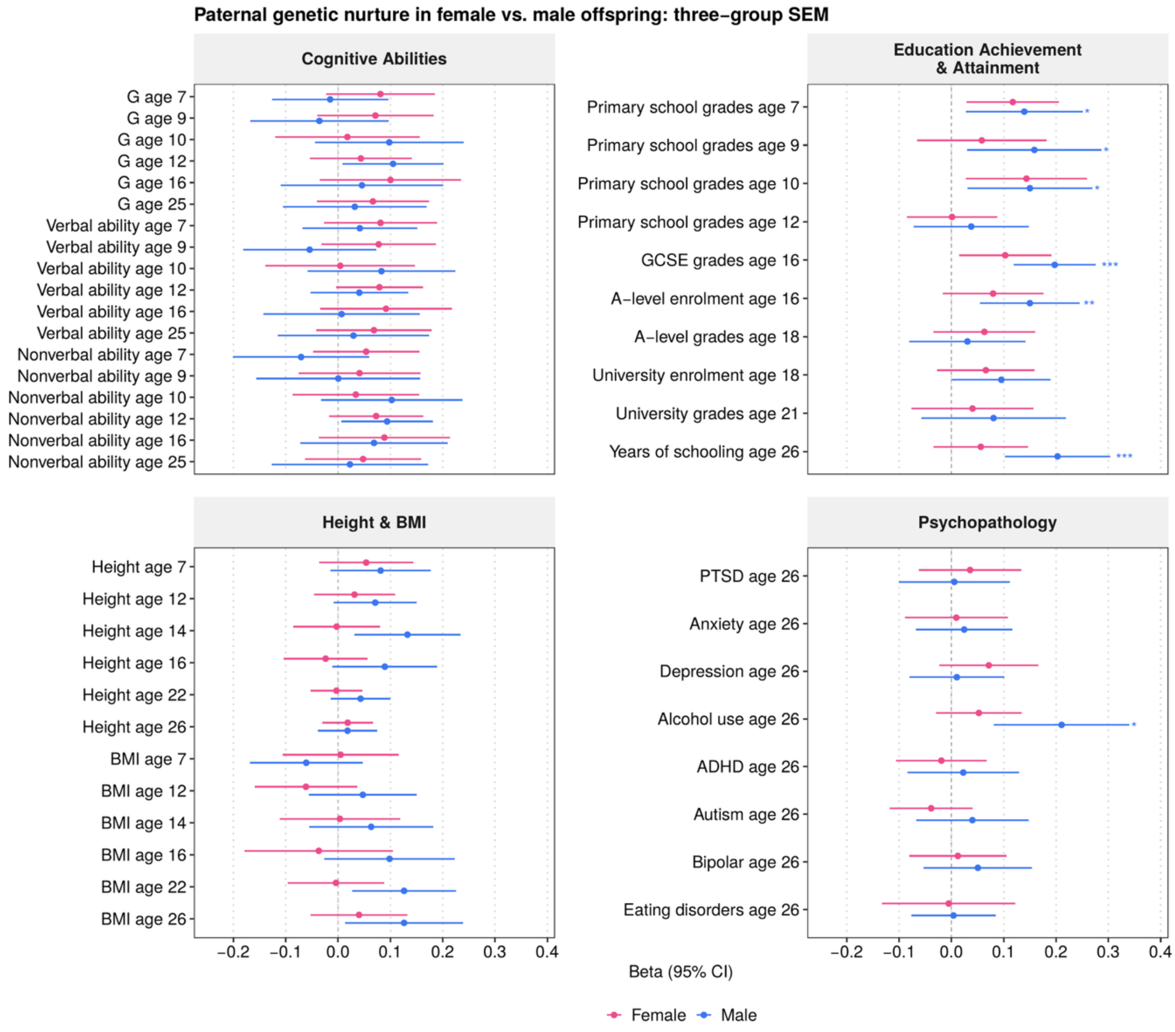

*Note*. The three-group structural equation model is fitted separately in female offspring (N = 767 to 1908 individuals, of whom 283 to 720 are from complete trios) and male offspring (N = 530 to 1566, of whom 235 to 636 are from complete trios), with child age and genotyping chip as covariates. The beta estimates are standardised and considered comparable to Pearson's r coefficient. Error bars represent 95% confidence intervals from a robust maximum likelihood estimator with clustering by family. Because fewer fathers than mothers were genotyped, paternal standard errors were larger than the corresponding maternal ones for every trait. Asterisks denote significance after false discovery rate correction (Benjamini–Hochberg) applied within each of the four broad domains and separately within each subgroup. Sex differences were tested for each trait using an independent-samples z test on the coefficients with the same multiple testing correction; none reached significance. Corresponding estimates are presented in Supplementary Table 4, with model fit in Supplementary Table 5. Discussed in Supplementary Note 2.
* FDR-adjusted p-value < 0.05; ** < 0.01; *** < 0.001

### Supplementary Figure 13. Sibling Indirect Genetic Estimates.

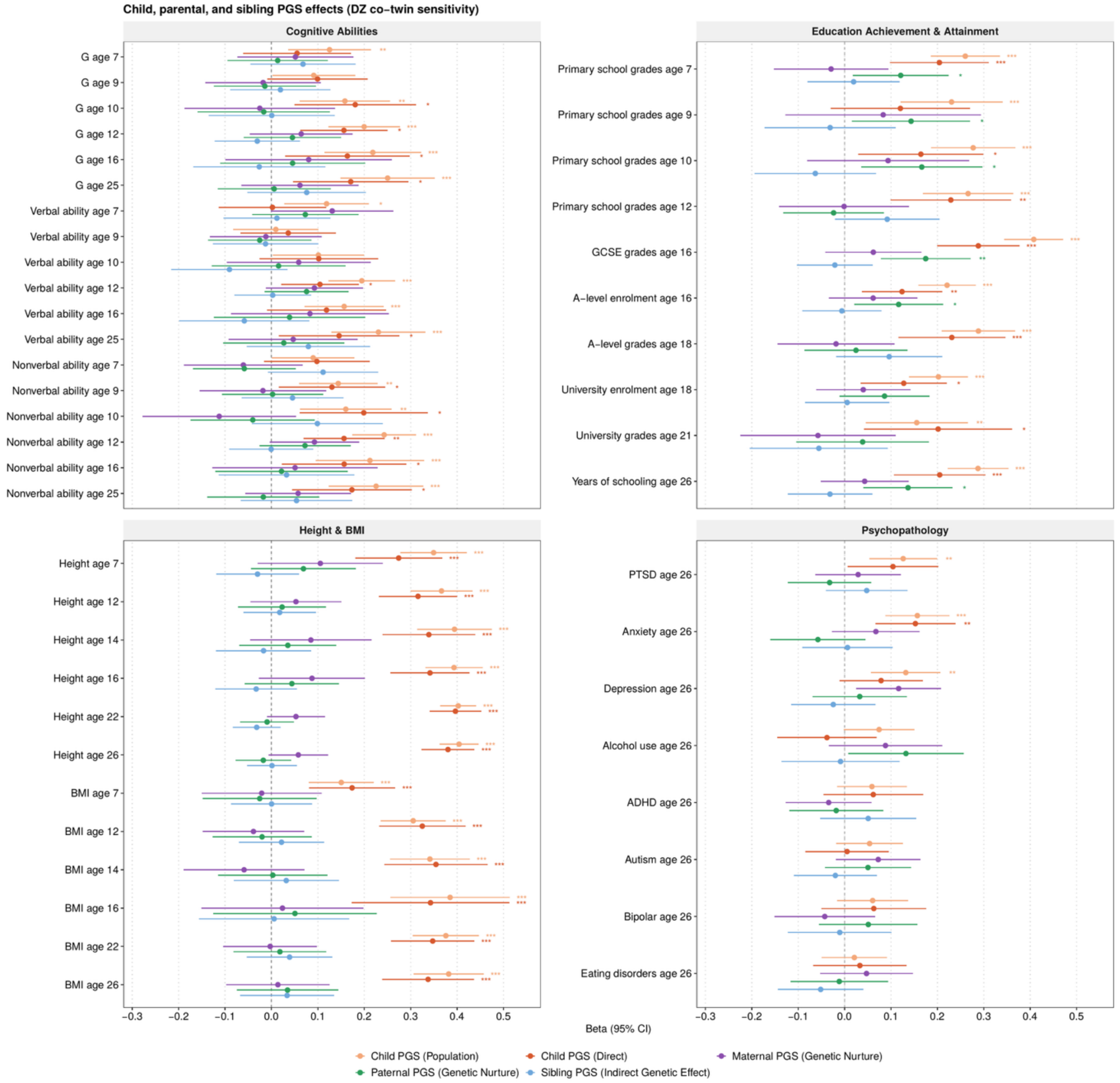

*Note*. Regressions are calculated in families in which both parents and both dizygotic twins were genotyped (N = 330 to 930 individuals from 183 to 466 families), adding the dizygotic co-twin's polygenic score to the trio model (quad model), with child age, sex and genotyping chip as covariates. Child polygenic score effects are shown at the population level (baseline) and as direct genetic effects; maternal and paternal effects index genetic nurture; the sibling coefficient indexes the indirect genetic effect of the dizygotic co-twin. The beta estimates are standardised and considered comparable to Pearson's r coefficient. Error bars represent 95% confidence intervals generated using a cluster-robust sandwich estimator accounting for relatedness within families. Asterisks denote significance after false discovery rate correction (Benjamini–Hochberg) applied within each of the four broad domains. Corresponding estimates are presented in Supplementary Table 2.

* FDR-adjusted p-value < 0.05; ** < 0.01; *** < 0.001

### Supplementary Figure 14. Sensitivity to Population Structure.

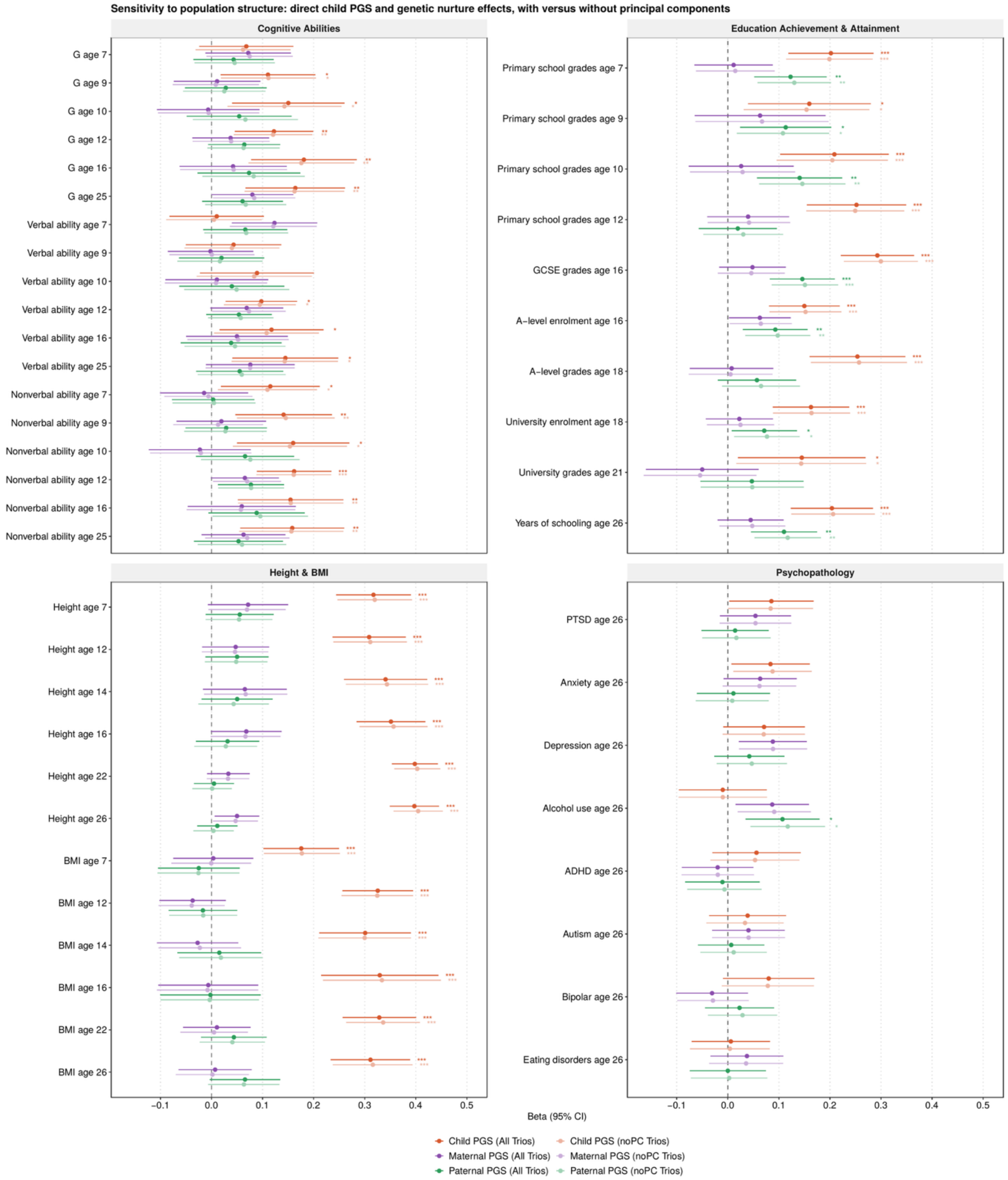

*Note*. Regressions are calculated using complete trios (N = 530 to 1356 individuals from 384 to 893 families), with child age, sex and genotyping chip as covariates in both models. The main models used polygenic scores pre-adjusted for the first ten genomic principal components (darker shades); the sensitivity models used unadjusted polygenic scores (lighter shades). The beta estimates are standardised and considered comparable to Pearson's r coefficient. Error bars represent 95% confidence intervals generated using a cluster-robust sandwich estimator accounting for relatedness within families. Asterisks denote significance after false discovery rate correction (Benjamini–Hochberg) applied within each of the four broad domains. Adjustment for principal components hardly changed the estimates, consistent with

TEDS being a genetically homogeneous UK cohort of predominantly white Europeans. Corresponding estimates are presented in Supplementary Table 2.
* FDR-adjusted p-value < 0.05; ** < 0.01; *** < 0.001

### Supplementary Figure 15. Sensitivity to Sex: Genetic Nurture Effects by Offspring Sex.

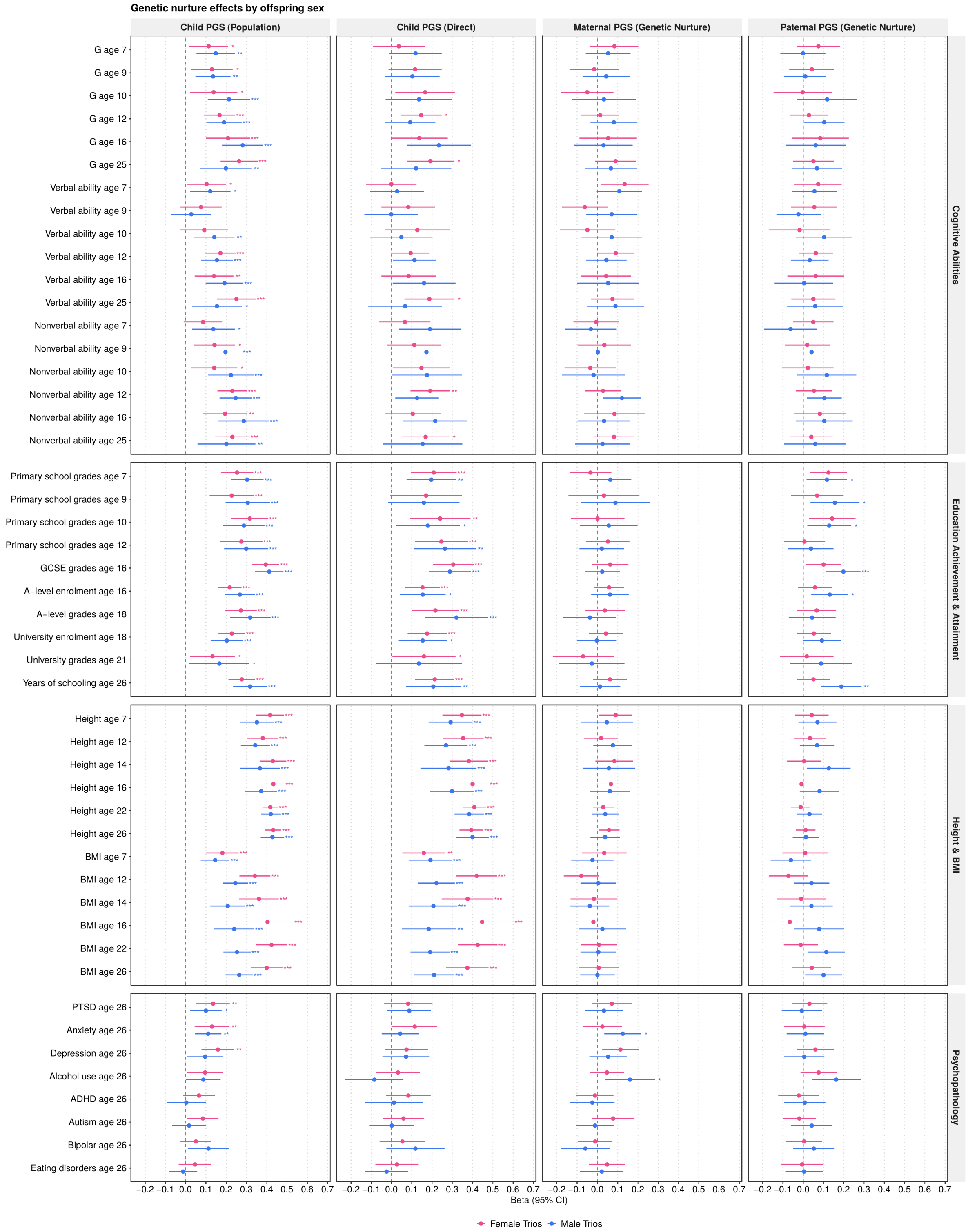

*Note*. Regressions are calculated separately in complete trios with female offspring (N = 283 to 720 individuals from 241 to 589 families) and male offspring (N = 235 to 636 individuals from 214 to 529 families), with child age and genotyping chip as covariates. The beta estimates are standardised and considered comparable to Pearson's r coefficient. Error bars represent 95% confidence intervals generated using a cluster-robust sandwich estimator accounting for relatedness within families.

Asterisks denote significance after false discovery rate correction (Benjamini–Hochberg) applied within each of the four broad domains and separately within each subsample. Because each subsample contains roughly half the analytic sample, standard errors are wider than in the main analysis and fewer estimates survive multiple testing correction. This reflects reduced precision rather than differences between the coefficients themselves. Corresponding estimates are presented in Supplementary Table 2.
* FDR-adjusted p-value < 0.05; ** < 0.01; *** < 0.001

#### Supplementary Figure 16. Sensitivity to Zygosity: Genetic Nurture Effects Among Monozygotic versus Dizygotic Twins.

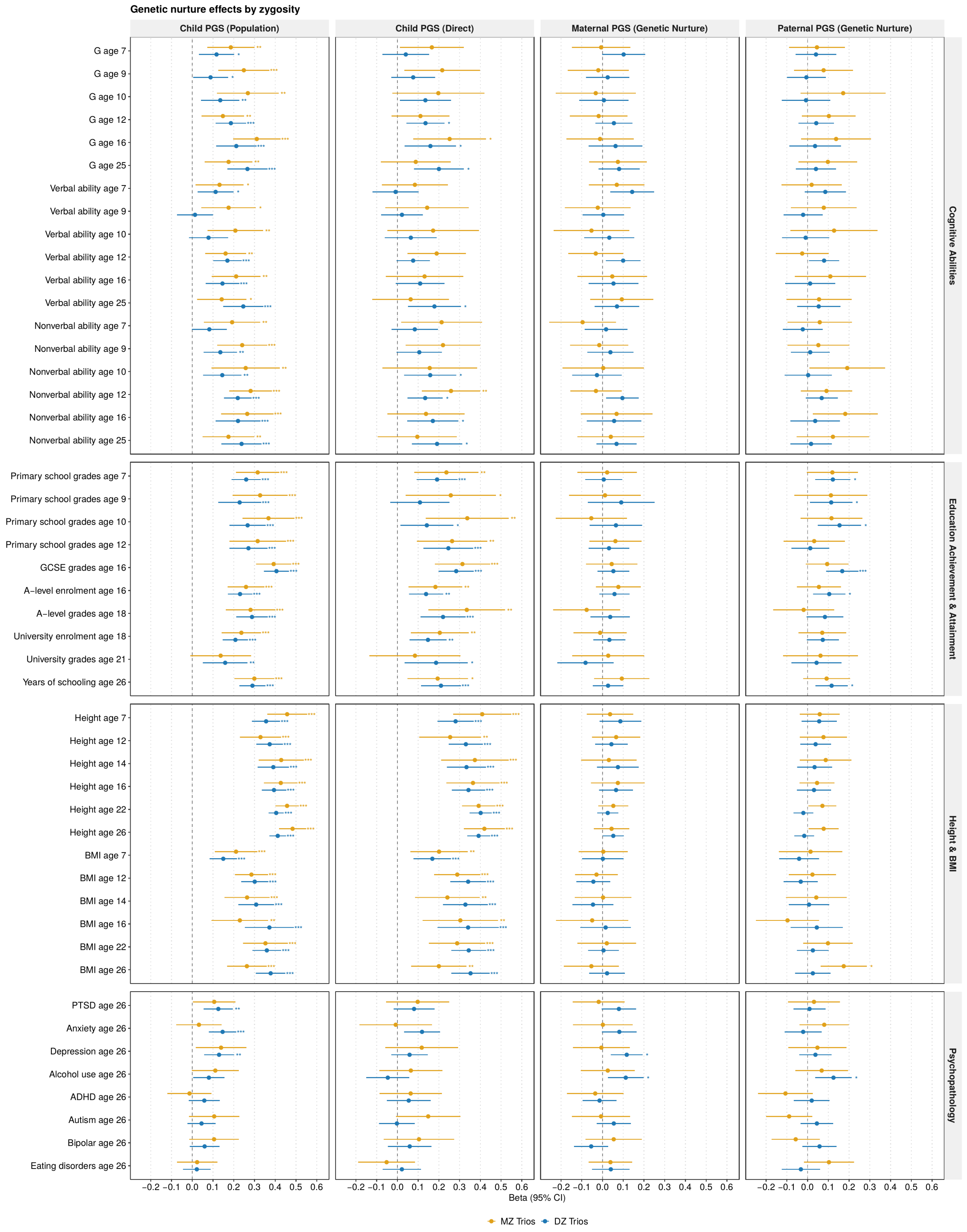

*Note*. Regressions are calculated separately in complete trios with monozygotic twins (N = 159 to 351 individuals from 159 to 351 families) and dizygotic twins (N = 368 to 1006 individuals from 224 to 542 families), with child age, sex and genotyping chip as covariates. For monozygotic pairs only one twin is retained, because co-twins share all polygenic scores and covariates. The beta estimates are standardised and considered comparable to Pearson's r coefficient. Error bars

represent 95% confidence intervals generated using a cluster-robust sandwich estimator accounting for relatedness within families. Asterisks denote significance after false discovery rate correction (Benjamini–Hochberg) applied within each of the four broad domains and separately within each subsample. The monozygotic subgroup is the smallest and its estimates are therefore less precise. Corresponding estimates are presented in Supplementary Table 2.
* FDR-adjusted p-value < 0.05; ** < 0.01; *** < 0.001

#### Supplementary Figure 17. Mediation of Genetic Nurture via Socio-Economic Status.

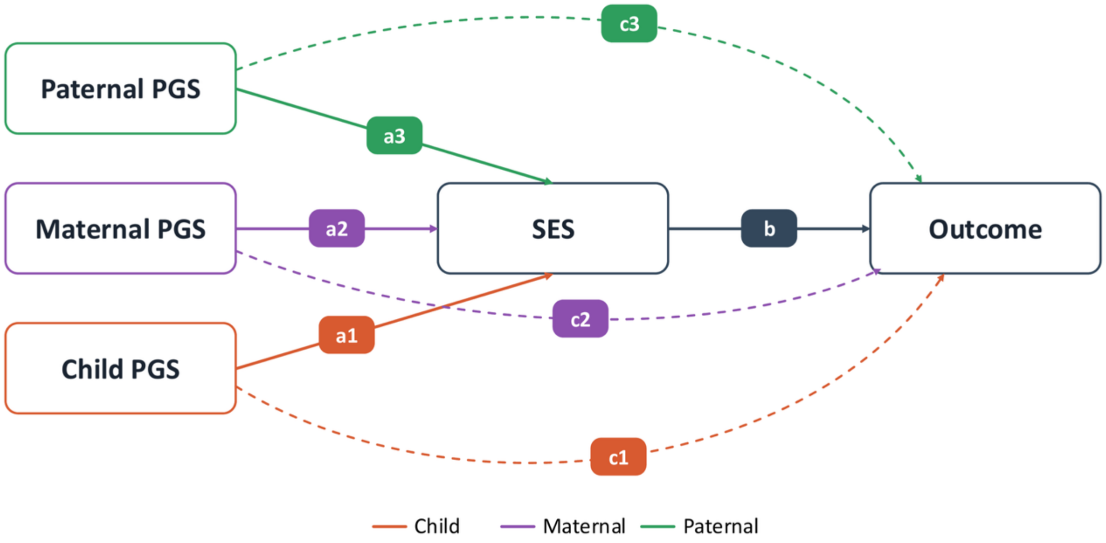

*Note.* Path diagram of the mediation model. The child's, maternal and paternal polygenic scores each predict family socio-economic status (paths $a_1$, $a_2$, $a_3$) and the outcome directly (paths $c_1$, $c_2$, $c_3$), with socio-economic status predicting the outcome (path b). Maternal and paternal indirect effects are $a_2 \times b$ and $a_3 \times b$. Total parental effects are the sum of the direct and indirect paths. All variables were residualised on child age, sex and genotyping chip before fitting, and models were estimated with cluster-robust standard errors accounting for relatedness within families. The diagram shows the model structure only. Estimates are presented in Supplementary Figures 18 and 19 and Supplementary Table 7. Discussed in Supplementary Note 3.

#### Supplementary Figure 18. Maternal Genetic Nurture Effects Mediated by Socio-Economic Status.

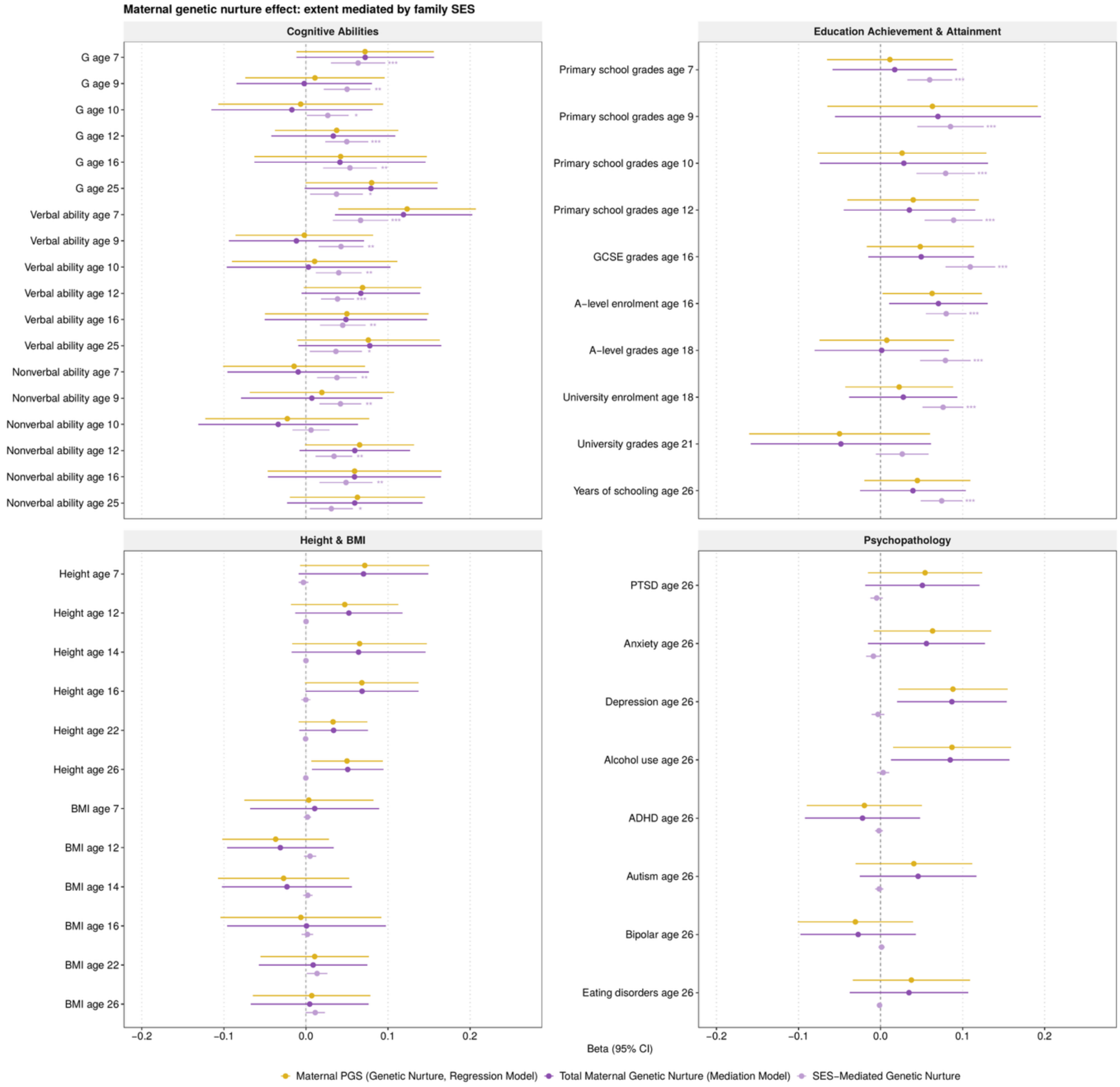

*Note*. Mediation model fitted in families with complete trios and non-missing socio-economic status at birth (N = 525 to 1338 individuals from 377 to 879 families), with child age, sex and genotyping chip residualised out beforehand. Three parameters are shown for each trait: the maternal genetic nurture estimate from the regression model (gold) shown in Supplementary Figure 2, the total maternal effect from the mediation model (dark purple), which is the sum of the direct and indirect paths, and the portion of that effect running through socio-economic status (light purple). The first two parameters are estimates of the same quantity from different models and are shown together as a consistency check. The beta estimates are standardised and considered comparable to Pearson's r coefficient. Error bars represent 95% confidence intervals calculated as the estimate ±1.96 standard errors, using cluster-robust standard errors accounting for relatedness within families. Asterisks denote significance after false discovery rate correction (Benjamini–Hochberg) applied within each of the four broad domains. Corresponding estimates are presented in Supplementary Table 7. Discussed in Supplementary Note 3.
* FDR-adjusted p-value < 0.05; ** < 0.01; *** < 0.001

#### Supplementary Figure 19. Paternal Genetic Nurture Effects Mediated by Socio-Economic Status.

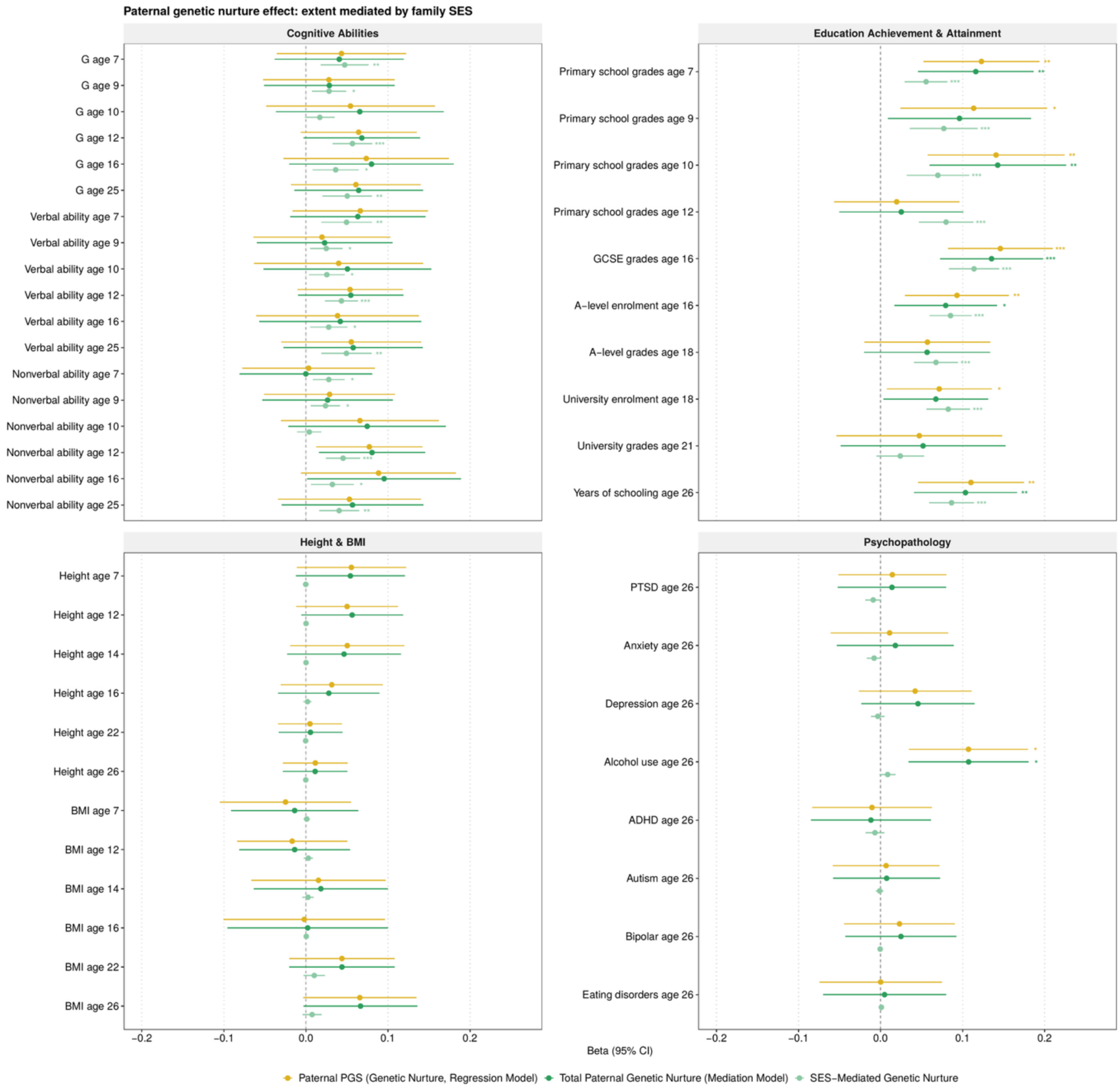

*Note*. Mediation model fitted in families with complete trios and non-missing socio-economic status at birth (N = 525 to 1338 individuals from 377 to 879 families), with child age, sex and genotyping chip residualised out beforehand. Three parameters are shown for each trait: the paternal genetic nurture estimate from the regression model (gold) shown in Supplementary Figure 2, the total paternal effect from the mediation model (dark green), which is the sum of the direct and indirect paths, and the portion of that effect running through socio-economic status (light green). The first two parameters are estimates of the same quantity from different models and are shown together as a consistency check. The beta estimates are standardised and considered comparable to Pearson's r coefficient. Error bars represent 95% confidence intervals calculated as the estimate ±1.96 standard errors, using cluster-robust standard errors accounting for relatedness within families. Asterisks denote significance after false discovery rate correction (Benjamini–Hochberg) applied within each of the four broad domains. Corresponding estimates are presented in Supplementary Table 7. Discussed in Supplementary Note 3.
* FDR-adjusted p-value < 0.05; ** < 0.01; *** < 0.001

#### Supplementary Figure 20. Twin ACE Estimates for All Outcomes in Families with Parental DNA.

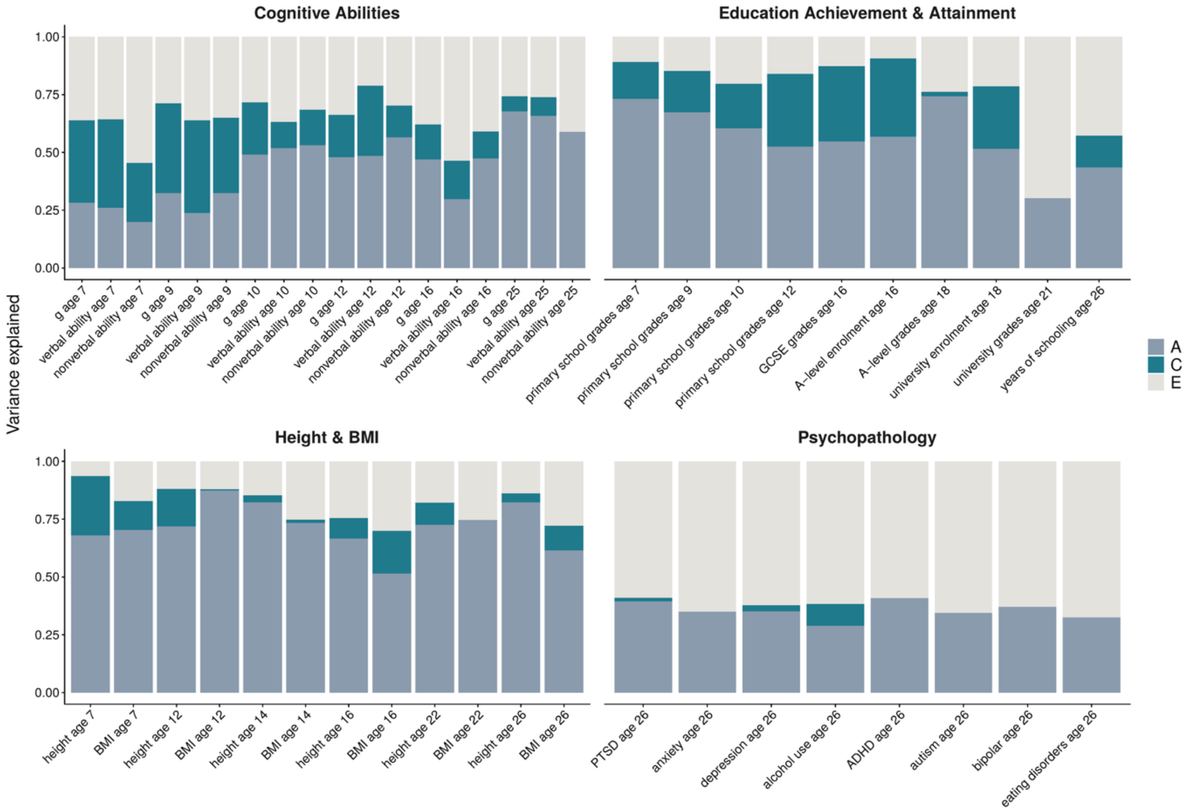

*Note.* Bars show the proportion of phenotypic variance attributable to additive genetic (A), shared environmental (C), and nonshared environmental (E) influences, which sum to one. Estimates are from univariate twin models fitted in TEDS families in which the offspring were genotyped and at least one parent provided DNA (151 to 476 monozygotic and 295 to 1259 dizygotic twin pairs). This subsample comprises both duo and trio families and is therefore broader than the complete-trio sample used for the genetic nurture analyses. For the 46 continuous measures, phenotypes were residualised on age and sex and standardised before modelling. A-level enrolment at age 16 and university enrolment at age 18 are binary and were modelled using a liability threshold model with age and sex as covariates. These two estimates refer to the latent liability rather than the observed dichotomy. Although different models were used, the estimates are essentially comparable. Estimates averaged within trait domains are shown in Figure 4. Sample size for each trait, point estimates, and 95% confidence intervals are presented in Supplementary Table 8.

### Supplementary Figure 21. Twin ACE Estimates for the Whole Sample.

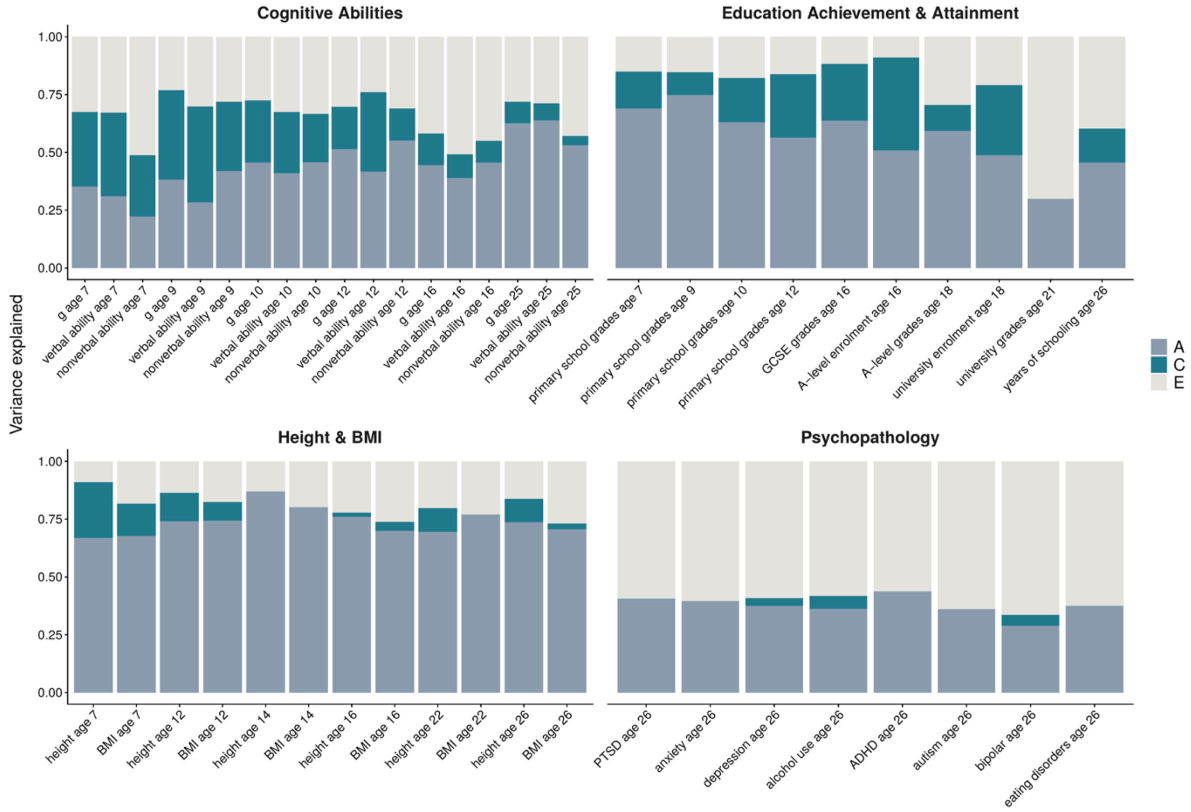

*Note*. Bars show the proportion of phenotypic variance attributable to additive genetic (A), shared environmental (C), and nonshared environmental (E) influences, which sum to one. Estimates are from univariate twin models fitted in the whole TEDS twin sample regardless of whether parental DNA was available (533 to 2391 monozygotic and 671 to 4319 dizygotic twin pairs), serving as a robustness check on Figure 4 and Supplementary Figure 20. Model specification is identical to Figure 4: the 46 continuous measures were residualised on age and sex and standardised before modelling, and the two binary measures were modelled using a liability threshold model with age and sex as covariates. Although different models were used, the estimates are essentially comparable. Estimates averaged within trait domains are shown in Supplementary Figure 22. Sample size for each trait, point estimates, and 95% confidence intervals are presented in Supplementary Table 9.

### Supplementary Figure 22. Twin ACE Estimates for the Whole Sample, Averaged Across Ages.

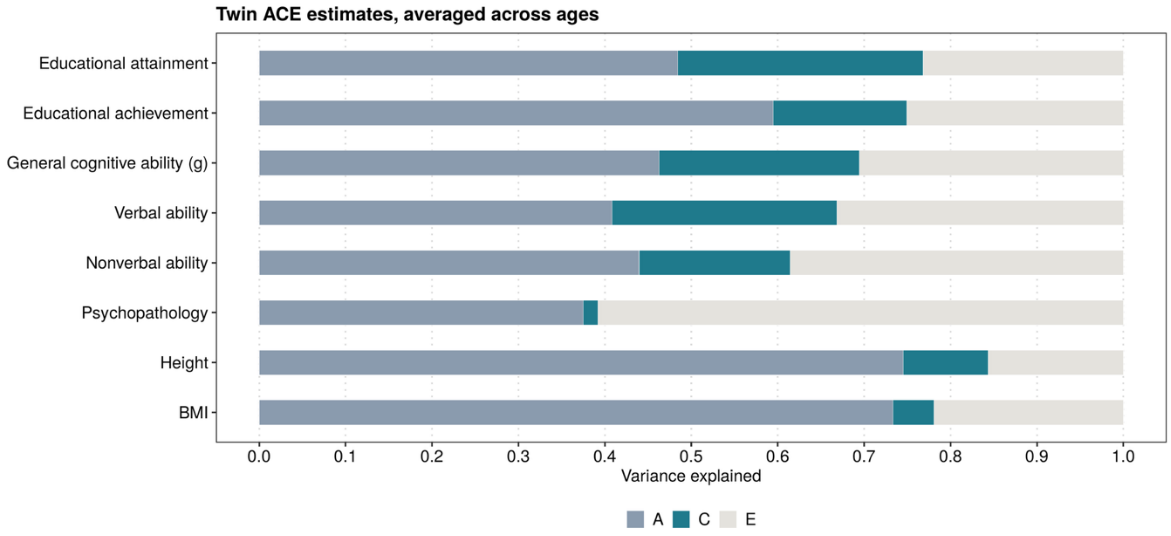

*Note.* Bars show the proportion of phenotypic variance attributable to additive genetic (A), shared environmental (C) and nonshared environmental (E) influences, averaged across ages within each of the eight trait domains. Estimates are from univariate twin models fitted in the whole TEDS twin sample regardless of whether parental DNA was available (533 to 2391 monozygotic and 671 to 4319 dizygotic twin pairs), as a sensitivity check for Figure 4. Estimates for all 48 outcomes are shown in Supplementary Figure 21 and presented in Supplementary Table 9.

#### Supplementary Figure 23. Maternal and Paternal Genetic Nurture within the Population Prediction and Shared Environment.

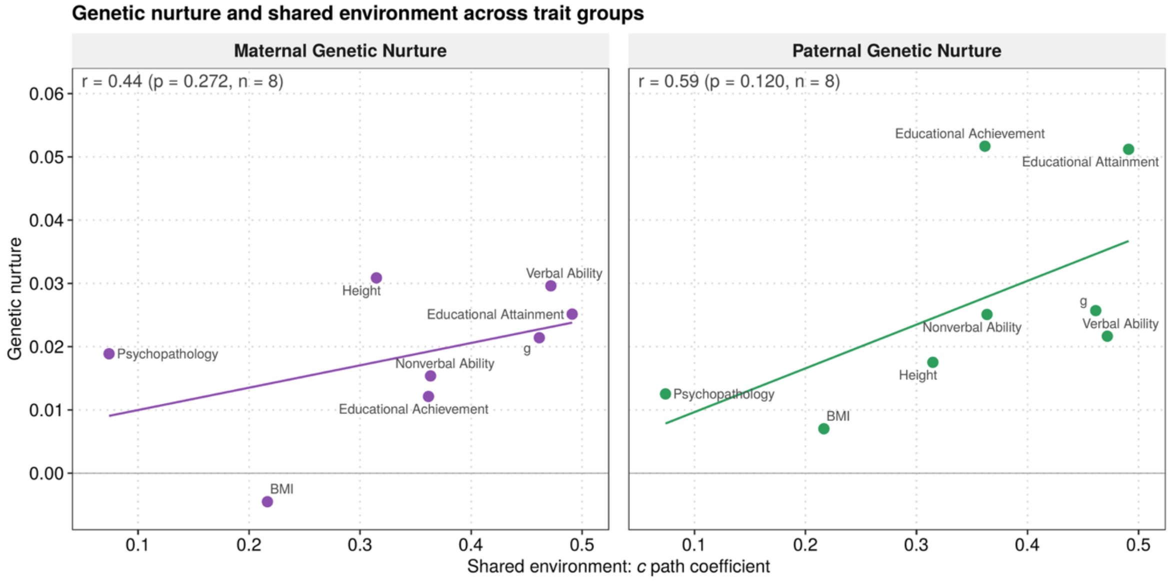

*Note.* As Figure 5, with maternal and paternal genetic nurture shown in separate panels. Each point represents one of eight trait domains. The x-axis shows the shared environmental path coefficient, calculated as the square root of $c^2$ (Figure 4) and averaged within each domain. The y-axis shows maternal (left) or paternal (right) genetic nurture within the population PGS prediction, that is, the parental PGS coefficient multiplied by the corresponding parent–child PGS correlation (Figure 3), averaged within each domain. Both axes are on a standard deviation scale. The lines show the least-squares regression across the eight domains (maternal: $r = 0.44$, $p = 0.27$; paternal: $r = 0.59$, $p = 0.12$; $n = 8$). The underlying estimates are presented in Supplementary Tables 3 (genetic nurture) and 8 (shared environment).

#### Supplementary Figure 24. Maternal and Paternal Genetic Nurture as a Proportion of the Population Prediction and Shared Environment.

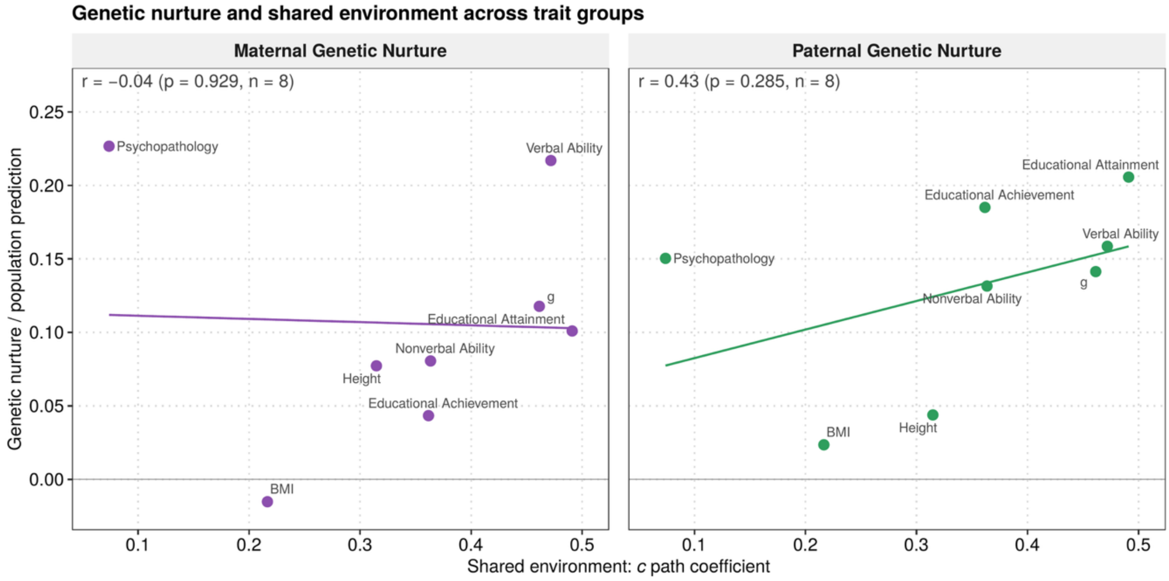

*Note.* As Figure 6, with maternal and paternal genetic nurture shown in separate panels. Each point represents one of eight trait domains. The x-axis shows the shared environmental path coefficient, calculated as the square root of $c^2$ (Figure 4) and averaged within each domain, as in Supplementary Figure 23. The y-axis shows maternal (left) or paternal (right) genetic nurture within the population PGS prediction divided by the population PGS prediction (Figure 3). Estimates were averaged within each domain before the ratio was calculated. The lines show the least-squares regression across the eight domains (maternal: $r = -0.04$, $p = 0.93$; paternal: $r = 0.43$, $p = 0.29$; $n = 8$). The underlying estimates are presented in Supplementary Tables 3 (genetic nurture) and 8 (shared environment).

### Supplementary Figure 25. Incremental Variance Explained by Genetic Nurture from Duos versus Trios.

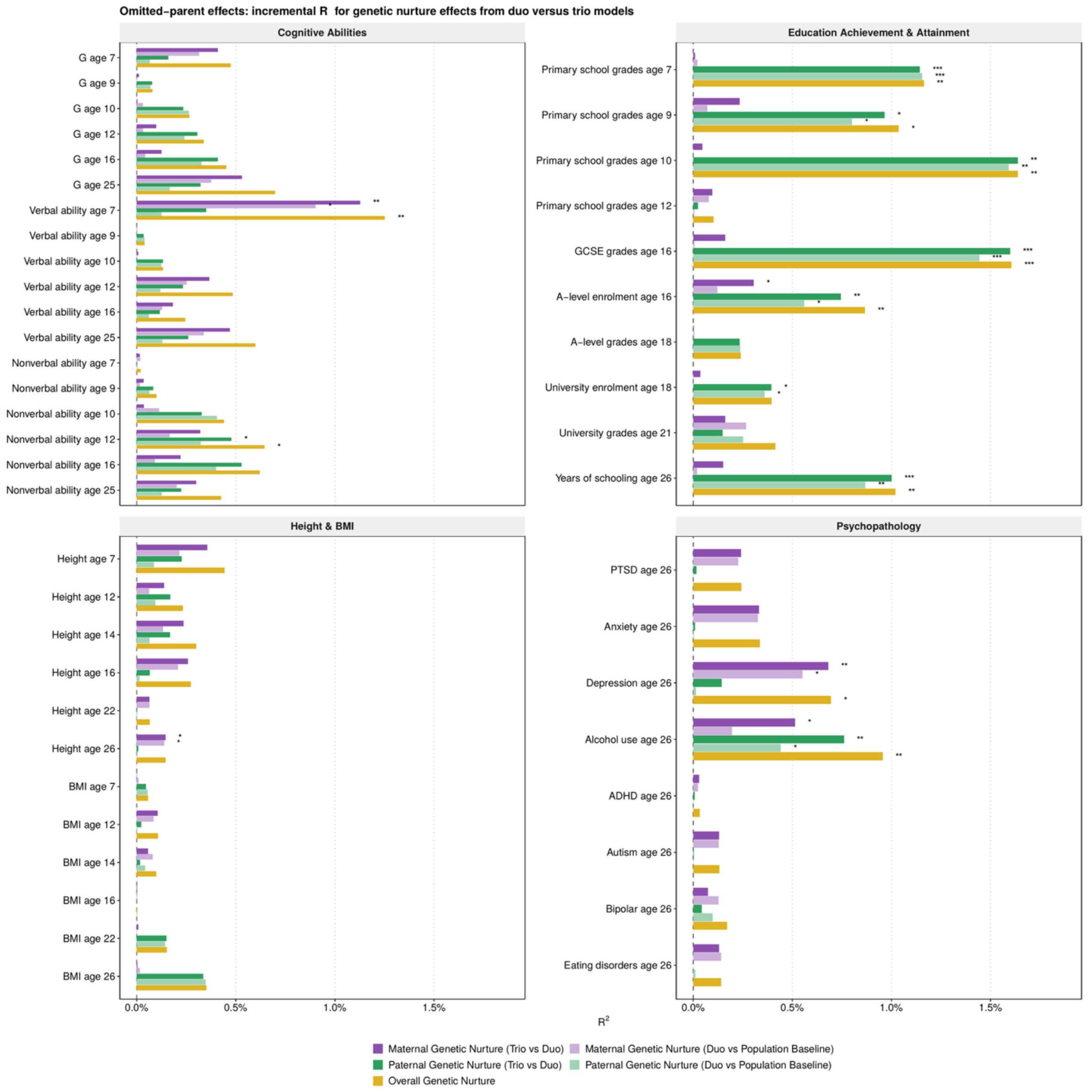

*Note.* Incremental variance explained (ΔR²) across three nested models: the population model (child PGS only), the duo models (child PGS and one parent's PGS) and the trio model (child PGS and both parents' PGS). For maternal genetic nurture, duo versus population baseline shows how much adding the mother's PGS to a model with the child's PGS improves the prediction of the child's outcome. One potential issue with this comparison is that the mother's PGS also partly indexes paternal genetic nurture through the correlation between the parents' PGS (assortative mating). We therefore also compared the trio model with the child–father duo model, which shows how much the mother's PGS adds beyond both the child's and the father's PGS. The same comparisons were made for paternal genetic nurture. Overall genetic nurture compares the trio model with the population baseline. Asterisks denote whether the two nested models differ significantly (cluster-robust Wald tests). Corresponding estimates are presented in Supplementary Table 2. Discussed in Supplementary Note 1.

#### Supplementary Figure 26. PGS–SES Correlation.

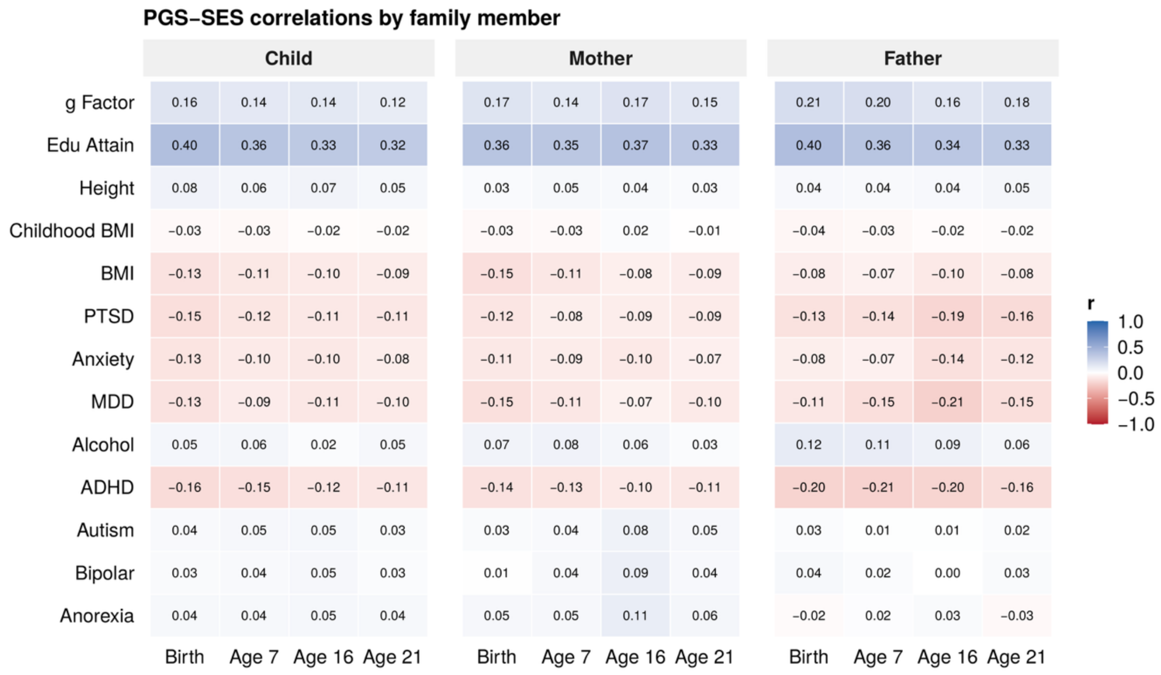

*Note*. Pearson correlations between each polygenic score and family socio-economic status at four time points (birth and ages 7, 16 and 21), shown separately for the child, the mother and the father. Correlations were calculated in the unrelated subsample (one twin per family) using pairwise complete observations on the polygenic scores after residualisation on the first ten genomic principal components. Correlations were of similar magnitude for parents and children, and were stable across the four socio-economic status measures. Corresponding estimates are presented in Supplementary Table 1. Discussed in Supplementary Note 3.

#### Supplementary Figure 27. Phenotype–SES Correlation.

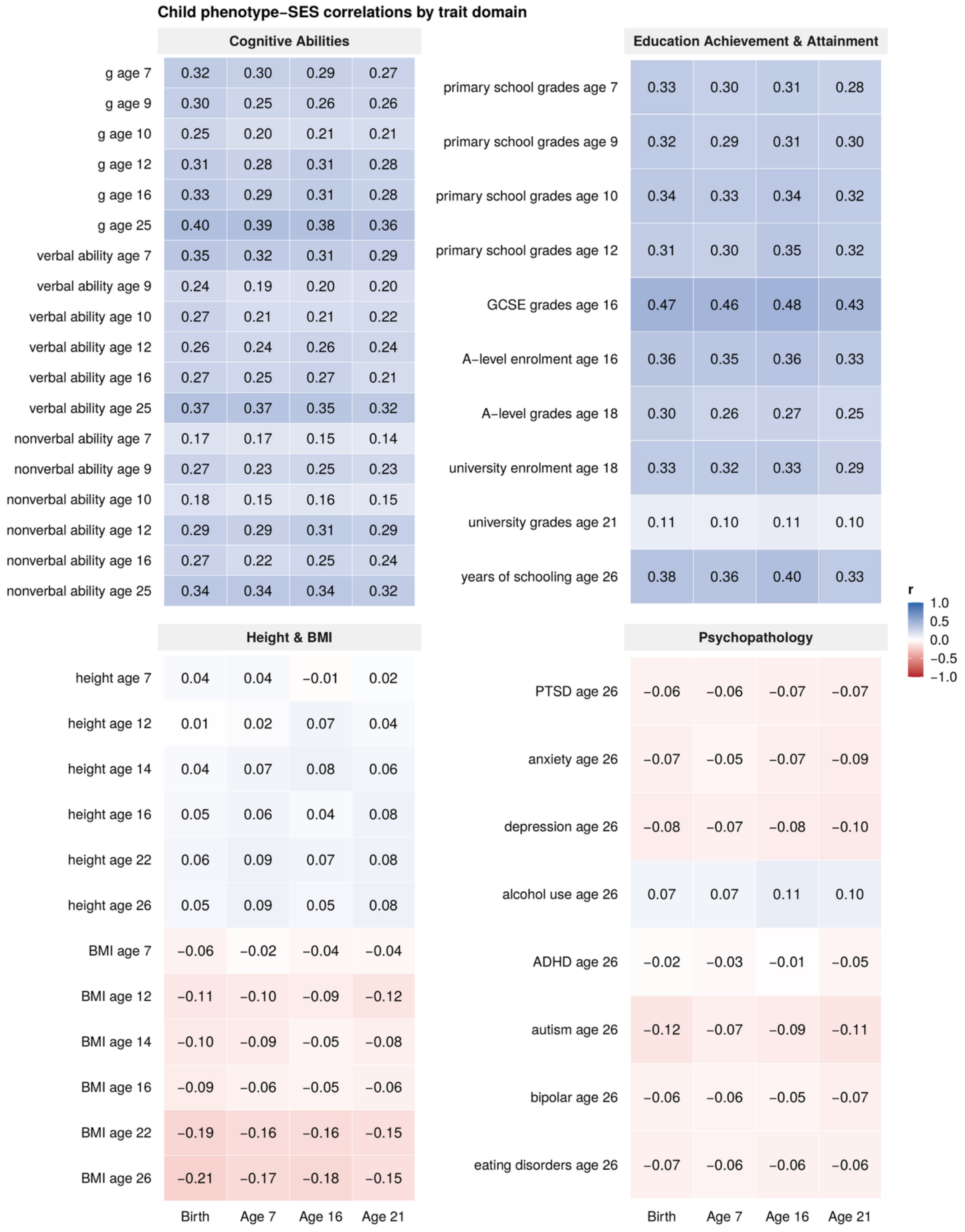

*Note*. Pearson correlations between each child phenotype and family socio-economic status at four time points (birth and ages 7, 16 and 21). Correlations were calculated in the unrelated subsample (one twin per family) using pairwise complete observations on raw phenotypes. Associations with socio-economic status were substantial for educational outcomes ($r = 0.10$ to $0.48$) and cognitive abilities ($r = 0.14$ to $0.40$), and close to zero for height ($r = -0.01$ to $0.09$) and psychopathology ($r = -0.12$ to $0.11$), with BMI intermediate and negative in adulthood ($r = -0.21$ to

−0.02). Corresponding estimates are presented in Supplementary Table 1. Discussed in Supplementary Note 3.

### Supplementary Tables

All supplementary tables are presented in a separate spreadsheet (GN_Supplementary_Tables.xlsx). Here, we present a table of contents.

**Supplementary Table 1.** Correlations among polygenic scores, phenotypes, principal components and family socio-economic status.
**Supplementary Table 2.** Regression estimates of child, maternal and paternal polygenic score effects.
**Supplementary Table 3.** Population polygenic score prediction expressed as direct and genetic nurture effects.
**Supplementary Table 4.** Three-group structural equation model estimates.
**Supplementary Table 5.** Model fit of the three-group structural equation models.
**Supplementary Table 6.** Summary of maternal and paternal genetic nurture estimates across models and samples.
**Supplementary Table 7.** Mediation of genetic nurture by family socio-economic status.
**Supplementary Table 8.** Twin ACE estimates in families with parental genotype data.
**Supplementary Table 9.** Twin ACE estimates in the whole twin sample.
**Supplementary Table 10.** Sample characteristics.
**Supplementary Table 11.** Comparison of phenotypes, SES and PGS between genotyped children with and without genotyped parents.
**Supplementary Table 12.** Comparison of phenotypes and SES between children with and without their own genotype data, among families without genotyped parents.
**Supplementary Table 13.** Descriptive statistics in complete trios.
**Supplementary Table 14.** Descriptive statistics in all families with at least one genotyped parent.
**Supplementary Table 15.** Regressions of phenotypes and family socio-economic status on covariates.
**Supplementary Table 16.** Regressions of polygenic scores on covariates.
